# Surprise: a unified theory and experimental predictions

**DOI:** 10.1101/2021.11.01.466796

**Authors:** Alireza Modirshanechi, Johanni Brea, Wulfram Gerstner

**Affiliations:** EPFL, School of Computer and Communication Sciences and School of Life Sciences, Lausanne, Switzerland

## Abstract

Surprising events trigger measurable brain activity and influence human behavior by affecting learning, memory, and decision-making. Currently there is, however, no consensus on the definition of surprise. Here we identify 16 mathematical definitions of surprise in a unifying framework, show how these definitions relate to each other, and prove under what conditions they are indistinguishable. We classify these surprise measures into four main categories: (i) change-point detection surprise, (ii) information gain surprise, (iii) prediction surprise, and (iv) confidence-correction surprise. We design experimental paradigms where different categories make different predictions: we show that surprise-modulation of the speed of learning leads to sensible adaptive behavior only for change-point detection surprise whereas surprise-seeking leads to sensible exploration strategies only for information gain surprise. However, since neither change-point detection surprise nor information gain surprise perfectly reflect the definition of ‘surprise’ in natural language, a combination of prediction surprise and confidence-correction surprise is needed to capture intuitive aspects of surprise perception. We formalize this combination in a new definition of surprise with testable experimental predictions. We conclude that there cannot be a single surprise measure with all functions and properties previously attributed to surprise. Consequently, we postulate that multiple neural mechanisms exist to detect and signal different aspects of surprise.

**Author note:** AM is grateful to Vasiliki Liakoni, Martin Barry, and Valentin Schmutz for many useful discussions in the course of the last few years, and to Andrew Barto for insightful discussions through and after EPFL Neuro Symposium 2021 on “Surprise, Curiosity and Reward: from Neuroscience to AI”. We thank K. Robbins and collaborators for their publicly available experimental data (Robbins et al., 2018). All code needed to reproduce the results reported here will be made publicly available after publication acceptance. This research was supported by Swiss National Science Foundation (no. 200020_184615). Correspondence concerning this article should be addressed to Alireza Modirshanechi, School of Computer and Communication Sciences and School of Life Sciences, EPFL, Lausanne, Switzerland.

## Introduction

Imagine you open the curtains one morning and find the street in front of your apartment covered by fresh snow. If you have expected a warm and sunny morning according to the weather forecast, you feel ‘surprised’ as you see the white streets; as a consequence of surprise, the activity of many neurons in your brain changes (Antony et al., 2021; Kolossa et al., 2015; Mars et al., 2008; Modirshanechi et al., 2019; Ostwald et al., 2012; Squires et al., 1976) and your pupils dilate (Antony et al., 2021; Nassar et al., 2012; Preuschoff et al., 2011). Surprise affects how we predict and perceive our future and how we remember our past (Antony et al., 2021; Dubey & Griffiths, 2020; Gershman et al., 2017; Gottlieb & Oudeyer, 2018; Oudeyer & Kaplan, 2009; Sinclair & Barense, 2018; Soltani & Izquierdo, 2019; Yu & Dayan, 2005). For example, some studies suggest that you would rely less on the weather forecast for your future plans after the snowy morning (Behrens et al., 2007; Findling et al., 2021; Gerstner et al., 2018; Nassar et al., 2010; Soltani & Izquierdo, 2019; Xu et al., 2021; Yu & Dayan, 2005). Other studies predict that you would remember more vividly the face of the random stranger who walked past the street in that very moment you felt surprised (Rouhani & Niv, 2021; Rouhani et al., 2018), and some predict that this moment of surprise might have even modified your memory of another snowy morning in the past (Exton-McGuinness et al., 2015; Gershman et al., 2017; Gershman et al., 2014; Sinclair & Barense, 2018). To understand and explain the computational role (Marr, 1982) of surprise in different brain functions, one first needs to ask ‘what does it really mean to be surprised?’ and formalize how surprise is perceived by our brain. For instance, when you see the white street, do you feel ‘surprised’ because what you expected turned out to be wrong (Faraji et al., 2018; Gläscher et al., 2010; Shannon, 1948; Tribus, 1961) or because you need to change your trust in the weather forecast (Baldi, 2002; Liakoni et al., 2021; Schmidhuber, 2010)?

Computational models of perception, learning, memory, and decision-making often assume that humans implicitly perceive their sensory observations as probabilistic outcomes of a generative model with hidden variables (Dubey & Griffiths, 2019; Findling et al., 2021; Fiser et al., 2010; Friston, 2010; Gershman et al., 2017; Liakoni et al., 2021; Schmidhuber, 2010; Soltani & Izquierdo, 2019; Yu & Dayan, 2005). In the example above, the observation is whether it snows or not and the hidden variables characterise how the probability of snowing depends on old observations and relevant context information (such as the current season, yesterday’s weather, and the weather forecast). Different brain functions are then modeled as aspects of statistical inference (Barber, 2012) and probabilistic control (Sutton & Barto, 2018) in such generative models (Behrens et al., 2007; Daw et al., 2011; Dubey & Griffiths, 2019; Findling et al., 2021; Friston, 2010; Friston et al., 2017; Gershman et al., 2017; Gläscher et al., 2010; Horvath et al., 2021; Liakoni et al., 2021; Meyniel et al., 2016; Nassar et al., 2012; Rouhani et al., 2020; Ryali et al., 2018; Schmidhuber, 2010; Soltani & Izquierdo, 2019; Yu & Dayan, 2005). In these probabilistic settings, surprise of an observation depends on the relation between the observation and our expectation of what to observe.

In the past decades, different definitions and formal measures of surprise have been proposed and studied (Baldi, 2002; Barto et al., 2013; Faraji et al., 2018; Friston, 2010; Gläscher et al., 2010; Itti & Baldi, 2006, 2009; Kolossa et al., 2015; Liakoni et al., 2021; Palm, 2012; Schmidhuber, 2010; Shannon, 1948; Tribus, 1961). These surprise measures have been successful both in explaining the role of surprise in different brain functions (Findling et al., 2021; Friston, 2010; Gershman et al., 2017; Liakoni et al., 2021; Schmidhuber, 2010; Soltani & Izquierdo, 2019; Yu & Dayan, 2005) and in identifying signatures of surprise in behavioral and physiological measurements (Gijsen et al., 2021; Gläscher et al., 2010; Kolossa et al., 2015; Lieder et al., 2013; Maheu et al., 2019; Mars et al., 2008; Meyniel, 2020; Meyniel et al., 2016; Modirshanechi et al., 2019; Mousavi et al., 2020; Ostwald et al., 2012; Preuschoff et al., 2011; Rubin et al., 2016). However, there are still many open questions (Baldi, 2002; Barto et al., 2013; Liakoni et al., 2021; Palm, 2012; Schmidhuber, 2010) including, but not limited to: (i) Since the notion of surprise has been linked to a multitude of computational roles and physiological phenomena, should we still assume that the word ‘surprise’ (and quantitative measures thereof) refers to a single common phenomenon or rather to a multitude of different phenomena? (ii) If ‘surprise’ refers to a multitude of different phenomena, can we assign different measures of surprise to different computational roles and different phenomena? In other words, are different surprise measures qualitatively distinguishable in experiments? (iii) Can we identify mathematical relations between different surprise measures? In particular, is one measure a special case of another one, completely distinct, or do they have some common ground? (iv) Does the word surprise as used in common language match one of the known mathematical definitions?

In this work, we analyze and discuss several previously proposed surprise measures (Baldi, 2002; Burda et al., 2019; Faraji et al., 2018; Friston, 2010; Gläscher et al., 2010; Itti & Baldi, 2006, 2009; Kolossa et al., 2015; Liakoni et al., 2021; Pathak et al., 2017; Schmidhuber, 2010; Shannon, 1948) in a unifying framework. First, we give definitions for each of these measures, show their similarities and differences, and prove and discuss their mathematical properties. We then propose experimental paradigms where different surprise measures entail different predictions for perception, learning, and decision-making. Our theoretical analysis provides a comprehensive understanding of current paradigms (Kuhn, 1962) for the study of surprise, shows that a single notion of surprise cannot explain the multitude of computational roles and physiological phenomena to which surprise has been linked, and shows how different measures of surprise relate to how we use the word ‘surprise’ in common language. Furthermore, we suggest a new definition of surprise which can capture some intuitive aspects of surprise perception that cannot be explained by current definitions. Our analyses enable experimental scientists to dissociate the contributions of different surprise measures to behavioral and physiological measurements.

### Subjective world-model: A unifying generative model

Our primary goal is to study the theoretical properties of different formal measures of surprise and their predictions for perception, learning, and decision-making in a common mathematical framework. To do so, we consider a generative model of a volatile environment (Fig. 1A) that captures a few key features of daily life and unifies many existing model environments in neuroscience and psychology (Behrens et al., 2007; Daw et al., 2011; Findling et al., 2021; Gijsen et al., 2021; Gläscher et al., 2010; Glaze et al., 2015; Heilbron & Meyniel, 2019; Horvath et al., 2021; Huys et al., 2015; Liakoni et al., 2021; Mars et al., 2008; Meyniel et al., 2016; Nassar et al., 2012; Nassar et al., 2010; Ostwald et al., 2012; Wilson et al., 2013; Xu et al., 2021). The generative model describes the subjective interpretation of the environment from the point of view of an agent (e.g., a human participant or an animal). Importantly, we assume that the agent takes the possibility into account that the environment may undergo abrupt changes at unknown points in time (similar to Glaze et al., 2015; Heilbron and Meyniel, 2019; Liakoni et al., 2021; Nassar et al., 2010; Xu et al., 2021). Note, however, that we do not assume that the environment has the same dynamics as those assumed by the agent.

**Figure 1:**
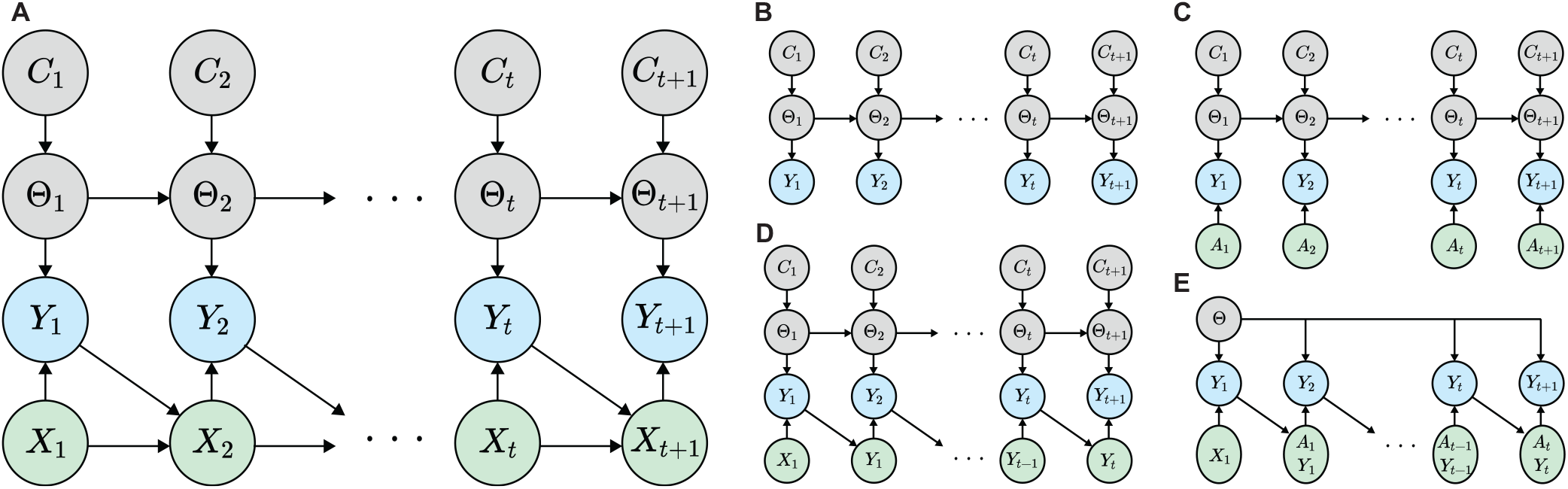
Subjective model of the environment. **A**. The Bayesian network (Barber, 2012) corresponding to the most general case of our generative model in Eq. 1 and Eq. 2. The arrows show conditional dependence, the grey nodes show the hidden variables (*C*_1:*t*+1_ and Θ_1:*t*+1_), the blue nodes show the observations (*Y*_1:*t*+1_), and the green nodes show the cue variables (*X*_1:*t*+1_). A variety of tasks can be written in the form of a reduced version of our generative model. Specifically: **B**. Standard generative model for studying passive learning in volatile environments (Adams & MacKay, 2007; Fearnhead & Liu, 2007; Liakoni et al., 2021; Nassar et al., 2012; Nassar et al., 2010; Wilson et al., 2013), **C**. Generative model corresponding to variants of bandit and reversal bandit tasks (Behrens et al., 2007; Findling et al., 2021; Horvath et al., 2021), where the cue variable *X*_*t*_ = *A*_*t*_ is a participant’s action, **D**. Generative model for modeling human inferences about binary sequences (Gijsen et al., 2021; Maheu et al., 2019; Meyniel et al., 2016; Modirshanechi et al., 2019; Mousavi et al., 2020), and **E**. classic Markov Decision Processes (MDPs) (Daw et al., 2011; Gläscher et al., 2010; Huys et al., 2015; Lehmann et al., 2019; Schultz et al., 1997; Sutton & Barto, 2018), where the cue variable *X*_*t*_ = (*A*_*t*−1_, *Y*_*t*−1_) consists of previous action and observation. See Appendix A: Special cases and links to related works for details.

### General definition

At each time *t*, the agent’s model of the environment is characterized by a tuple of 4 random variables (*X*_*t*_, *Y*_*t*_, Θ_*t*_, *C*_*t*_) (Fig. 1A). *X*_*t*_ and *Y*_*t*_ are observable, whereas Θ_*t*_ and *C*_*t*_ are unobservable (hidden). We refer to *X*_*t*_ as the cue and to *Y*_*t*_ as the observation at time *t*. Examples of an observation are an image on a computer screen (da Silva & Hare, 2020; Daw et al., 2011; Gläscher et al., 2010; Kolossa et al., 2015; Mars et al., 2008; Xu et al., 2021), an auditory tone (Imada et al., 1993; Lieder et al., 2013; Meyniel et al., 2016; Walz et al., 2015), and an electrical or mechanical stimulation (Esmaeili et al., 2021; Gijsen et al., 2021; Ostwald et al., 2012). The cue variable *X*_*t*_ can be interpreted as a predictor of the next observation, since it describes all observable information that is relevant for predicting the observation *Y*_*t*_. Examples of a cue variable are the previous observation *Y*_*t*−1_ (Maheu et al., 2019; Meyniel et al., 2016; Modirshanechi et al., 2019), the last action of a participant (which we will denote by *A*_*t*−1_) (Behrens et al., 2007; Findling et al., 2021; Horvath et al., 2021), and a conditioned stimulus in Pavlovian conditioning tasks (Gershman et al., 2017; Misanin et al., 1968).

At time *t*, given the cue variable *X*_*t*_ = *x*, the agent assumes that the observation *Y*_*t*_ = *y* comes from a time-invariant conditional distribution *P*_*Y* |*X*_(*y*|*x*; *θ*) parameterized by the hidden variable Θ_*t*_ = *θ*. We do not put any constraints on the sets to which *x, y*, and *θ* belong; in a non-parametric setup, *θ* can even have infinite dimension (Gershman & Blei, 2012; Ghahramani, 2013). We refer to Θ_*t*_ as the environment parameter at time *t*. The sequence of variables Θ_1:*t*_ = [Θ_1_, …, Θ_*t*_] describe the temporal dynamics of the observations *Y*_1:*t*_ given the cue variables *X*_1:*t*_ in the agent’s model of the environment. Similar to well-known models of volatile environments (Adams & MacKay, 2007; Behrens et al., 2007; Fearnhead & Liu, 2007; Findling et al., 2021; Glaze et al., 2015; Heilbron & Meyniel, 2019; Liakoni et al., 2021; Meyniel et al., 2016; Nassar et al., 2012; Nassar et al., 2010; Wilson et al., 2013; Xu et al., 2021; Yu & Dayan, 2005), the agent assumes that the environment undergoes abrupt changes at random points in time. An abrupt change at time *t* is specified by the event *C*_*t*_ = 1 and happens with a probability *p*_*c*_ ∈ [0, 1); otherwise *C*_*t*_ = 0. If the environment abruptly changes at time *t* (i.e., *C*_*t*_ = 1), then the agent assumes that the environment parameter Θ_*t*_ is sampled from a prior distribution *π*^(0)^ independently of Θ_*t*−1_; if there is no change (*C*_*t*_ = 0), then Θ_*t*_ remains the same as Θ_*t*−1_. We refer to *p*_*c*_ as the change-point probability.

We use ℙ to refer to probability mass functions (pmf) for discrete random variables and to probability density functions (pdf) for continuous random variables. In general, we show random variables by capital letters and their values by small letters. However, for any pair of arbitrary random variables *W* and *V* and their values *w* and *v*, whenever there is no risk of ambiguity, we either drop the capital- or the small-letter notation and, for example, write ℙ(*W* = *w*|*V* = *v*) as ℙ(*w*|*v*). When there is a risk of ambiguity, we keep the capital notation for the random variables, e.g., we write ℙ(*W* = *v, V* = *v*) as ℙ(*W* = *v, v*). Given this convention, the agent’s model of the environment described above is formalized in Definition 1.

#### Definition 1.

*(Subjective world-model) An agent’s model of the environment is defined for t >* 0 *as a joint probability distribution over Y*_1:*t*_, *X*_1:*t*_, Θ_1:*t*_, *and C*_1:*t*_ *as*

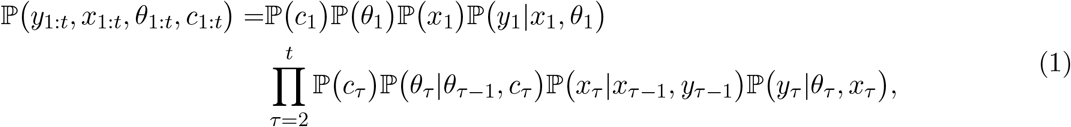

*where c*_1_ *is by definition equal to* 1 *(i*.*e*., 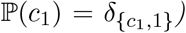, ℙ(*θ*_1_) = *π*^(0)^(*θ*_1_) *for an arbitrary distribution π*^(0)^, *and*

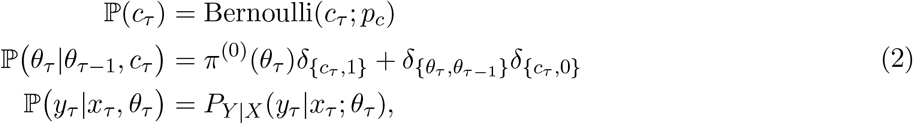

*where δ is the delta distribution. We do not make any assumption about* ℙ(*x*_1_) *and* ℙ(*x*_*τ*_ |*x*_*τ*−1_, *y*_*τ*−1_).

See Table 1 for a summary of the notation. Fig. 1B-E shows how the generative model of Definition 1 relates to other existing model environments in neuroscience and psychology – see Appendix A: Special cases and links to related works for details.

**Table 1:**
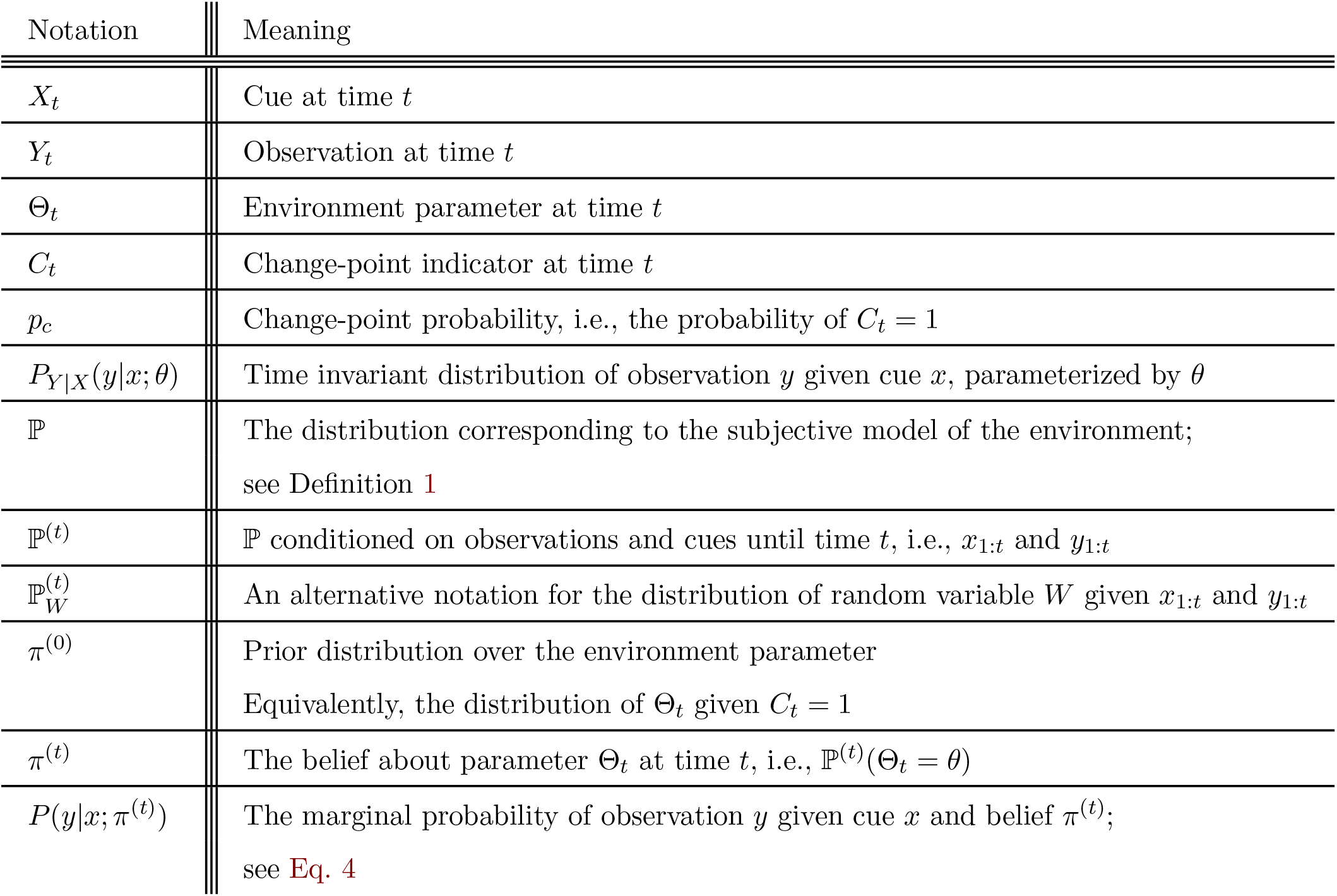
Summary of notation introduced in Subjective world-model: A unifying generative model

### Additional notation, belief, and marginal probability

We define ℙ^(*t*)^ as ℙ conditioned on the sequences of observations *y*_1:*t*_ and cue variables *x*_1:*t*_. For example, for an arbitrary random variable *W* with value *w*, we write ℙ(*w*|*y*_1:*t*_, *x*_1:*t*_) = ℙ^(*t*)^(*w*). We occasionally use an alternative notation and write ℙ^(*t*)^(*W* = *w*) as 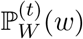 where the random variable is specified in the subscript.

Following this notation, we define an agent’s belief about the parameter Θ_*t*_ at time *t* as the posterior distribution

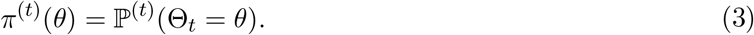

We can equivalently write 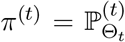. The belief plays a crucial role in the perception of surprise (see next sections), and we assume that an agent constantly updates its belief as it makes new observations; see Barber, 2012 and Liakoni et al., 2021 for examples of inference algorithms in generative models similar to ours.

Another important quantity is the marginal probability of observing *y* given the cue *x* and a belief *π*^(*t*)^:

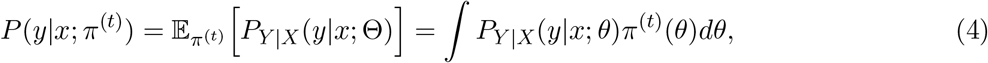

where the integration is replaced by summation whenever *θ* is discrete.

## Theories of surprise: A technical review

Conditioned on the previous observations *y*_1:*t*_ and cue variables *x*_1:*t*+1_, how surprising is the next observation *y*_*t*+1_? In this section, we address this question by examining previously proposed measures of surprise (Baldi, 2002; Burda et al., 2019; Faraji et al., 2018; Friston, 2010; Gläscher et al., 2010; Itti & Baldi, 2006, 2009; Kolossa et al., 2015; Liakoni et al., 2021; Pathak et al., 2017; Schmidhuber, 2010; Shannon, 1948). We define all surprise measures in the same mathematical framework and discuss their differences and similarities.

Surprise measures are commonly used in experiments to study whether a behavioral or physiological variable *Z* (e.g., the amplitude of the EEG P300 component (Kolossa et al., 2015; Meyniel et al., 2016)) is sensitive to or representative of surprise. Given two measures of surprise 𝒮 and 𝒮′, a typical experimental question is which one of them (if any) more accurately explains the variations of the variable *Z* (Gijsen et al., 2021; Kolossa et al., 2015; Mousavi et al., 2020; Ostwald et al., 2012; Visalli et al., 2021); see Fig. 2A1. However, if there exists a one-to-one (monotonic) mapping between 𝒮 and 𝒮′ (e.g., as in Fig. 2A2), then the two surprise measures have the same explanatory power with respect to *Z* – because any function of 𝒮 can be written in terms of 𝒮′ and vice-versa^1^. We, therefore, say 𝒮 and 𝒮′ are indistinguishable if there exists a strictly increasing function *f* : ℝ → ℝ such that 𝒮 = *f*(𝒮′) for all choices of belief *π*^(*t*)^, cue *x*_*t*_, and observation *y*_*t*_ (c.f. Fig. 2A2). Loosely speaking, we consider two surprise measures indistinguishable if whenever 𝒮 increases 𝒮′ increases as well. One of our goals in this section is to determine under what conditions different surprise measures are indistinguishable (Fig. 2B).

**Figure 2:**
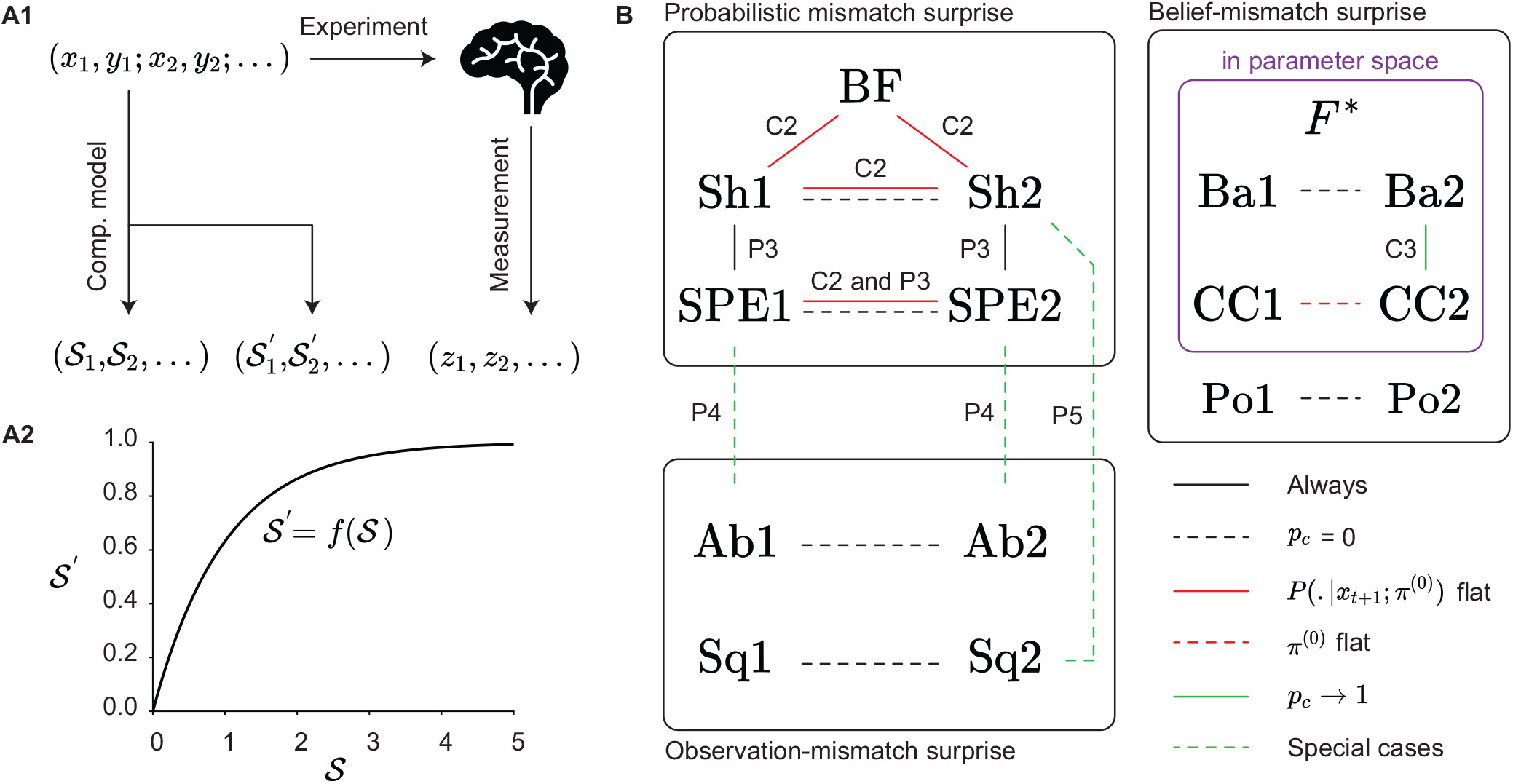
Indistinguishable surprise measures. **A**. A typical question in human and animal experiments is whether a surprise measure 𝒮 explains the variations of a behavioral or physiological variable *Z* better than an alternative surprise measure 𝒮′. **A1**. A common experimental paradigm: A sequence of cues *x*_1:*t*_ and observations *y*_1:*t*_ is presented to participants, the sequence *z*_1:*t*_ is measured, and the sequence of surprise values 𝒮_1:*t*_ or 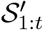 is predicted by computational modeling. Then statistical tools are used to study whether the sequence 𝒮_1:*t*_ or 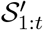 is more informative about the sequence of measurements *z*_1:*t*_. **A2**. If there exists a strictly increasing function *f* such that 𝒮′ = *f*(𝒮), then the two surprise measures are equally informative about the measurable variable *Z*. In this case, 𝒮 and 𝒮′ are ‘indistinguishable’. **B**. Schematic of the theoretical relation between different measures of surprise. A line connecting two measures indicates that the two measures are indistinguishable, i.e., one is a strictly increasing function of the other under the condition corresponding to the color and the type of the line. The conditions are shown on the bottom right of the panel: a solid black line means the two measures are always indistinguishable; a dashed black line corresponds to the condition *p*_*c*_ = 0; a solid red line corresponds to the prior marginal probability *P* (.|*x*_*t*+1_; *π*^(0)^) being flat; a dashed red line corresponds to the prior belief *π*^(0)^ being flat; a solid green line corresponds to the limit of *p*_*c*_ → 1; and a dashed green line means that the relation holds only for some special cases (e.g., Gaussian task). Two lines indicate that one of the conditions is sufficient for the two measures to be indistinguishable. The text beside each line shows where in the text the existence of the mapping is proven, e.g., P3 and C2 stand for Proposition 3 and Corollary 2, respectively. The purple box includes surprise measures that are computed in the parameter (Θ_*t*_) space, whereas the surprise measures outside of the purple box are computed in the space of observations (*Y*_*t*_). Abbreviations: Ab: Absolute error surprise, Sq: Squared error surprise, BF: Bayes Factor surprise, Sh: Shannon surprise, SPE: State Prediction Error, Ba: Bayesian surprise, Po: Postdictive surprise, CC: Confidence Corrected surprise, and *F*^∗^: Minimized Free Energy. See section Theories of surprise: A technical review for details.

Based on how they depend on an agent’s belief *π*^(*t*)^, we divide existing surprise measures into three categories: (i) probabilistic mismatch, (ii) observation-mismatch, and (iii) belief-mismatch surprise measures (Fig. 3). *Probabilistic mismatch* surprise measures depend on the belief *π*^(*t*)^ through the marginal probability *P*(*y*_*t*+1_|*x*_*t*+1_; *π*^(*t*)^); an example is the Shannon surprise (Shannon, 1948). *Observation-mismatch* surprise measures depend on the belief *π*^(*t*)^ through some estimate of the next observation *ŷ*_*t*+1_ according to the belief *π*^(*t*)^; an example is the absolute difference between *y*_*t*+1_ and *ŷ*_*t*+1_ (Rouhani & Niv, 2021). To compute the *belief-mismatch* surprise measures, however, we need to have the whole distribution *π*^(*t*)^; an example is the Bayesian surprise (Baldi, 2002; Schmidhuber, 2010).

**Figure 3:**
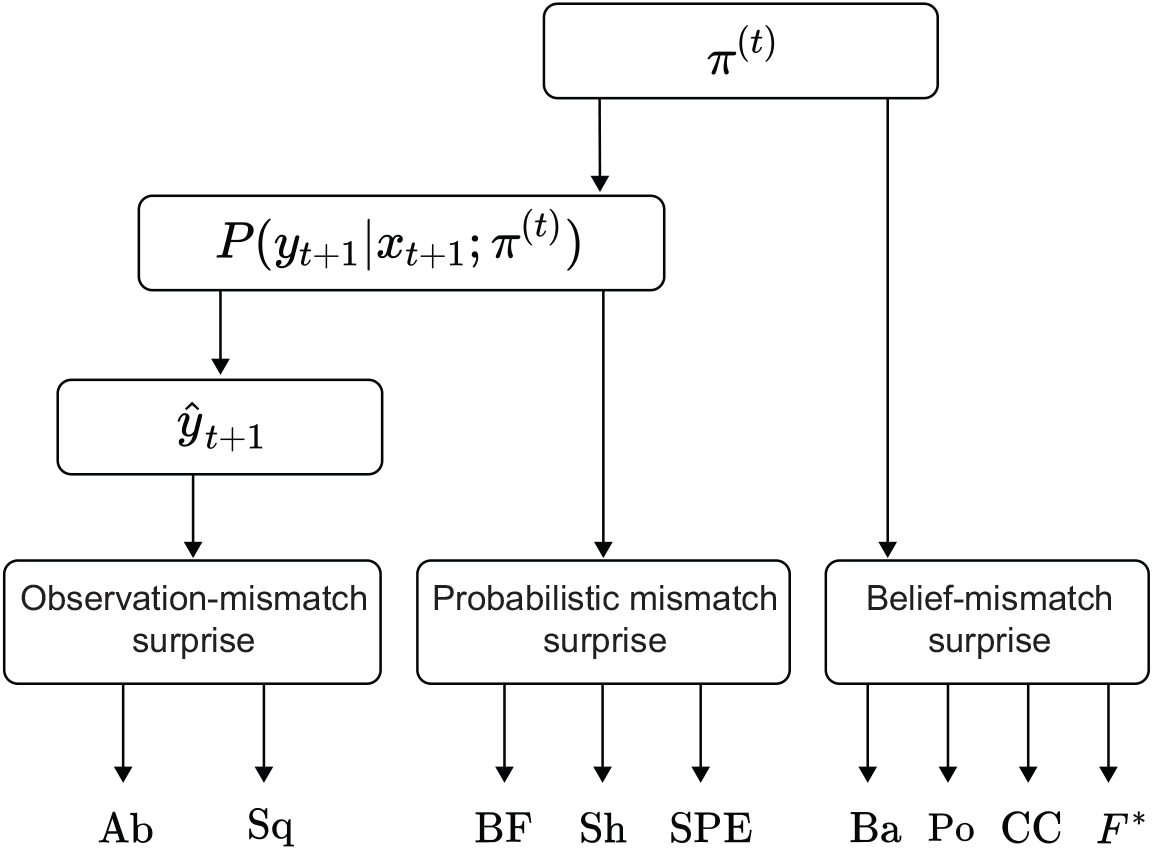
Categorization of surprise measures based on the form of their dependence upon the belief. Surprise depends on expectations. Therefore, all surprise measures depend on the belief *π*^(*t*)^. However, the specific form of the dependence changes between one measure and another. ‘Observation-mismatch’ surprise measures use the marginal distribution *P*(*y*|*x*_*t*+1_; *π*^(*t*)^) to calculate an estimate *ŷ*_*t*+1_ of the next observation, which is then compared with the real observation *y*_*t*+1_ by an error function such as ||*ŷ*_*t*+1_ − *y*_*t*+1_||_1_, where ||.||_1_ stands for *ℓ*_1_-norm. ‘Probabilistic mismatch’ surprise measures use the marginal probability *P*(*y*_*t*+1_|*x*_*t*+1_; *π*^(*t*)^) directly, without extracting a specific estimate. ‘Belief-mismatch’ surprise measures use the belief *π*^(*t*)^ directly, without extracting the marginal distribution *P*(*y*|*x*_*t*+1_; *π*^(*t*)^). Abbreviations: Ab: Absolute error surprise, Sq: Squared error surprise, BF: Bayes Factor surprise, Sh: Shannon surprise, SPE: State Prediction Error, Ba: Bayesian surprise, Po: Postdictive surprise, CC: Confidence Corrected surprise, and *F*^∗^: Minimized Free Energy. See section Theories of surprise: A technical review for details.

### Probabilistic mismatch surprise 1: Bayes Factor surprise

An abrupt change in the parameters of the environment influences the sequence of observations. Therefore, a sensible way to define the surprise of an observation is that ‘surprise’ measures the probability of an abrupt change in the eye of the agent, given the present observation. To detect an abrupt change, it is not enough to measure how unexpected the observation is according to the current belief of the agent. Rather, the agent should measure how much more expected the new observation is under the prior belief than under the current belief. The Bayes Factor surprise was introduced by Liakoni et al., 2021 to quantify this concept of surprise.

Here, we apply their definition to our generative model. Similar to Xu et al., 2021, we define the Bayes Factor surprise of observing *y*_*t*+1_ given the cue *x*_*t*+1_ as the ratio of the marginal probability of observing *y*_*t*+1_ given *x*_*t*+1_ and *C*_*t*+1_ = 1 (i.e., assuming a change) to the marginal probability of observing *y*_*t*+1_ given *x*_*t*+1_ and *C*_*t*+1_ = 0 (i.e. assuming no change); formally, we write

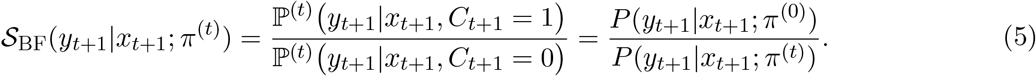

The name arises because 𝒮_BF_(*y*_*t*+1_|*x*_*t*+1_; *π*^(*t*)^) is the Bayes Factor (Bayarri & Berger, 1997; Efron & Hastie, 2016; Kass & Raftery, 1995) used in statistics to test whether a change has occurred at time *t*. For a given *P*(*y*_*t*+1_|*x*_*t*+1_; *π*^(0)^), the Bayes Factor surprise is a decreasing function of *P*(*y*_*t*+1_|*x*_*t*+1_; *π*^(*t*)^): Hence, more probable events are perceived as less surprising. However, the key feature of 𝒮_BF_(*y*_*t*+1_|*x*_*t*+1_; *π*^(*t*)^) is that it measures not only how unexpected (unlikely) the observation *y*_*t*+1_ is according to the current belief *π*^(*t*)^ but also how expected it would be if the agent had reset its belief to the prior belief. More precisely, for a given *P*(*y*_*t*+1_|*x*_*t*+1_; *π*^(*t*)^), the Bayes Factor surprise is an increasing function of *P*(*y*_*t*+1_|*x*_*t*+1_; *π*^(0)^).

Such a comparison is necessary to evaluate whether a reset of the belief (or an increase in the update rate of the belief) can be beneficial in order to have a more accurate estimate of the environment’s parameters (c.f. Soltani and Izquierdo, 2019). This intuition is formulated in a precise way by Liakoni et al., 2021 in their Proposition 1, where they show that, in the generative model of Fig. 1B, the exact Bayesian inference for the update of *π*^(*t*)^ to *π*^(*t*+1)^ upon observing *y*_*t*+1_ leads to a learning rule modulated by the Bayes Factor surprise. Proposition 1 below states that this result is also true for our more general generative model (Definition 1 and Fig. 1A).

#### Proposition 1.

*(Extension of Proposition 1 of Liakoni et al., 2021) For the generative model of Definition 1, the Bayes Factor surprise can be used to write the updated (according to exact Bayesian inference) belief π*^(*t*+1)^, *after observing y*_*t*+1_ *with the cue x*_*t*+1_, *as*

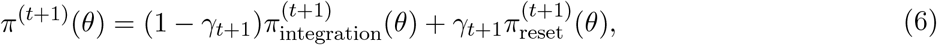

*where γ*_*t*+1_ *is an adaptation rate modulated by the Bayes Factor surprise*

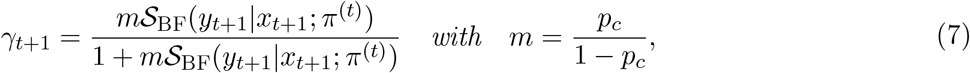

*and*

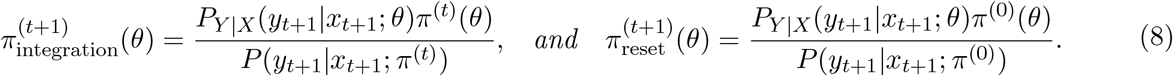

Therefore, the Bayes Factor surprise 𝒮_BF_ controls the trade-off between the integration of the new observation into the old belief 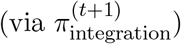 and resetting the old belief to the prior belief 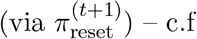 Liakoni et al., 2021.

### Probabilistic mismatch surprise 2: Shannon surprise

No matter if there has been an abrupt change (*C*_*t*+1_ = 1) or not (*C*_*t*+1_ = 0), an unlikely event may be perceived as surprising. Therefore, another way to measure the surprise of an observation is to quantify how unlikely the observation is in the eye of the agent. Shannon surprise (Shannon, 1948), also known as surprisal (Tribus, 1961), is a way to formalize this concept of surprise. It comes from the field of information theory (Shannon, 1948) and statistical physics (Tribus, 1961) and is widely used in neuroscience (Gijsen et al., 2021; Kolossa et al., 2015; Konovalov & Krajbich, 2018; Kopp & Lange, 2013; Maheu et al., 2019; Mars et al., 2008; Meyniel et al., 2016; Modirshanechi et al., 2019; Mousavi et al., 2020; Visalli et al., 2021).

Formally, in the generative model of Definition 1, one can define the Shannon surprise of observing *y*_*t*+1_ given the cue *x*_*t*+1_ as

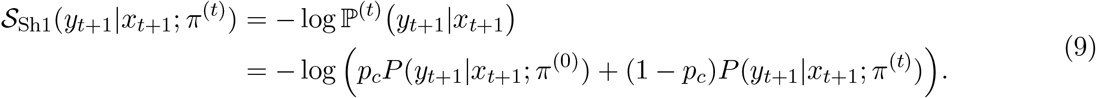

The Shannon surprise 𝒮_Sh1_ measures how unexpected or unlikely *y*_*t*+1_ is considering the possibility that there might have been an abrupt change in the environment. As a result, for a fixed *P*(*y*_*t*+1_|*x*_*t*+1_; *π*^(*t*)^), the Shannon surprise is a decreasing function of *P*(*y*_*t*+1_|*x*_*t*+1_; *π*^(0)^) (c.f. Eq. 9), whereas the Bayes Factor surprise is an increasing function of *P*(*y*_*t*+1_|*x*_*t*+1_; *π*^(0)^) (c.f. Eq. 5). This essential difference between the Shannon and the Bayes Factor surprise has been exploited by Liakoni et al., 2021 in order to propose experiments where these two measures of surprise make different predictions.

Experimental evidence (Nassar et al., 2012; Nassar et al., 2010; Wilson et al., 2013) indicates that in volatile environments like the one in Fig. 1B, human participants do not actively consider the possibility that there *may be* an abrupt change while predicting the *next* observation *y*_*t*+1_ – even though they update their belief after observing *y*_*t*+1_ by considering the possibility that there *might have been* a change before the *current* observation at time *t* + 1. To arrive at a Shannon surprise measure consistent with this observation, we suggest a 2nd definition as

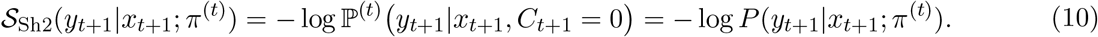

In other words, 𝒮_Sh2_(*y*_*t*+1_|*x*_*t*+1_; *π*^(*t*)^) neglects the potential presence of change-points, and, therefore, it is independent of both *p*_*c*_ and *P*(*y*_*t*+1_|*x*_*t*+1_; *π*^(0)^). For a non-volatile environment that does not allow for abrupt changes (*p*_*c*_ = 0), the two definitions of Shannon surprise are identical: 𝒮_Sh1_ = 𝒮_Sh2_ (Fig. 2B).

Proposition 2 shows that the Bayes Factor surprise 𝒮_BF_ is related to 𝒮_Sh1_ and 𝒮_Sh2_:

#### Proposition 2.

*(Relation between the Shannon surprise and the Bayes Factor surprise) In the generative model of Definition 1, the Bayes Factor surprise* 𝒮_BF_(*y*_*t*+1_|*x*_*t*+1_; *π*^(*t*)^) *can be written as*

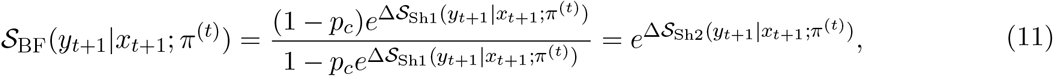

*where*

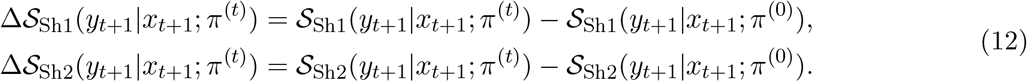

Proposition 2 states that the Bayes Factor 𝒮_BF_(*y*_*t*+1_|*x*_*t*+1_; *π*^(*t*)^) has a behavior similar to the *difference* in Shannon surprise (i.e., Δ𝒮_Sh1_ or Δ𝒮_Sh2_) as opposed to Shannon surprise itself (i.e., 𝒮_Sh1_ or 𝒮_Sh2_). The difference in Shannon surprise (i.e., Δ𝒮_Sh1_ or Δ𝒮_Sh2_) compares the Shannon surprise under the current belief with that under the prior belief. Two direct consequences of this proposition are summarized in Corollaries 1 and 2.

Corollary 1 states that the modulation of learning as presented in Proposition 1 can also be written in the form of the difference in Shannon surprise (i.e., Δ𝒮_Sh1_ or Δ𝒮_Sh2_).

#### Corollary 1.

*The adaptation rate γ*_*t*+1_ *in Proposition 1 can be written as*

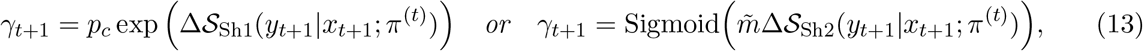

*with* 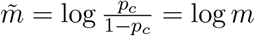 *(c*.*f. Proposition 1) and* 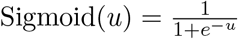

Corollary 2 indicates that, under a flat prior, the Bayes Factor surprise and the two definitions of the Shannon surprise are indistinguishable from each other (Fig. 2B):

#### Corollary 2.

*(Flat prior prediction) In the generative model of Definition 1, if the probability of observing y*_*t*+1_ *with the cue x*_*t*+1_ *is flat under the prior belief π*^(0)^ *(i*.*e*., *if P*(*y*_*t*+1_|*x*_*t*+1_; *π*^(0)^) *is uniform), then there are one-to-one mappings between* 𝒮_BF_(*y*_*t*+1_|*x*_*t*+1_; *π*^(*t*)^), 𝒮_Sh1_(*y*_*t*+1_|*x*_*t*+1_; *π*^(*t*)^), *and* 𝒮_Sh2_(*y*_*t*+1_|*x*_*t*+1_; *π*^(*t*)^).

A consequence of Corollary 2 is that experimental paradigms with flat priors of the agent cannot be used to distinguish 𝒮_BF_ from 𝒮_Sh1_ or 𝒮_Sh2_ (c.f. Fig. 2).

### Probabilistic mismatch surprise 3: State prediction error

The State Prediction Error (SPE) was introduced by Gläscher et al., 2010 in the context of model-based reinforcement learning in Markov Decision Processes (MDPs – c.f. Fig. 1E) (Daw et al., 2011; Daw et al., 2005; Sutton & Barto, 2018). Similar to the Shannon surprise, the SPE considers less probable events as the more surprising ones.

Whenever observations *y*_1:*t*_ come from a discrete distribution so that we have *P*_*Y* |*X*_ (*y*_*t*+1_|*x*_*t*+1_; *θ*) ∈ [0, 1] for all *θ, x*_*t*+1_, and *y*_*t*+1_, we can generalize the definition of Gläscher et al., 2010 to the setting of our generative model. Analogously to our two definitions of Shannon surprise (c.f. Eq. 9 and Eq. 10), we give also two definitions for SPE:

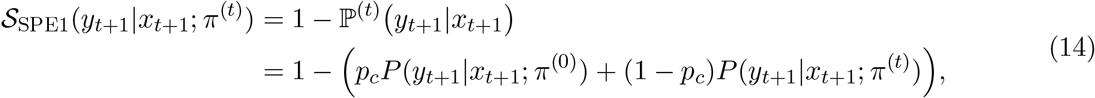

and

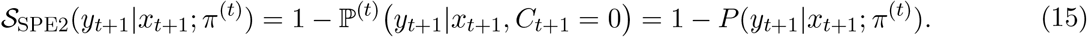

In non-volatile environments (*p*_*c*_ = 0), the two definitions of SPE are identical (Fig. 2B). In particular, in an MDP without abrupt changes (*p*_*c*_ = 0; Fig. 1E), both definitions are equal to 1 −ℙ ^(*t*)^(*s*_*t*_, *a*_*t*_ → *s*_*t*+1_), where ℙ ^(*t*)^(*s*_*t*_, *a*_*t*_ → *s*_*t*+1_) is an agent’s estimate (at time *t*) of the probability of the transition to state *s*_*t*+1_ after taking action *a*_*t*_ in state *s*_*t*_; c.f. Gläscher et al., 2010.

Proposition 3 states that both definitions (𝒮_SPE1_ and 𝒮_SPE2_) can always be written as invertible functions of Shannon surprise (Fig. 2B):

#### Proposition 3.

*(Relation between the Shannon surprise and the SPE) In the generative model of Definition 1, for i* ∈ {1, 2}, *the state prediction error* S_SPE*i*_(*y*_*t*+1_|*x*_*t*+1_; *π*^(*t*)^), *can be written as*

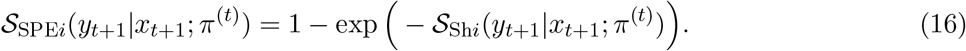

Therefore, the SPE and the Shannon surprise are in principle indistinguishable (Fig. 2).

### Observation-mismatch surprise: Absolute and squared errors

Assume an agent predicts *ŷ*_*t*+1_ for the next observation *y*_*t*+1_. Then, a measure of surprise can be defined as the prediction error or the mismatch between the prediction *ŷ*_*t*+1_ and the actual observation *y*_*t*+1_ (Heydari & Holroyd, 2016; Nassar et al., 2010; Prat-Carrabin et al., 2021; Rouhani et al., 2020; Talmi et al., 2013) (Fig. 3). For the sake of completeness, we discuss four possible definitions for observation-mismatch surprise measures in this section.

Before turning to an ‘observation-mismatch’, we first need to define an agent’s prediction for the next observation. Analogously to our two definitions for the Shannon surprise (c.f. Eq. 9 and Eq. 10), we define two different predictions for the next observation *y*_*t*+1_ given the cue *x*_*t*+1_:

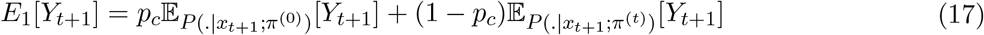

and

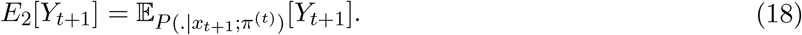

Although *E*_1_[*Y*_*t*+1_] is a more reasonable prediction for *y*_*t*+1_ given the fact that there is always a possibility of an abrupt change according to our generative model of the environment (Definition 1), Nassar et al., 2010 have shown that, in a Gaussian task, *E*_2_[*Y*_*t*+1_] explains human participants’ predictions better than *E*_1_[*Y*_*t*+1_].

We note that the observation *y*_*t*+1_ is in general case multi-dimensional (Niv et al., 2015). As two natural ways of measuring mismatch, we define the squared and the absolute error surprise, for *i* ∈ {1, 2} as

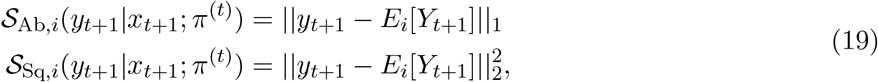

where ||.||_1_ and ||.||_2_ stand for the, *ℓ*_1_- and, *ℓ*_2_-norms, respectively, and *E*_1_ and *E*_2_ are defined in Eq. 17 and Eq. 18, respectively. Similar definitions have been used in neuroscience (Heydari & Holroyd, 2016; Nassar et al., 2010; Prat-Carrabin et al., 2021; Talmi et al., 2013) and machine learning (Burda et al., 2019; Pathak et al., 2017). For two special cases, we show in Propositions 4 and 5 below that the absolute and the squared error surprise can be written as functions of the SPE and the Shannon surprise (Fig. 2B).

#### Proposition 4.

*(Relation between the observation-mismatch surprise measures and the SPE for categorical distributions) In the generative model of Definition 1, if Y*_*t*+1_ *is represented as one-hot coded vectors, i*.*e*., *vectors with one element equal to 1 and the others equal to 0, then we have, for i* ∈ {1, 2},

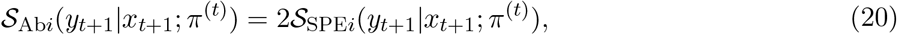

*and*

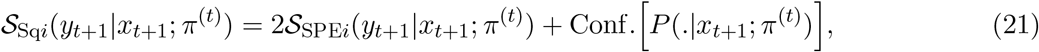

*where* Conf. *P* (.|*x*_*t*+1_; *π*^(*t*)^) *can be seen as a measure of confidence in the prediction (see Appendix B: Proofs)*.

#### Proposition 5.

*(Relation between the squared error surprise and the Shannon surprise for Gaussian distributions – from Pathak et al*., *2017) In the generative model of Definition 1, if the marginal distribution of Y*_*t*+1_ ∈ ℝ^*N*^ *given the cue x*_*t*+1_ *and under the belief π*^(*t*)^ *is a Gaussian distribution with a covariance matrix equal to σI*_*N*×*N*_, *where I*_*N*×*N*_ *is the N* × *N identity matrix, then* 𝒮_Sq2_(*y*_*t*+1_|*x*_*t*+1_; *π*^(*t*)^) *is an invertible function of* 𝒮_Sh2_(*y*_*t*+1_|*x*_*t*+1_; *π*^(*t*)^).

We note that, according to Proposition 3, the SPE is an invertible function of the Shannon surprise. Hence, for categorical distributions with one-hot coding, the SPE, the Shannon surprise, and the absolute error surprise are indistinguishable, and for Gaussian distributions with unit diagonal covariance, the SPE, the Shannon surprise, and the squared error surprise are indistinguishable (Fig. 2).

### Belief-mismatch surprise 1: Bayesian surprise

Another way to think about surprise is to define surprising events as those that change an agent’s belief about the world (Baldi, 2002; Storck et al., 1995). Bayesian surprise or information gain (Baldi & Itti, 2010; Itti & Baldi, 2006, 2009; Schmidhuber, 2010) is a way to formalize this concept of surprise. Whereas the Bayes Factor surprise measures how likely it is that the environment has changed given the new observation, the Bayesian surprise measures how much the agent’s belief changes given the new observation.

Bayesian surprise (Baldi, 2002; Itti & Baldi, 2006, 2009; Schmidhuber, 2010; Storck et al., 1995) has been originally introduced in non-volatile environments, i.e., where there is no change (*p*_*c*_ = 0) and as a result Θ_1_ = Θ_2_ = … = Θ_*t*_ = Θ. In this case, the Bayesian surprise of observing *y*_*t*+1_ with cue *x*_*t*+1_ is defined as 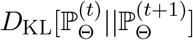 (Baldi, 2002; Itti & Baldi, 2006, 2009; Schmidhuber, 2010; Storck et al., 1995), where *D*_KL_ stands for the Kullback-Leibler (KL) divergence (Cover, 1999) (c.f. Table 1 for notation). Hence, in non-volatile environments, Bayesian surprise measures the pseudo-distance *D*_KL_ between two distributions, i.e., the belief 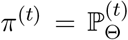 before and the belief 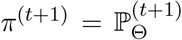 after observing *y*_*t*+1_. To generalize this definition to volatile environments, we have to choose the equivalent two distributions that we want to compare. The natural choice for 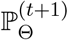 is 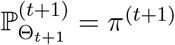; however, it is unclear whether 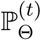 should be taken as the momentary belief 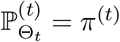 or its one-step forward-propagation 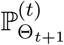the next observation *y*_*t*+1_ is integrated. If *p*_*c*_ ≠ 0, the two choices are different:

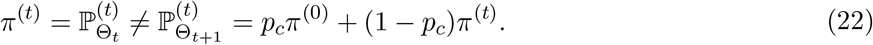

Therefore, for the case of volatile environments, we give two definitions for the Bayesian surprise:

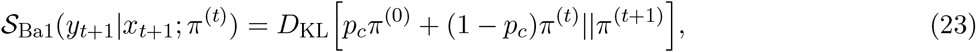

and

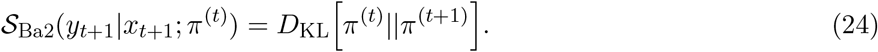

The 1st definition is more consistent with the original definition of the Bayesian surprise (Baldi, 2002; Itti & Baldi, 2006, 2009; Schmidhuber, 2010; Storck et al., 1995) applied to our generative model because the belief before the observation should include the knowledge that the environment is volatile. However, the 2nd definition looks more intuitive from the neuroscience perspective (Gijsen et al., 2021; Mousavi et al., 2020). Note that, in Eq. 23 and Eq. 24, the observation *y*_*t*+1_ does not appear explicitly on the right hand side; the observation has, however, influenced the update of the belief to its new distribution *π*^(*t*+1)^. For the case of *p*_*c*_ = 0, the two definitions are identical (Fig. 2B).

In Lemma 1 and Remark 1 in Appendix B: Proofs, we show that the Bayesian surprise is correlated with the difference between the Shannon surprise and its expectation (over all possible values of Θ_*t*+1_). There are two consequences of this observation. First, Bayesian surprise is distinguishable from Shannon surprise since it cannot be found only as a function of Shannon surprise. Second, we need access to the full belief distribution *π*^(*t*)^ for computing the expectation (Fig. 3).

In general, surprise measures similar to the Bayesian surprise can be defined also by measuring the change in the belief via distance or pseudo-distance measures different from the KL-divergence (Baldi, 2002; Baldi & Itti, 2010).

### Belief-mismatch surprise 2: Postdictive surprise

We saw that the Bayesian surprise measures how much the new belief *π*^(*t*+1)^ has changed after observing *y*_*t*+1_. Kolossa et al., 2015 introduced ‘postdictive surprise’ with a similar idea in mind but focused on changes in the marginal distribution *P* (*y*|*x*_*t*+1_; *π*^(*t*+1)^) (c.f. Eq. 4). More precisely, whereas the Bayesian surprise measures the amount of update in the space of distributions over the parameters (i.e., how differently the agent thinks about the parameters), the postdictive surprise measures the amount of update in the space of distributions over the observations (i.e., how differently the agent predicts the next observations).

Analogous to our two definitions for the Bayesian surprise (c.f. Eq. 23 and Eq. 24), there are two definitions for the postdictive surprise in volatile environments:

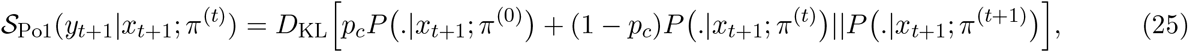

and

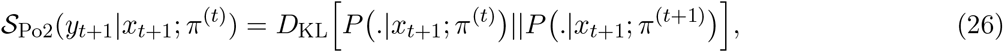

where the dot refers to a dummy variable *y* that is integrated out when evaluating *D*_KL_. Note that for *p*_*c*_ = 0, the two definitions are identical (Fig. 2B).

Although the amount of update is computed over the space of observations, 𝒮_Po1_ and 𝒮_Po2_ cannot be categorized as probabilistic mismatch surprise measures, since the update depends explicitly on the belief *π*^(*t*)^. The statement becomes more obvious in our Lemma 2 in Appendix B: Proofs.

### Belief-mismatch surprise 3: Confidence Corrected surprise

Since surprise arises when an expectation is violated, the violation of an agent’s expectation should be more surprising when the agent is more confident about its expectation. Based on the observation that neither Shannon nor Bayesian surprise explicitly captures the concept of confidence, Faraji et al., 2018 proposed the ‘Confidence Corrected Surprise’ as a new measure of surprise that explicitly takes confidence into account.

To define the Confidence Corrected surprise, we first define *π*_flat_ as the flat (uniform) distribution over the space of parameters, i.e., over the set to which Θ_*t*_ belongs. Then, following Faraji et al., 2018, we define the normalized likelihood after observing *y*_*t*+1_ (i.e., the posterior given the flat prior) as

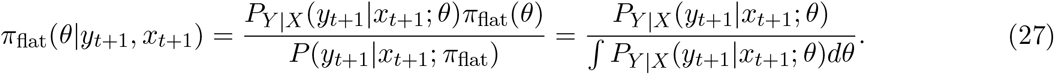

If the prior *π*^(0)^ is equal to *π*_flat_ (i.e., if the prior is uniform), then *π*_flat_(*θ*|*y*_*t*+1_, *x*_*t*+1_) is the same as 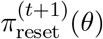 defined in Proposition 1. Note that the prior *π*_flat_ does not necessarily need to be a proper distribution (i.e., does not necessarily need to be normalized) as long as ∫ *P*_*Y* |*X*_ (*y*_*t*+1_|*x*_*t*+1_; *θ*)*dθ* is finite and the posterior *π*_flat_(.|*y*_*t*+1_, *x*_*t*+1_) is a proper distribution (Efron & Hastie, 2016). Using this terminology, the original definition for the Confidence Corrected surprise is (Faraji et al., 2018)

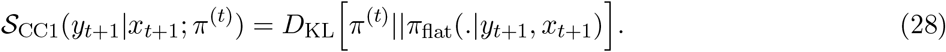

To interpret 𝒮_CC1_, Faraji et al., 2018 defined the commitment (or confidence) *C*[*π*] corresponding to an arbitrary belief *π* as its negative entropy (Cover, 1999), i.e.,

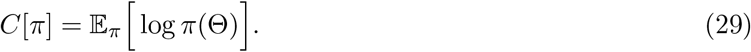

Then, in a non-volatile environment (i.e., *p*_*c*_ = 0), they show that 𝒮_CC1_ can be written as (Faraji et al., 2018)

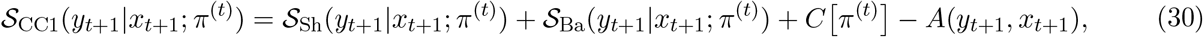

where *A*(*y*_*t*+1_, *x*_*t*+1_) = 𝒮_Sh_(*y*_*t*+1_|*x*_*t*+1_; *π*_flat_) + *C*[*π*_flat_] is independent of the current belief *π*^(*t*)^; note that because *p*_*c*_ = 0, we have 𝒮_Sh1_ = 𝒮_Sh2_ and 𝒮_Ba1_ = 𝒮_Ba2_. Therefore, in a non-volatile environment (i.e., *p*_*c*_ = 0), 𝒮_CC1_ is correlated with the sum of the Shannon and the Bayesian surprise regularized by the confidence of the agent’s belief. However, such an interpretation is no longer possible in volatile environments (*p*_*c*_ *>* 0), and Eq. 30 must be replaced by Proposition 6 below.

In order to account for the information of the true prior *π*^(0)^ and to avoid the cases where *π*_flat_(.|*y*_*t*+1_, *x*_*t*+1_) is not a proper distribution, we also give a 2nd definition for the Confidence Corrected surprise as

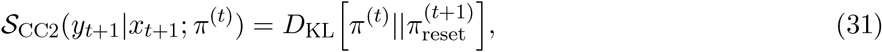

where 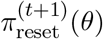 is defined in Proposition 1. Whenever *π*^(0)^ = *π*_flat_, the two definitions are identical (Fig. 1B). Proposition 6 shows how the Confidence Corrected surprise relates to the Shannon surprise, the Bayesian surprise, and the confidence in the general case.

#### Proposition 6.

*(Relation between the Confidence Corrected surprise, Shannon surprise, and Bayesian surprise) For the generative model of Definition 1, the original definition of the Confidence Corrected surprise can be written as*

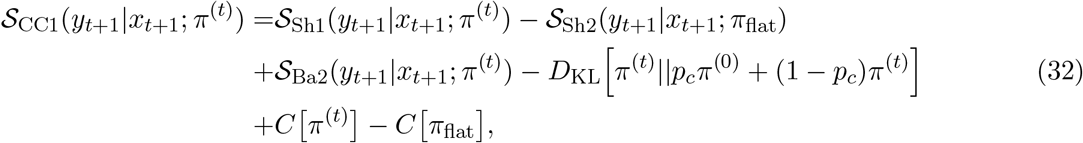

*and our 2nd definition can be written as*

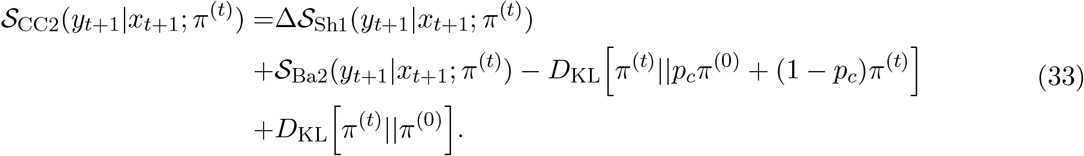

Proposition 6 conveys three important messages. First, both definitions of the Confidence Corrected surprise depend on differences in the Shannon surprise as opposed to the Shannon surprise itself (c.f. first line in Eq. 32 and Eq. 33). Second, both definitions depend on the difference between the Bayesian surprise (i.e., the change in the belief given the new observation) and the *a priori* expected change in the belief (because of the possibility of a change in the environment; c.f. second line in Eq. 32 and Eq. 33). Third, both definitions regularize the contributions of Shannon surprise and Bayesian surprise by the relative confidence of the current belief compared to either the flat or the prior belief (c.f. third line in Eq. 32 and Eq. 33). ‘Relative confidence’ quantifies how different the current belief is with respect to a reference belief; note that [*C π*^(*t*)^]– [*C π*_flat_]= *D*_KL_ [*π*^(*t*)^||*π*_flat_].

Hence, the Confidence Corrected surprise should be distinguishable from both the Shannon and the Bayesian surprise (for *p*_*c*_ *<* 1). An interesting consequence of Proposition 6, however, is that 𝒮_CC2_ is identical to 𝒮_Ba2_ when the environment becomes so volatile that its parameter changes at each time step (i.e., in the limit of *p*_*c*_ → 1):

#### Corollary 3.

*For the generative model of Definition 1, when p*_*c*_ → 1, *we have* 𝒮_CC2_(*y*_*t*+1_|*x*_*t*+1_; *π*^(*t*)^) = 𝒮_Ba2_(*yt*+1|*xt*+1; *π*^(*t*)^).

### Belief-mismatch surprise 4: Minimized free energy

Although an agent can perform computations over the joint probability distribution in Eq. 1 and Eq. 2, finding the belief *π*^(*t*+1)^(*θ*) (i.e., the posterior distribution in Eq. 3) can be computationally intractable (Barber, 2012; Liakoni et al., 2021). Therefore, it has been argued that the brain uses approximate inference (instead of exact Bayesian inference) for finding the belief (Daw & Courville, 2008; Faraji et al., 2018; Findling et al., 2021; Fiser et al., 2010; Friston, 2010; Friston et al., 2017; Liakoni et al., 2021; Mathys et al., 2011). An approximation of the belief *π*^(*t*+1)^(*θ*) can for example be found via variational inference (Blei et al., 2017; MacKay, 2003) over a family of distributions *q*(*θ*; *ϕ*) parameterized by *ϕ*. Such approaches are popular in neuroscience studies of learning and inference in the brain (Friston, 2010; Friston et al., 2017; Gershman, 2019).

Formally, in variational inference, the belief *π*^(*t*+1)^(*θ*) is approximated by 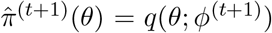, where *ϕ*^(*t*+1)^ is the minimizer of the variational loss or free energy, i.e., *ϕ*^(*t*+1)^ = arg min_*ϕ*_ *F* ^(*t*+1)^(*ϕ*) (Friston, 2010; Friston et al., 2017; Liakoni et al., 2021; MacKay, 2003; Markovic et al., 2021; Sajid et al., 2021). To define *F* ^(*t*+1)^(*ϕ*), we introduce a new notation:

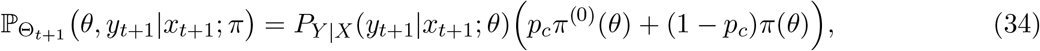

where *π*(*θ*) is an arbitrary distribution over the parameter space. Using this notation, we can write joint distribution over the observation and the parameter ℙ^(*t*)^(*θ*_*t*+1_, *y*_*t*+1_| *x*_*t*+1_) as 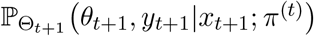 the and the updated belief 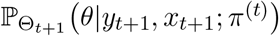. The variational loss or free energy can then be defined as (Friston, 2010; Friston et al., 2017; Liakoni et al., 2021; Markovic et al., 2021; Sajid et al., 2021)

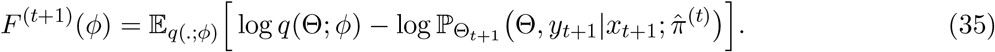

For any value of *ϕ*, one can show that (Blei et al., 2017; Sajid et al., 2021)

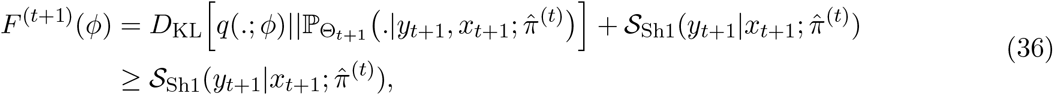

where the right side of the inequality is independent of *ϕ*, and 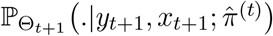 is the exact Bayesian update of the belief (according to the generative model in Definition 1) given the latest approximation of the belief 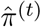 (Liakoni et al., 2021; Markovic et al., 2021).

The minimized free energy *F*^*^ = min_*ϕ*_ *F* ^(*t*+1)^(*ϕ*) has been interpreted as a measure of surprise (Friston, 2010; Friston et al., 2017; Schwartenbeck et al., 2013), which, according to Eq. 36, can be seen as an approximation of 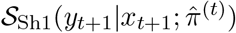. The parametric family of *q*(.; *ϕ*) and its relation to the exact belief *π*^(*t*+1)^ determine how well *F*^*^ approximates 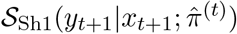 (Fig. 2B). More precisely, the minimized free energy measures both how unlikely the new observation is (i.e., how large 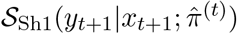 is) and how imprecise the best parametric approximation of the belief 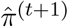 is (i.e., how large 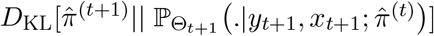 is) Therefore, the minimized free energy is in the category of belief-mismatch surprise measures (Fig. 3).

### A brief review of experimental results

The surprise definitions discussed above have been used in various experiments with the goal of identifying neural and behavioral signatures of surprise. In this subsection, we briefly review some of the existing experimental results.

The neural signatures of the Shannon surprise (and the state prediction error, c.f. Proposition 3) have been found in numerous EEG (Gijsen et al., 2021; Kolossa et al., 2015; Kopp & Lange, 2013; Mars et al., 2008; Meyniel et al., 2016; Modirshanechi et al., 2019; Mousavi et al., 2020), MEG (Maheu et al., 2019; Meyniel, 2020; Mousavi et al., 2020), and fMRI (Gläscher et al., 2010; Konovalov & Krajbich, 2018; Loued-Khenissi & Preuschoff, 2020) studies. In particular, it is well-known that the amplitude of the EEG P300 component correlates with the Shannon surprise (Kolossa et al., 2015; Kopp & Lange, 2013; Mars et al., 2008; Meyniel et al., 2016; Modirshanechi et al., 2019; Mousavi et al., 2020). Modirshanechi et al., 2019 showed that such correlations can be exploited to decode the Shannon surprise of stimuli from EEG signals independently of the stimulus modality. Maheu et al., 2019 showed that the MEG amplitudes at different time windows correlate with different values of the Shannon surprise calculated according to different inference styles. The expected (over observations) Shannon surprise (usually interpreted as uncertainty) has also been shown to have signatures in EEG (Kopp & Lange, 2013), MEG (Meyniel, 2020), and fMRI (Konovalov & Krajbich, 2018; Loued-Khenissi & Preuschoff, 2020).

The signatures of the Bayesian and the Confidence Corrected surprise have been compared to those of the Shannon surprise (Gijsen et al., 2021; Kolossa et al., 2015; Mars et al., 2008; Mousavi et al., 2020; Ostwald et al., 2012; Visalli et al., 2021). Whereas some studies have reported that the EEG and MEG amplitudes were always best explained by only one of these surprise measures (Mars et al., 2008; Mousavi et al., 2020; Ostwald et al., 2012), other studies have reported separate significant contributions of different surprise measures to explaining variations of EEG signals (Gijsen et al., 2021; Kolossa et al., 2015; Visalli et al., 2021). In particular, the amplitude of the EEG P300 component at the frontal and frontocentral electrodes (called P3a component) has been reported to be explained better by the Bayesian surprise than the Shannon surprise, and the amplitude of the EEG P300 component at the parietal electrodes (called P3b component) has been reported to be explained better by the Shannon surprise than the Bayesian surprise (Kolossa et al., 2015; Visalli et al., 2021). In addition to the Bayesian and the Shannon surprise, Kolossa et al., 2015 reported a significant positive correlation between the postdictive surprise and the EEG amplitudes in a later time window (i.e., the EEG slow wave component). Gijsen et al., 2021 focused on the early EEG components and reported strong evidence for the existence of separate signatures for all Shannon, Bayesian, and Confidence Corrected surprise.

The Bayesian surprise has been shown to attract human attention (Itti & Baldi, 2006, 2009) and drive exploration in reward-driven experiments (Horvath et al., 2021) and has also been used in computational models of human curiosity (Gottlieb & Oudeyer, 2018; Gottlieb et al., 2013; Oudeyer et al., 2016; Schmidhuber, 2010). As mentioned earlier, surprise measures similar to the Bayesian surprise and postdictive surprise can be defined also by measuring the change in the belief and the marginal probability, respectively, via distance or pseudo-distance measures different from the KL-divergence (c.f. Eq. 23, Eq. 24, Eq. 25, and Eq. 26) (Baldi, 2002). Findling et al., 2021 showed that introducing computational noise proportional to such a measure of surprise into the inference mechanism can explain human adaptive behavior in a wide range of experiments. Antony et al., 2021 showed that a similar measure of surprise plays a role in the segmentation of memory and correlates with pupil dilation and the activity of dopamine-related regions of the brain in fMRI. The influence of surprise on memory has also been studied in reward-driven experiments where it has been shown that high unsigned reward prediction errors increase the memorability of task-independent stimuli (Rouhani & Niv, 2021; Rouhani et al., 2018); in such settings, reward can be defined as a component of the observation *y*_*t*_, and the unsigned reward prediction error can be seen as the absolute error surprise (c.f. Eq. 19).

In volatile environments like the one in Fig. 1B, human behavior has been well explained by exact or approximate Bayesian inference (Behrens et al., 2007; Heilbron & Meyniel, 2019; Nassar et al., 2012; Nassar et al., 2010; Wilson et al., 2013), supporting the idea that the Bayes Factor surprise modulates the speed of learning (c.f. Proposition 1). Nassar et al., 2012 showed that the adaptation rate *γ*_*t*+1_ (c.f. Proposition 1) correlates with the changes in pupil diameter in a Gaussian task with abrupt changes. In a recent study, Xu et al., 2021 showed that the Bayes Factor surprise modulates human reinforcement learning in a volatile multi-step decision-making experiment and correlates with the EEG P300 amplitude at frontal electrodes. They showed that such a correlation is independent of the correlation of the reward prediction error and novelty with the EEG P300 amplitude (Xu et al., 2021). Although these observations support the computation and the use of the Bayes Factor surprise in the brain, the Bayes Factor surprise and the Shannon surprise are indistinguishable in the above experiments as they all used a flat distribution for the prior marginal probabilities (c.f. Corollary 2). Liakoni et al., 2021 proposed a modification of the Gaussian task of Nassar et al., 2012 where the Bayes Factor surprise and the Shannon surprise have different behaviors.

Most of these previous studies have focused on one measure of surprise and its role and signatures in behavioral and physiological measurements. The few examples that considered more than one surprise measure (Gijsen et al., 2021; Kolossa et al., 2015; Mars et al., 2008; Mousavi et al., 2020; Ostwald et al., 2012; Visalli et al., 2021) (we note that 3 out of 6 were published after 2020) have focused on model-selection methods to compare different models and did not look for fundamentally different predictions of these measures. However, as we will see in the next section, well-designed experiments allow us to dissociate different surprise measures based on qualitatively different predictions.

### Summary and intermediate discussion (i)

In a unified framework, we discussed 9 previously proposed measures of surprise: (1) the Bayes Factor surprise; (2) the Shannon surprise; (3) the State Prediction Error; (4) the Absolute and (5) the Squared error surprise; (6) the Bayesian surprise; (7) the Postdictive surprise; (8) the Confidence Corrected surprise; and (9) the Minimized Free Energy. We considered different ways to define some of these measures in volatile environments and, overall, analyzed 16 different definitions of surprise.

Based on how different definitions depend on the belief *π*^(*t*)^, we divided them into three groups of probabilistic mismatch (𝒮_BF_, 𝒮_Sh1_, 𝒮_Sh2_, 𝒮_SPE1_, and 𝒮_SPE2_), observation-mismatch (𝒮_Ab1_, 𝒮_Ab2_, 𝒮_Sq1_, and 𝒮_Sq2_), and belief-mismatch (𝒮_Ba1_, 𝒮_Ba2_, 𝒮_Po1_, 𝒮_Po2_, 𝒮_CC1_, 𝒮_CC2_, and *F*^*^) surprise measures (Fig. 3). We then showed how these measures relate to each other theoretically and, more importantly, under which conditions they are invertible functions of each other (i.e., they are indistinguishable – Fig. 2A); these links are summarized in Fig. 2B.

Importantly, our theoretical analysis provides foundations for biological mechanism of surprise computation and surprise-modulated learning (Berlemont & Nadal, 2021; Frémaux & Gerstner, 2016; Gerstner et al., 2018; Iigaya, 2016; Soltani & Izquierdo, 2019). For example, it has been argued that the computation of observation-mismatch surprise measures is biologically more plausible than more abstract measures such as Shannon surprise (Iigaya, 2016). Our results identify conditions under which observation-mismatch surprise measures behave identically as those probabilistic mismatch surprise measures that are optimal for adaptive learning (c.f. Fig. 2B, Proposition 1, and Corollary 1).

Measures of surprise in neuroscience have been previously divided into two categories (Faraji et al., 2018; Gijsen et al., 2021; Hurley et al., 2011): ‘puzzlement’ and ‘enlightenment’ surprise. Puzzlement surprise measures how puzzling a new observation is for an agent, whereas enlightenment surprise measures how much the new observation has enlightened the agent and changed its belief – a concept closely linked but not identical to the ‘Aha! moment’ (Dubey et al., 2021; Kounios & Beeman, 2009). The Bayesian and the Postdictive surprise can be categorized as enlightenment surprise since both quantify information gain (Fig. 4). Based on our theoretical analyses, however, we suggest to further divide measures of puzzlement surprise into 3 sub-categories (Fig. 4):

**Figure 4:**
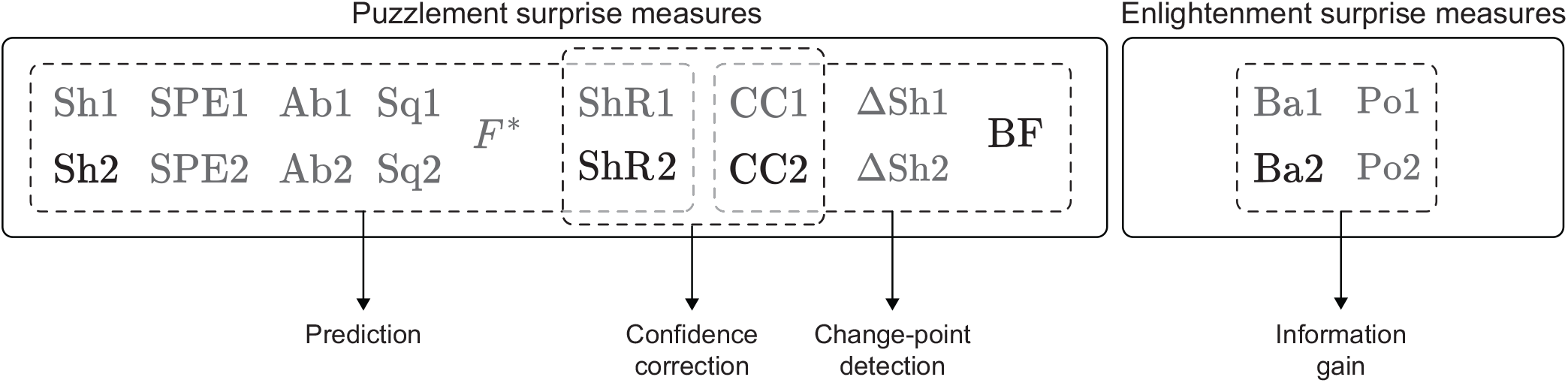
Conceptual categorization of surprise measures. Measures of puzzlement surprise (Faraji et al., 2018) can be further classified into 3 sub-categories of surprise measures highlighting (i) prediction, (ii) change-point detection, and (iii) confidence correction. According to surprise measures focused on prediction, the agent’s puzzle is finding the most accurate prediction of the next observation. According to surprise measures focused on change-point detection, the agent’s puzzle is to detect environmental changes. Surprise measures focused on confidence correction do not determine a specific puzzle for the agent but stress that confidence should explicitly influence puzzlement. The enlightenment surprise measures can be seen as measures of information gain. Black color shows the representative surprise measure of each category that we analyze in our case-studies. In addition to the 16 definitions of surprise discussed in section Theories of surprise: A technical review, we included in the figure the difference in Shannon surprise (ΔSh1 and ΔSh2) introduced in Proposition 2 and the regularized Shannon surprise (ShR1 and ShR2) introduced in section Regularized Shannon surprise: A new direction. Abbreviations: Ab: Absolute error surprise, Sq: Squared error surprise, BF: Bayes Factor surprise, Sh: Shannon surprise, SPE: State Prediction Error, Ba: Bayesian surprise, Po: Postdictive surprise, CC: Confidence Corrected surprise, *F*^*^: Minimized Free Energy, and ShR: Regularized Shannon surprise.

1. ‘Prediction surprise’: Surprise measures that quantify how unpredicted, unexpected, or unlikely the new observation is, including the Shannon surprise, State Prediction Error, the Minimized Free Energy, and all observation-mismatch surprise measures (Fig. 4). According to these measures, the agent’s puzzle is to find the most accurate predictions of the next observations.
2. ‘Change-point detection surprise’: Surprise measures designed to modulate the learning rate and to identify environmental changes, including the Bayes Factor surprise and the difference in Shannon surprise (c.f. Corollary 1; Fig. 4). According to these measures, the agent’s puzzle is to detect environmental changes.
3. ‘Confidence correction surprise’: Surprise measures that explicitly account for the agent’s confidence, including the Confidence Corrected surprise. The idea is that higher confidence (or higher commitment to a belief) leads to more puzzlement, where the puzzle is either to detect environmental changes or to find the most accurate prediction. We will introduce a new measure in this family in section Regularized Shannon surprise: A new direction that explicitly captures confidence and defines the agent’s puzzle as finding the most accurate predictions (Fig. 4).

Following this classification, we focus in the following section on one representative example from each of these sub-categories, and, whenever there are two definitions of one surprise measure, we take the second one: we include, in our analysis, the Shannon surprise 𝒮_Sh_ = 𝒮_Sh2_, the Confidence Corrected surprise 𝒮_CC_ = 𝒮_CC2_, the Bayesian surprise 𝒮_Ba_ = 𝒮_Ba2_, and the Bayes Factor surprise S_BF_ (colored in black in Fig. 4).

## Different definitions make different predictions: Case studies

In the previous section, we clustered definitions of surprise into 4 main categories based on important conceptual differences (Fig. 4). In this section, we design experimental paradigms where these different definitions make different predictions. Our goal is (i) to better understand the behavior of each measure, (ii) to examine whether they are consistent with common sense, and (iii) to aid future experimental studies to dissociate the contribution of different measures of surprise to explaining biological and behavioral observations. In particular, we are interested in the links of surprise to (1) perception, (2) learning, and (3) decision-making. For each of these links, we present one case-study (in total 3) where we present different thought experiments as well as propositions for actual experiments.

### Case-study 1: Confidence influences measures of surprise differently

In our first case-study, we address the following question: ‘how does an agent’s confidence in its belief and predictions influence its perception of surprise?’. To do so, we study two thought experiments and propose two actual experiments where different surprise measures make different predictions for the dependence of surprise upon confidence.

### General setting: Task formulation

All our experiments can be formulated in the form of a categorical task: Each observation *y*_*t*_ can be seen as a number between 1 and *N* sampled from a categorical distribution, e.g., the observations are generated by rolling a die, flipping a coin, or playing a random musical note. We assume that there is no cue variable *x*_*t*_ and no change in the environment (*p*_*c*_ = 0). In human experiments, these assumptions should be communicated to participants through instructions at the beginning of the experiment. Because there is no change and there are exactly *N* possibilities for *y*_*t*_, Θ = Θ_1_ = … = Θ_*t*_ is a vector of *N* probabilities [*P*_1_, …, *P*_*N*_] corresponding to different values of *y*_*t*_. Since there are no cues, the conditional distribution *P*_*Y* |*X*_ is specified as

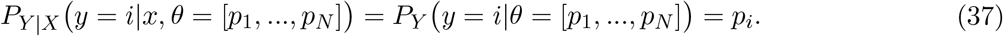

An example of a categorical task is the oddball paradigm (Squires et al., 1976). In a typical oddball task, participants are exposed to a sequence of binary stimuli (i.e., *N* = 2), e.g., they listen to an auditory sequence composed of two different musical notes, where one stimulus is less frequent than the other, e.g., *p*_1_ = 0.9 and *p*_2_ = 0.1. The frequent stimulus is often called ‘standard’, and the infrequent stimulus is often called ‘deviant’. Oddball tasks are among the most widely used paradigms for the study of the neural and behavioral signatures of surprise (Huettel et al., 2002; Maheu et al., 2019; Meyniel et al., 2016; Modirshanechi et al., 2019; Rubin et al., 2016; Squires et al., 1976). We will also consider generalized oddball task with *N* = 3 stimuli.

As a natural choice for a categorical task, we assume that the prior and the current beliefs are both Dirichlet distributions^2^

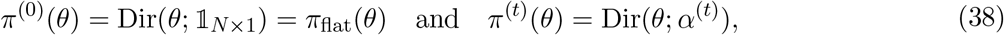

where 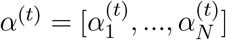 are the parameters of the Dirichlet distribution for the belief at time *t*, and the initial belief *π*^(0)^ is the uniform distribution, i.e., a Dirichlet distribution with parameters *α*^(0)^ = 𝕝._*N* ×1_. At any time-point *t* ≥ 0, we can write

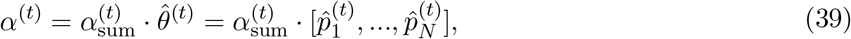

where we define 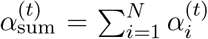 and 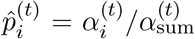. The vector 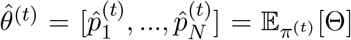 is an estimate of *θ* given the belief *π*^(*t*)^. We discuss the parameter 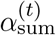 below when we define confidence. Using this notation, we can write the marginal probability as 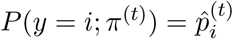. For example, with the belief *π*^(*t*)^ at time 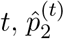 is the estimated occurrence probability of the second category (e.g., the deviant note in an auditory oddball task). For the initial belief *π*^(0)^, we have 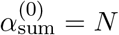 and 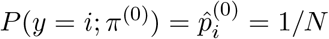 for all *i* between 1 and *N*.

Because *P* (*y*; *π*^(0)^) is a flat distribution, 𝒮_BF_ and 𝒮_Sh_ are invertible functions of each other and hence indistinguishable (Fig. 2). Therefore, in this section, we do not explicitly include 𝒮_BF_ in our comparisons as all qualitative results for 𝒮_Sh_ hold true also for 𝒮_BF_; in particular, if 𝒮_Sh_ increases or decreases, 𝒮_BF_ does as well. Moreover, in the setting described above, we have 𝒮_Sh1_ = 𝒮_Sh2_, 𝒮_Ba1_ = 𝒮_Ba2_, and 𝒮_CC1_ = 𝒮_CC2_ due to the assumptions *p*_*c*_ = 0 and *π* = *π*_flat_ (Fig. 2B).

### General setting: Confidence definition

To study the effect of confidence on the perception of surprise, we first need to agree how to measure confidence given a belief *π*^(*t*)^. In Eq. 29, we have defined the confidence given a belief *π*^(*t*)^ by the negative entropy *C*[*π*^(*t*)^] (Faraji et al., 2018). However, when *π*^(*t*)^ is a Dirichlet distribution, there is no analytic expression for *C*[*π*^(*t*)^]. For reasons of practicality, we therefore define here confidence in a categorical task as the inverse variance

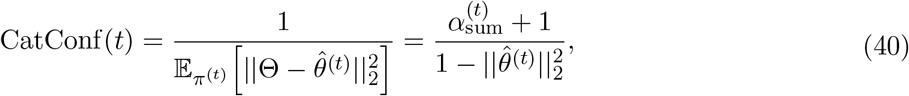

where CatConf stands for Categorical Confidence, 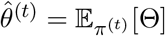 is the current estimate of the parameter, ||.||_2_ stands for the, *ℓ*_2_-norm, and 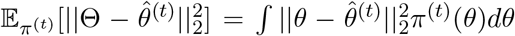 is the variance of Θ given the belief *π*^(*t*)3^. According to Eq. 40, for a fixed estimate 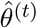, confidence is an increasing function of 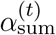 (c.f. Fig. 5A), and, for a fixed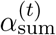, confidence is an increasing function of 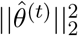 (c.f. Fig. 5B). The parameter 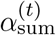 has been interpreted as the number of samples the belief *π*^(*t*)^ is worth (Efron & Hastie, 2016). We note that 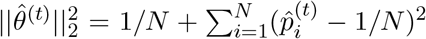 is a measure of the difference between the estimate 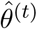 and the uniform distribution over *N* categories – i.e., 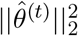 takes its maximum value (corresponding to maximum confidence) when the estimate 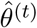 has a probability of 1 for one category and zero for the rest, and it takes its minimum value (corresponding to minimum confidence) when 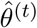 is distributed uniformly over all categories.

**Figure 5:**
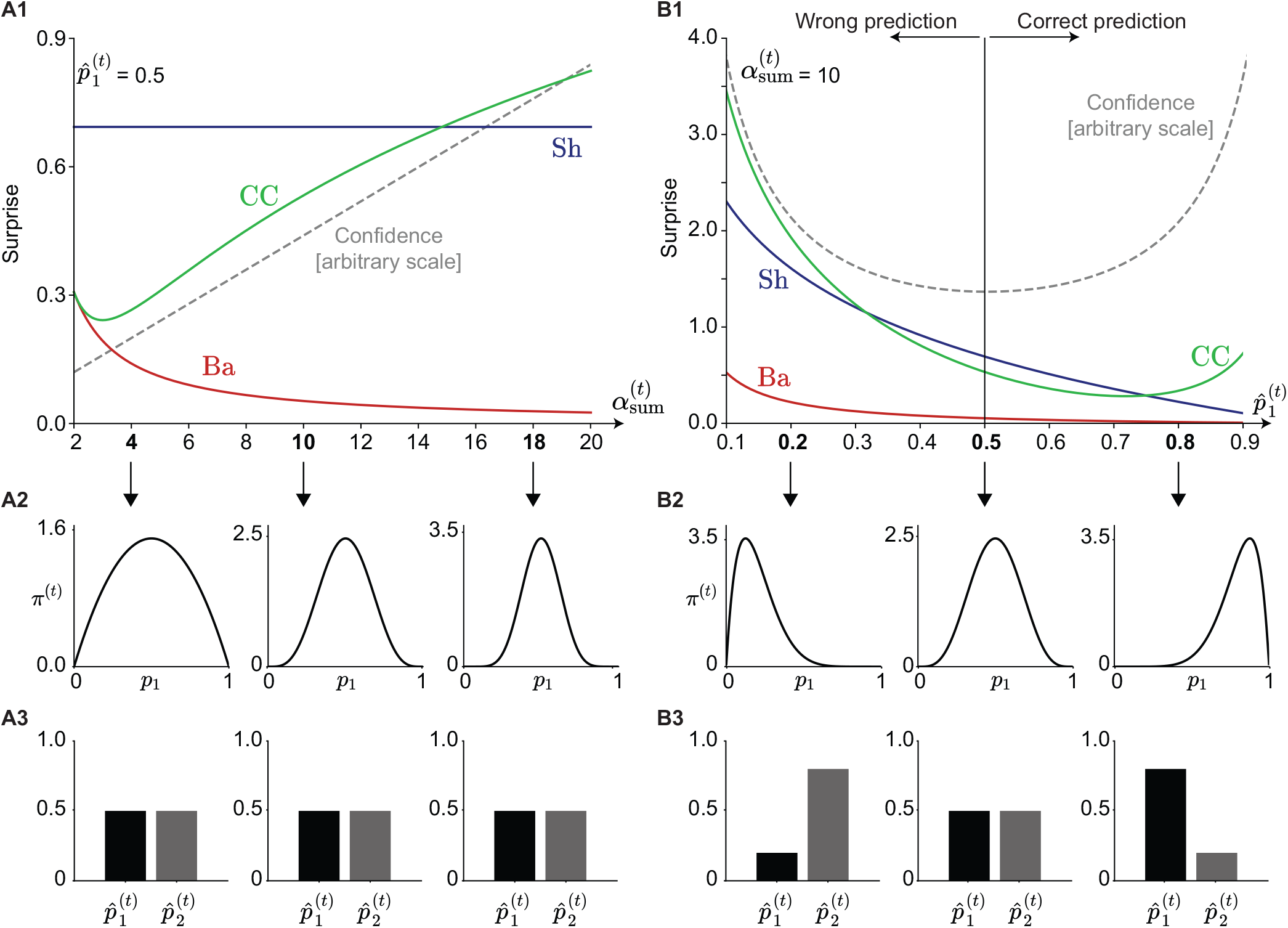
Coin flipping. **A**. Effect of 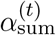 on surprise when 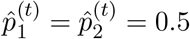 is fixed. **A1**. The Shannon (Sh, blue), the Bayesian (Ba, red), and the Confidence Corrected (CC, green) surprise for *Y*_*t*+1_ = 1 as functions of 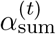. The dashed grey line shows the confidence as defined in Eq. 40 (on an arbitrary scale). **A2**. Three instances of the belief *π*^(*t*)^ with same expected value, corresponding to different values of 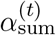 (indicated by arrows). **A3**. The estimate of the parameters 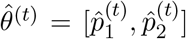 corresponding to the beliefs in panel A2. **B**. Effect of 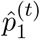 on surprise when 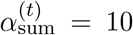 is fixed. **B1**. The Shannon (Sh, blue), the Bayesian (Ba, red), and the Confidence Corrected (CC, green) surprise for *Y*_*t*+1_ = 1 as functions of 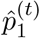. The dashed grey curve shows the confidence. We consider 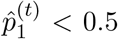 as an indication of *wrong* prediction of *Y*_*t*+1_ and 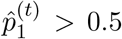 as an indication of *correct* prediction of *Y*_*t*+1_. **B2**. Three instances of the belief *π*^(*t*)^ corresponding to different values of 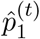 with same 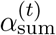. **B3**. The estimate of the parameters 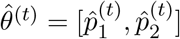 corresponding to the beliefs in panel B2. We note that the qualitative behavior of different measures with respect to 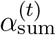 remains the same for other values of 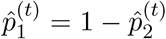, their qualitative behavior with respect to 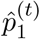 and remains the same for other values of 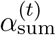.

As we show below, there is no general rule for the dependence of different surprise measures on confidence as defined in Eq. 40. Rather, depending on different values of 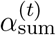 and 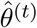, confidence affects surprise measures differently.

### Thought experiment: (i) Coin flipping

Imagine we flip a coin. How surprising is it to see a head? How does our surprise change depending on how long we have owned the coin? Or depending on how much we believe in its fairness?

As our first thought experiment, we consider a coin flipping experiment where the observation *Y*_*t*+1_ is either a head (*Y*_*t*+1_ = 1) or a tail (*Y*_*t*+1_ = 2) – i.e., *N* = 2. Our goal is to study the effect of confidence as defined in Eq. 40 on surprise of observing a head (*Y*_*t*+1_ = 1) depending on how often we have flipped the coin (i.e., how large 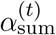 is) and depending on our estimate of its bias (i.e., how large 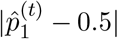 is).

We first fix 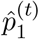 and study the behavior of different surprise measures as a function of the parameter 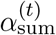.In other words, we assume that we have a fixed estimate of the coin’s bias (expressed by 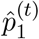), but our confidence about our estimation (measured by how often we have flipped it, 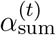) varies (grey, Fig. 5A1). We observe that the Shannon surprise 𝒮_Sh_ of *Yt*+1= 1 is independent of 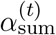 (blue, Fig. 5A1), as it is expected from a probabilistic mismatch surprise measure (Fig. 3). The Bayesian surprise 𝒮_Ba_, however, decreases with increasing 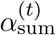 (red, Fig. 5A1). The reason is that the more confident we are about our belief, the less a single new observation changes our belief. In Appendix C: Methods for case-studies, we show that these results are not limited to the coin flipping experiment (*N* = 2) but are true for any categorical task with arbitrary *N*. The Confidence Corrected surprise S_CC_, on the other hand, does not change monotonically with respect to 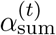 (green, Fig. 5A1): It is a decreasing function of confidence for small values of 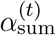 but an increasing function of confidence for large values of 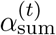.

Next we fix the parameter 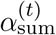 (i.e., how often we have flipped the coin) and study the behavior of different surprise measures as functions of our estimate of the coin’s bias 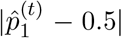. For *N* = 2, the term 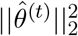 in Eq. 40 is equal to 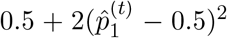, thus confidence is an increasing function of the estimated bias 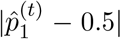 (grey, Fig. 5B1). The effect of confidence on surprise of observing *Y*_*t*+1_ = depends on whether the agent’s prediction has been wrong (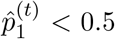, i.e., *Y*_*t*+1_ = 1 was observed even though it was thought to be less likely than *Y*_*t*+1_ = 2) or correct 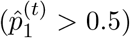; c.f. Fig. 5B1. For a wrong prediction, surprise increases with increasing confidence, i.e., with decreasing 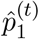 when 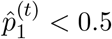, for all 3 surprise measures (Fig. 5B1, left). This means that all surprise measures predict that a wrong prediction made with higher confidence leads to higher surprise than a wrong prediction made with little confidence. For a correct prediction, higher confidence decreases Shannon 𝒮_Sh_ and Bayesian 𝒮_Ba_ surprise (blue and red, respectively, Fig. 5B1, right). In Appendix C: Methods for case-studies, we show that these results are not limited to the coin flipping experiment and are true for any categorical task. Interestingly, the Confidence Corrected surprise 𝒮_CC_ does not change monotonically with respect to the confidence for a correct prediction (green, Fig. 5B1, right). For 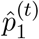 slightly greater than 0.5, it decreases with an increase in confidence, but then, for larger values of 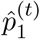, the slope turns and S_CC_ increases with an increase in confidence. In other words, 𝒮_CC_ predicts that the more we expect to observe a head (*Y*_*t*+1_ = 1), the more surprised we are upon observing it. This prediction looks counter-intuitive.

To summarize, the three paradigmatic measures of surprise show qualitative differences depending on 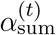 given a fixed estimate of the parameter 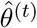. The Shannon surprise 𝒮_Sh_ stays constant, the Bayesian surprise 𝒮_Ba_ decreases, and the Confidence Corrected surprise 𝒮_CC_ first decreases and then increases. Similarly, they show qualitative differences depending on the estimate of the parameter 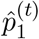 given a fixed 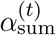.

### Thought experiment: (ii) CEO selection

Faraji et al., 2018 discussed a thought experiment where a few employees are waiting for the outcome of the next CEO selection. The idea is to analyze how surprise upon observing the outcome changes as a function of employees’ confidence in their beliefs. We model the example of Faraji et al., 2018 as a categorical task with a Dirichlet belief.

We assume there are *N* = 3 candidates for the next CEO and model an employee’s expectation about the outcome as a Dirichlet belief parameterized by 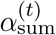 and 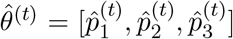. All employees have spent the same number of years in the company so that we can assume that 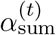 has the same value for all of them. We analyze the dependency of the surprise felt when the 1st candidate is selected (i.e., *Y*_*t*+1_ = 1) as a function of 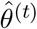. In particular, we follow Faraji et al., 2018 and assume that 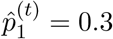 is fixed among all employees, i.e., they all estimate a probability of 0.3 for the selection of the 1st candidate, but their estimated selection probabilities for the other candidates vary. Faraji et al., 2018 argued that an employee who puts a probability of 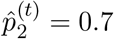 to the selection of the 2nd candidate 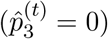 is more surprised when the 1st candidate is selected than another employee who considers the chances for candidates 2 and 3 to be identical 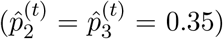 and close to that of the 1st candidate 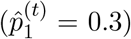 We therefore look at the surprise of the 1st candidate being selected (i.e., *Y*_*t*+1_ = 1) as a function of the employees’ expectation 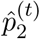 for the 2nd candidate; note that 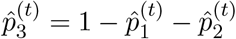.

The Shannon surprise 𝒮_Sh_ and the Bayesian surprise 𝒮_Ba_ are independent of 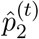, which implies that the surprise of observing *Y*_*t*+1_ = 1 is independent of how estimations 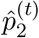 and 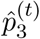 ^*t*)^ are distributed across the other candidates (blue and red, Fig. 6A1) – in Appendix C: Methods for case-studies, we show that this result is not limited to the CEO selection experiment but is true for any categorical task. The Confidence Corrected surprise S_CC_, however, has an interesting U-shape relation with 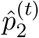 (green, Fig. 6A1) – similar to the dependence of confidence on 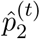 (grey, Fig. 6A1). The minimum surprise corresponds to 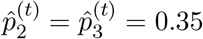 where confidence goes through its minimum and an employee’s estimate of the outcome probability is as uninformative as possible (Fig. 6A). This means that, given the same expectation for the selection of the 1st candidate, the employees who have a high expectation for the selection of the 2nd or the 3rd candidate are going to be more surprised when hearing that the 1st candidate was selected than the employees who consider all three candidates about equally likely to be selected. This behavior looks plausible and is exactly what 𝒮_CC_ was designed to capture (Faraji et al., 2018). 𝒮_CC_, therefore, explains aspects of surprise perception that are not captured by 𝒮_Sh_ and 𝒮_Ba_.

**Figure 6:**
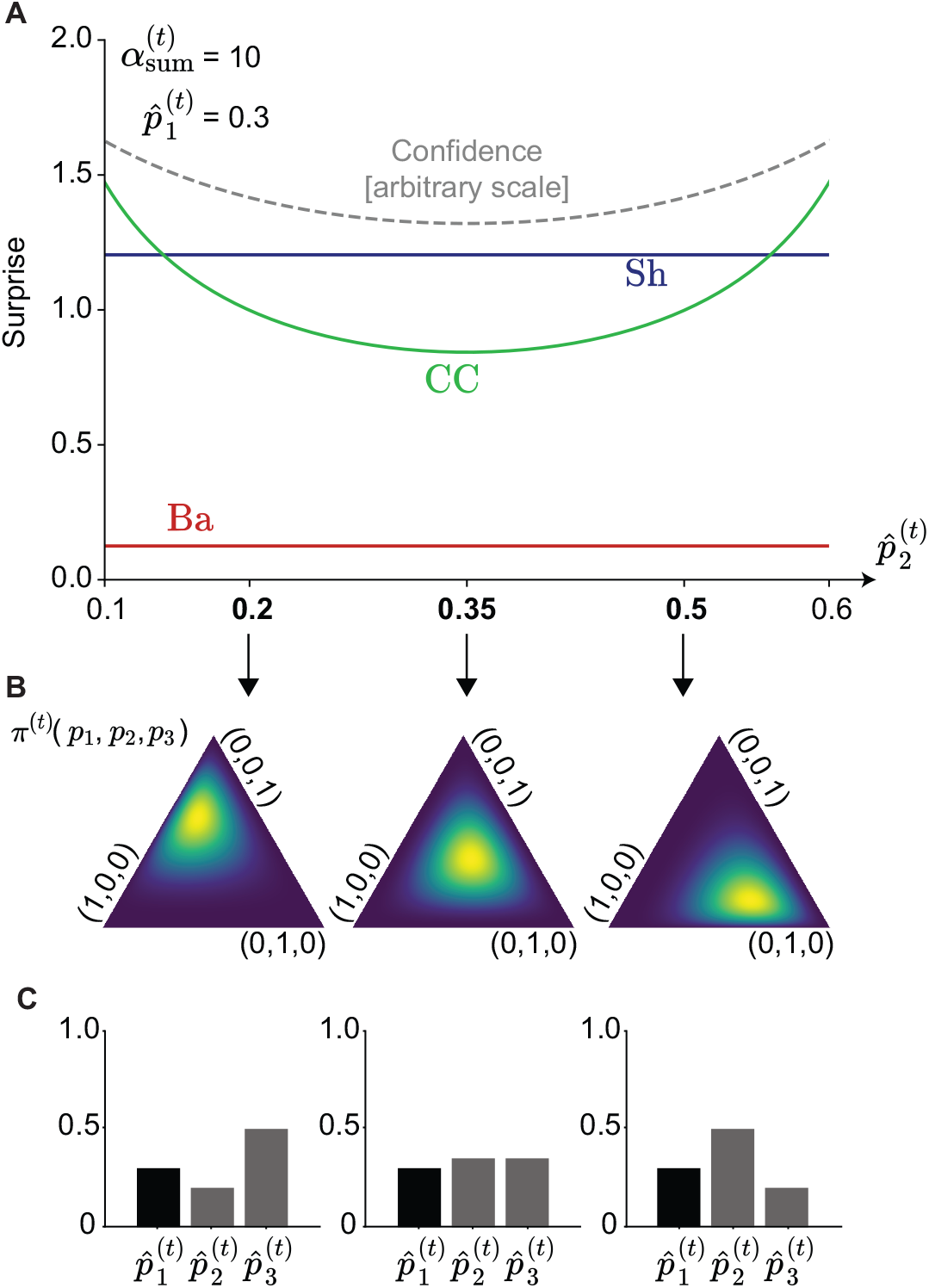
CEO selection. **A**. The Shannon (Sh, blue), the Bayesian (Ba, red), and the Confidence Corrected (CC, green) surprise for *Y* = 1 as functions of 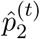 (with 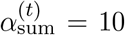 and 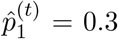 fixed).The dashed grey curves show the confidence as defined in Eq. 40 (on an arbitrary scale). **B**. Three instances of the belief *π*^(*t*)^ corresponding to different values of 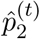 in panel A. Note that the space is shown as a triangle because *θ* = [*p*_1_,*p*_2_,*p*_3_] lives inside the area specified by *p*_1_ + *p*_2_ + *p*_3_ = 1, *p*_1_ ≥ 0, *p*_2_ ≥ 0, and *p*_3_ ≥ 0. **C**. The estimate of the parameters 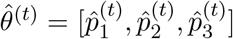 corresponding to the beliefs in panels A. We note that the qualitative behavior of different measures with respect to 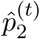 remains the same for other choices of 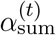 and 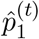.

To summarize, 𝒮_CC_ can capture aspects of surprise perception that are consistent with our intuition and are not captured by 𝒮_Sh_ and 𝒮_Ba_. However, 𝒮_CC_ has also some non-intuitive behavior that appears to contradict common sense – as observed in the coin flipping experiment. Our results suggest that a new definition of surprise may be needed for capturing the intuitively correct behavior in both thought experiments (Fig. 5 and Fig. 6). We address this problem in section Regularized Shannon surprise: A new direction.

### Proposed experiment: (i) Classic oddball task (*N* = 2)

The different predictions of different surprise measures with respect to changes in 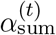 (for a fixed 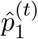, Fig. 5A) and 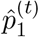 (for a fixed 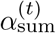, Fig. 5B) in the coin flipping experiment can be exploited in a classic oddball task (*N* = 2) that aims at determining which measure of surprise correlates best with a behavioral or physiological measurement *Z* (Fig. 2A1). In particular, our analyses (Fig. 5) suggest that different surprise measures have a different behavior at the beginning (early phase, Fig. 7A) and at the end (late phase, Fig. 7A) of an oddball task. First, as time passes, participants receive more samples to make their estimations, i.e., 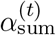 moves towards the right side of Fig. 5A1. Second, with time they learn that the standard stimuli are more likely to happen, i.e., 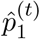 moves towards the right side of Fig. 5B1. Therefore, according to the Shannon and the Bayesian surprise, we expect the standard stimuli to be more surprising during the early phase ofthe experiment than the late phase (Fig. 7B). The opposite holds for the Confidence Corrected surprise (Fig. 7B). Moreover, the Shannon and the Confidence Corrected surprise predict, as a function of time, an increase in the surprise upon observing a deviant stimulus, whereas the Bayesian surprise predicts the opposite (Fig. 7C). Therefore, we find a double dissociation. Thus, depending on how the measurement *Z* behaves in the two cases (Fig. 7B and Fig. 7C), we can uniquely determine with which of the three surprise measures it correlates. As a proof of concept, we analyzed a publicly available visual oddball dataset (Robbins et al., 2018) (18 participants) with the EEG recordings at central electrodes, i.e., Cz, C1, and C2. We find that the EEG amplitude at around 450ms after the stimulus onset has a behavior consistent with the predictions of 𝒮_Sh_ but not with those of 𝒮_Ba_ or 𝒮_CC_. See Appendix D: EEG analysis and Fig. 7D-F for details.

**Figure 7:**
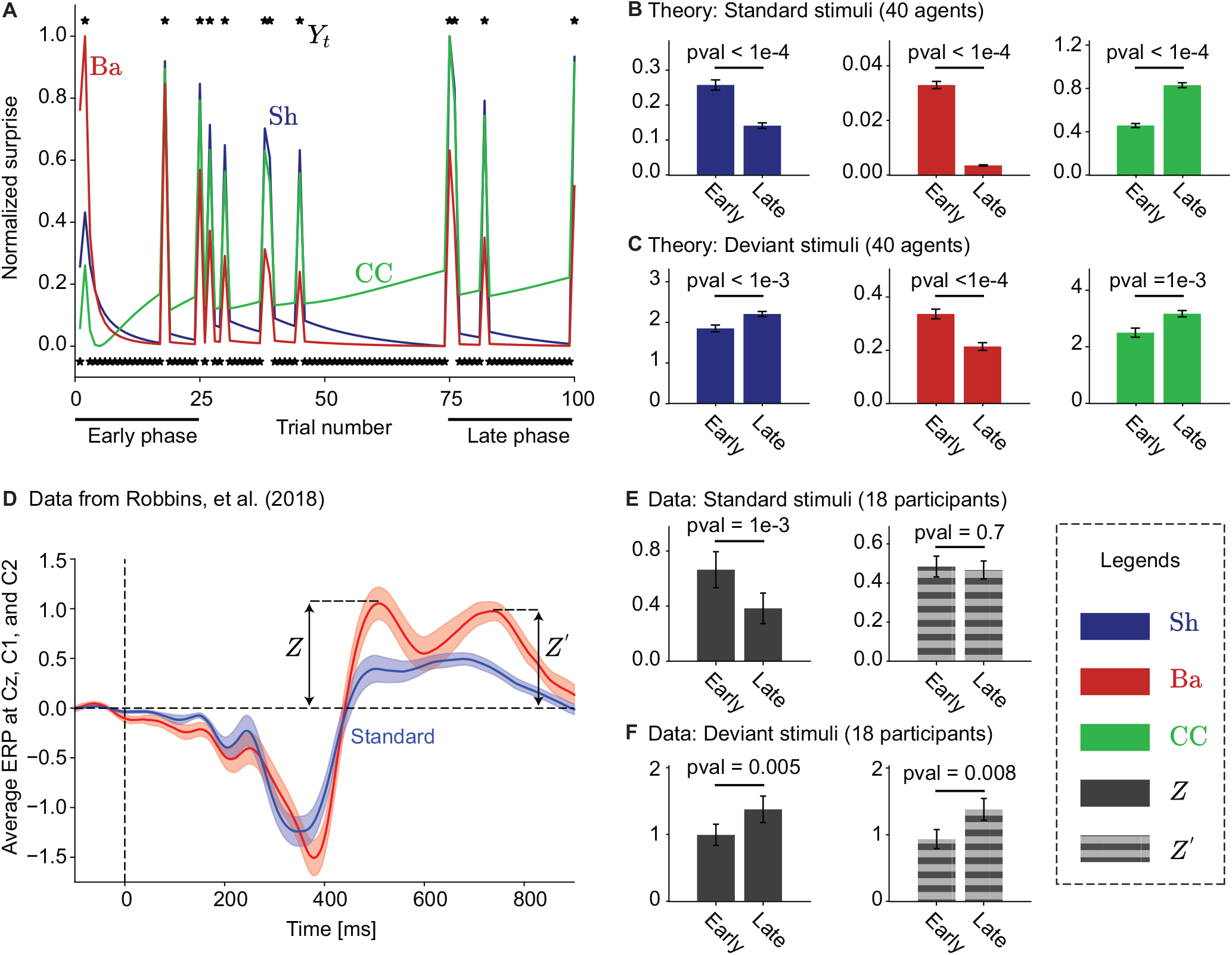
Different measures of surprise have different predictions for the early and the late phase of oddball tasks. **A**. The Shannon (Sh, blue), the Bayesian (Ba, red), and the Confidence Corrected (CC, green) surprise over time for a sequence of 100 binary stimuli (0.1 probability for the deviant stimulus); see Appendix C: Methods for case-studies for simulation details. The surprise sequences were linearly scaled between 0 and 1 before plotting. Each star shows one observation *Y*_*t*_; the standard (*Y*_*t*_ = 1) is plotted at −0.05, and the deviant (*Y*_*t*_ = 2) is plotted at 1.05. The early phase consists of the first 25 stimuli and the late phase of the last 25 stimuli. **B**. Average (over 40 random seeds) surprise values of standard stimuli in the early (first 25 out of 100 trials) and the late (last 25 out of 100 trials) phase of the task. **C**. Average (over 40 random seeds) surprise values of deviant stimuli in the early and the late phase of the task. **D**. Event related potential (ERP) averaged over central electrodes (Cz, C1, and C2, averaged over 18 participants after normalization) – raw dataset from Robbins et al., 2018 (see Appendix D: EEG analysis). The red curve corresponds to the deviant ERP and the blue curve to the standard ERP. *Z* and *Z*^′^ highlight two different physiological candidates for surprise values. Shaded areas show the standard error of the mean (over 18 participants). **E**. Average (over 18 participants) *Z* and *Z*^′^ for the standard stimuli in the early (first 75 out of ∼270 trials) and the late (last 75 out of ∼270 trials) phase of the task in Robbins et al., 2018. **F**. Average (over 18 participants) *Z* and *Z* ^*′*^for the deviant stimuli in the early and the late phase of the task in Robbins et al., 2018. In all panels, error bars show the standard error of the mean. Overall, *Z* matches the prediction of the Shannon surprise, and *Z*^′^ can be sensitive to both the Shannon and the Confidence Corrected surprise but not the Bayesian surprise; see Appendix D: EEG analysis for details.

Since our proposed approach relies on qualitative differences (such as increase or decrease) of surprise measures, it goes beyond general methods of naive model selection (Gijsen et al., 2021; Kolossa et al., 2015; Konovalov & Krajbich, 2018; Kopp & Lange, 2013; Maheu et al., 2019; Mars et al., 2008; Meyniel et al., 2016; Modirshanechi et al., 2019; Mousavi et al., 2020; Ostwald et al., 2012; Visalli et al., 2021) and enables us to reject different theories in a clear and reliable manner (c.f. Nassar and Frank, 2016). This opens the door to more theory-driven and principled approaches for future experimental studies.

### Proposed experiment: (ii) Generalized oddball task (*N* = 3)

The U-shape relation between the Confidence Corrected surprise and 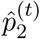 in the CEO selection experiment (Fig. 6) can be tested in a generalized oddball task with *N* = 3 stimuli (similar to Mars et al., 2008). The approach is similar to that of the binary oddball task in Fig. 7: We have a physiological measurement *Z* that is thought to be sensitive to surprise; we want to test whether its behavior is consistent with the predictions of 𝒮_CC_ or with those of 𝒮_Sh_ and 𝒮_Ba_.

To do so, we design a sequence of stimuli consisting of two phases separated by an abrupt change (Fig. 8A). The idea is to keep the occurrence frequency of the 1st stimulus *p*_1_ fixed throughout the whole sequence and change the balance between *p*_2_ and *p*_3_ from Phase 1 to Phase 2 (Fig. 8B). In particular, we consider the case that *p*_2_ = *p*_3_ in Phase 1 (before the change) while *p*_2_ *≪ p*_3_ in Phase 2 (after the change). In this case, 𝒮_Sh_ and 𝒮_Ba_ predict that the surprise of observing *Y*_*t*_ = 1 is the same in Phase 1 as in Phase 2, whereas 𝒮_CC_ predicts that observing *Y*_*t*_ = 1 is more surprising in Phase 2 than in Phase 1 (Fig. 8C). Such an experiment will show whether a given physiological indicator of surprise is influenced by confidence or not.

**Figure 8:**
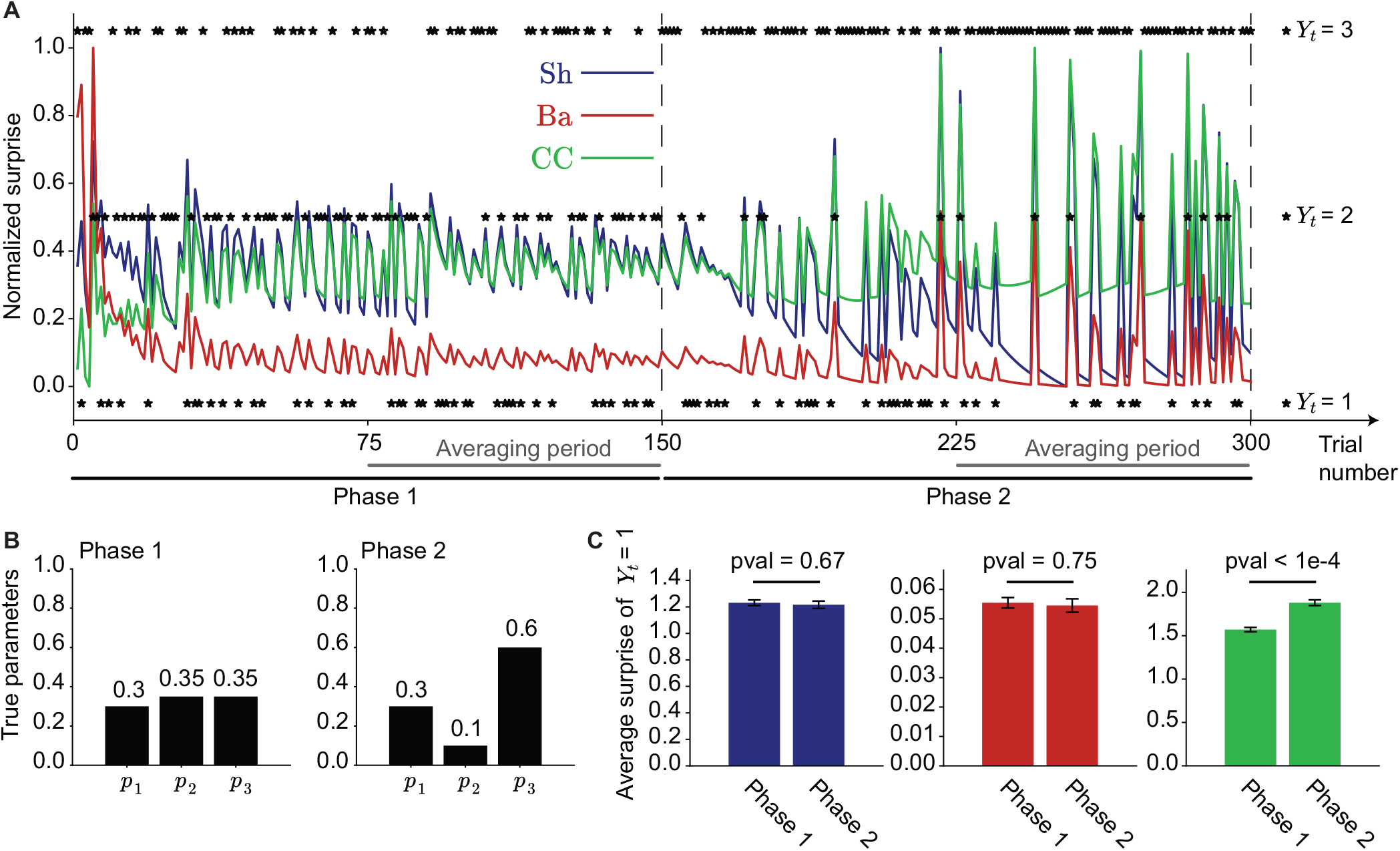
Testing the prediction of the CEO selection experiment in a generalized oddball task with *N* = 3 categorical stimuli. **A**. The Shannon (Sh, blue), the Bayesian (Ba, red), and the Confidence Corrected (CC, green) surprise over time for a sequence of 300 stimuli; see Appendix C: Methods for case-studies for simulation details. Each star shows one observation *Y*_*t*_ (right vertical axis), and the surprise sequences were linearly scaled between 0 and 1 before plotting. There is an abrupt change in the true distribution of the observations at *t* = 151 dividing the sequence into two phases, i.e., ‘Phase 1’ (1 ≤ *t* ≤ 150) and ‘Phase 2’ (151 ≤ *t* ≤ 300). The probability of *Y*_*t*_ = 1 remains the same through the whole sequence (*P*_1_ = 0.3), but *Y*_*t*_ = 2 becomes significantly less likely after the change-point (see panel B). To avoid complications concerning how participants update their belief, we focus on the 2nd half of each phase (indicated by the grey line and called ‘Averaging period’) where they have a relatively accurate estimate of probabilities. **B**. The true underlying distribution in Phase 1 (left) and Phase 2 (right). **C**. Average (over 40 random seeds) surprise values of *Y*_*t*_ = 1 in Phase 1 (trials 76 to 150) and Phase 2 (trials 226 to 300).

Interestingly, SanMiguel et al., 2021 have recently conducted a very similar experiment, parallel to our study and unknown to us. In a generalized oddball task with *N* = 11 stimuli, they observed that increasing *p*_2_ from 1*/*11 to 10*/*11 (while keeping *p*_1_ = 1*/*11 fixed) increases an indicator of surprise in EEG (the Mismatch Negativity amplitude; c.f. Näätänen et al., 2007) in response to *Y*_*t*_ = 1. This observation matches the predictions of 𝒮_CC_ but not those of S_Sh_ and 𝒮_Ba_.

### Case-study 2: Learning in volatile environments

In our second case-study, we investigate the link between surprise and learning. According to previous theoretical (Faraji et al., 2018; Findling et al., 2021; Frémaux & Gerstner, 2016; Gerstner et al., 2018; Liakoni et al., 2021; Yu & Dayan, 2005) and experimental (Behrens et al., 2007; Findling et al., 2021; Heilbron & Meyniel, 2019; Nassar et al., 2012; Nassar et al., 2010; Soltani & Izquierdo, 2019; Xu et al., 2021) studies, surprising events modulate the speed of learning in the brain. The argument is that surprising events are caused by inadequate expectations. Hence, surprise indicates the need for updating the source of those expectations (i.e., the belief).

In volatile environments similar to our generative model (Fig. 1A), an unexpected event can occur either due to an abrupt change in the parameters of the environment (also called unexpected uncertainty (Soltani & Izquierdo, 2019; Yu & Dayan, 2005)) or due to the stochasticity of the environment (i.e., pure randomness, also called expected uncertainty (Soltani & Izquierdo, 2019; Yu & Dayan, 2005)). While in the former case forgetting the old observations and increasing the speed of learning are necessary to explain the rapid adaptive behavior observed in humans (Behrens et al., 2007; Nassar et al., 2012; Nassar et al., 2010; Xu et al., 2021), in the latter case, forgetting the earlier observations leads to unnecessary loss of information. Therefore, not all unexpected events should change the speed of learning (Liakoni et al., 2021; Soltani & Izquierdo, 2019; Yu & Dayan, 2005).

Here, we compare different surprise measures based on how informative they are about environmental changes. Our goal is to determine which measures are most useful to modulate the speed of learning. To do so, we study a thought experiment and propose an actual experiment where different surprise measures make different predictions on how an abrupt change in the environment influences surprise of upcoming observations.

### General setting: Task formulation

Both our experiments can be formulated in the same paradigm with a volatile environment (*p*_*c*_ *>* 0) and binary observations (*Y*_*t*_ ∈ {1, 2}). At time *t*, an oracle announces the cue *x*_*t*_ ∈ [0, 1] as a prediction of the probability of the event *Y*_*t*_ = 1 (e.g., a cursor is located on an axis between 0 and 1 on a computer screen). A hidden parameter indicates whether the probability predicted by the oracle is correct (Θ_*t*_ = 1) or whether the oracle’s prediction is uninformative (Θ_*t*_ = 0) and *Y*_*t*_ comes from a uniform distribution. Formally, we write

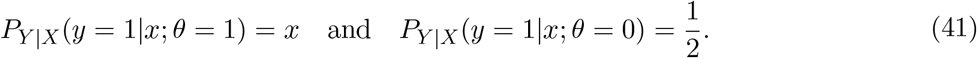

The belief *π*^(*t*)^(*θ* = 1) = *b*^(*t*)^ ∈ [0, 1] shows how much an agent has trust in the oracle, whereas *π*^(*t*)^(*θ* = 0) = 1 − *b*^(*t*)^ shows how much it believes in the unpredictability of the outcome.

### Thought experiment: Belief in Forecasts

We started the introduction of the paper by discussing an example of a wrong, and potentially surprising, weather forecast. A farmer who believed in the weather forecast and planned to outdoor work on a sunny morning will be surprised (and possibly angry) when he finds bad weather in the morning. Another farmer who thinks that weather forecasts are uninformative anyway will be less surprised than the first farmer.

Similarly, some people believe in opinion polls or predictions of election outcomes in the media, while others do not. Imagine two citizens A and B who have been living for many years in a country where each year an important election takes place between a blue party (*y*_*t*+1_ = 1) and a red party (*y*_*t*+1_ = 2). Each year the media announce, a week before the election, the probability *x*_*t*+1_ for the blue party to win (*Y*_*t*+1_ = 1). The media base their predictions on all available cues, including opinion polls, extrapolations from previous years, and elaborate mathematical models; however, nobody knows how reliable these predictions really are. Neither citizen A nor B has other independent cues. The citizens’ trust that the media are correct (Θ_*t*_ = 1) is represented by *b*^(*t*)^ which indicates how much they believe that the election is predictable and in agreement with the forecast of media. A trust of *b*^(*t*)^ = 0 implies that a citizen believes that the outcome is unpredictable and both parties are equally likely to win. Hence, given the forecast *x*_*t*+1_ by the media, a citizen with the belief *b*^(*t*)^ estimates the probability of the blue party to win as

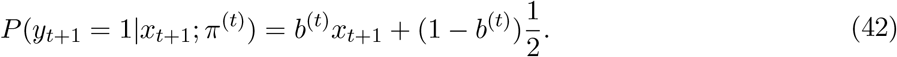

Importantly, the belief *b*^(*t*)^ of a citizen is not fixed but evolves with experience. It could be that for the first 20 years, the media did not know how to interpret data, relied on biased opinion polls, or used a bad mathematical model. Unknown to the citizens, starting in year 21, they may have a change in personnel and thereafter use a near-perfect mathematical model. Our claim, based on the results of earlier studies (Behrens et al., 2007; Faraji et al., 2018; Findling et al., 2021; Frémaux & Gerstner, 2016; Gerstner et al., 2018; Heilbron & Meyniel, 2019; Liakoni et al., 2021; Nassar et al., 2012; Nassar et al., 2010; Soltani & Izquierdo, 2019; Xu et al., 2021; Yu & Dayan, 2005), is that surprise such as ‘oh, this year the prediction was correct!’ is used by citizens to change their belief *b*^(*t*)^ in the trustworthiness of the forecast. Our goal is to find which surprise measure is most useful to indicate the need for such a change.

To do so, let us suppose that citizen A has a high trust (e.g., *b*^(*t*)^ = 0.9) in the forecast and citizen B a low trust (e.g., *b*^(*t*)^ = 0.05). We assume that the media predicted a 90-percent probability of the blue party to win (*x*_*t*+1_ = 0.9) and consider the amount of surprise of citizen A and B at the moment when the results of the election are announced. Under the hypothesis that surprise about the election outcome is used to change the belief, we expect that the following should occur:

- Expectation 1: Citizen A who has a strong trust in the media (e.g., *b*^(*t*)^ = 0.9) is more surprised if the actual outcome of the election proves the media wrong (red party wins) than if the election’s outcome (blue party wins) is consistent with the prediction of the media.
- Expectation 2: Citizen B who believes in unpredictability of the election (e.g., *b*^(*t*)^ = 0.05) is more surprised if the actual outcome of the election proves the media right (blue party wins as predicted) than if the the prediction is wrong.
- Expectation 3: If the outcome of the election is against the prediction of the media, then citizen A is more surprised than citizen B.

Note that, in the first two points, we compare the surprise value of two different outcomes for the *same* belief, whereas, in the last point, we compare the surprise value as a function of the belief *b*^(*t*)^.

For the case of citizen A (strong trust in the media) all definitions of surprise match our Expectation 1 (compare the blue and the red curves in Fig. 9 for high values of *b*^(*t*)^). If we follow the red curve from high belief to very low belief (i.e., if we decrease *b*^(*t*)^), then different measures of surprise show different behaviors. As *b*^(*t*)^ decreases, only the Bayes Factor surprise 𝒮_BF_ (Fig. 9A1) and the Shannon surprise 𝒮_Sh_ (Fig. 9A2) decrease and match our Expectation 3. Importantly, the red and blue curves cross for 𝒮_BF_ but not for 𝒮_Sh_. Hence, after a certain point (*b*^(*t*)^ *< b*^(0)^), 𝒮_BF_ predicts that the media being right is more surprising than the media being wrong – matching our Expectation 2. However, 𝒮_Sh_ has a different behavior: as *b*^(*t*)^ decreases, the surprise of the media being wrong gets closer to the surprise of the media being right, i.e., the more citizens believe in randomness, the smaller the difference in how surprised they are by either event. In other words, 𝒮_Sh_ indicates that when citizens believe in the unpredictability of the outcome, they do not care about the prediction of the media, and as a result, the prediction of the media does not affect their perception of surprise. Therefore, 𝒮_BF_ matches all our three Expectations, but 𝒮_Sh_ matches only Expectations 1 and 3.

**Figure 9:**
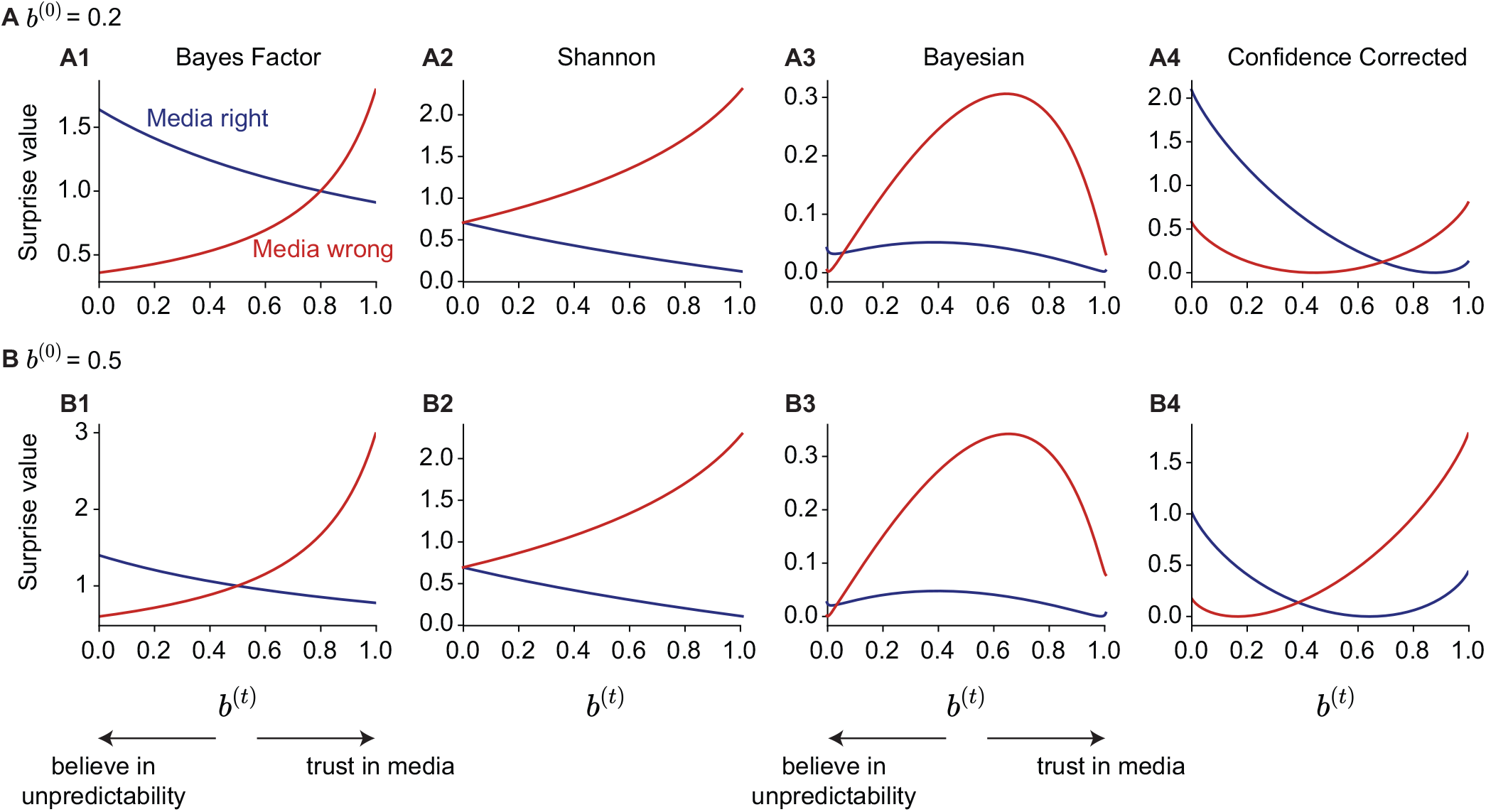
Surprise upon election of red or blue party as a function of trust *b*^(*t*)^ in the media. From left to right: Bayes Factor, Shannon, Bayesian, and Confidence Corrected surprise as functions of *b*^(*t*)^, for different prior belief: **A**. *b*^(0)^ = 0.2 and **B**. *b*^(0)^ = 0.5. The prediction of the media for the blue party to win is the same for all panels and equal to *X*_*t*_ = 0.9. The red curves correspond to the red party winning the election (i.e., *Y*_*t*_ = 2 and the media being wrong), and the blue curves correspond to the blue party winning the election (i.e., *Y*_*t*_ = 1 and the media being right). The change-point probability *p*_*c*_ is assumed to be 0.03. However, 𝒮_BF_, 𝒮_Sh_, and 𝒮_CC_ are independent of *p*_*c*_. The qualitative behavior of 𝒮_Ba_ remains the same for other values of *p*_*c*_ up to 0.2 (i.e., when there is on average one change-point per 5 time-points), but it becomes similar to the one of 𝒮_CC_ for the more volatile environments (i.e., for higher values of *p*_*c*_; c.f. Corollary 3). For all values of *p*_*c*_, the Bayesian surprise 𝒮_Ba1_ (equal to 𝒮_Ba2_ for *p*_*c*_ = 0; c.f. Eq. 23 and Eq. 24) has the same qualitative behavior as 𝒮_Ba2_ shown in panel A3 and B3.

Both 𝒮_Ba_ (Fig. 9A3) and 𝒮_CC_ (Fig. 9A4) are qualitatively distinguishable from 𝒮_BF_ and 𝒮_Sh_, since neither 𝒮_Ba_ nor 𝒮_CC_ has a monotonic relation with the degree of trust in media *b*^(*t*)^. Therefore, neither 𝒮_Ba_ nor 𝒮_CC_ matches our Expectation 3.

To summarize, different surprise measures have qualitatively different predictions for the surprise after an election outcome as a function of the trust *b*^(*t*)^ in the media. Importantly, only 𝒮_BF_ matches our expectations for a surprise measure that is useful for modulation of the speed of learning (c.f. Proposition 1). The results remain the same if the prior belief *b*^(0)^ has a different value (Fig. 9B).

### Proposed experiment: Association learning task

In this section, we propose an actual experiment to test predictions of the different surprise measures analogously to our thought experiment for belief in forecasts. Our proposed experiment is similar to the ones used in studies of association learning (Gershman et al., 2017; Niv et al., 2015): Participants should infer whether the predictions of an oracle *x*_1:*t*_ (e.g., cues or conditioned stimuli) are or are not associated with the observations *y*_1:*t*_. An alternative would be the design of a go no-go paradigm (Mars et al., 2008; Walz et al., 2015) with a cue variable *x*_*t*_ preceding each stimulus. In both types of experiment, we allow for associations to change abruptly from time to time.

In order to formalize experimental predictions, we consider a task with 150 trials (Fig. 10). At each time *t*, the oracle announces either *X*_*t*_ = 0.9 or *X*_*t*_ = 0.1, randomly chosen. For the first 50 trials, we assume that the observations are independent of the oracle’s prediction, i.e., Θ_*t*_ = 0 for 1 ≤ *t* ≤ 50 and *Y*_*t*_ ∼ Bernoulli(0.5). Then, unknown to the participants, there is an abrupt change at trial 51, and, forthe next 50 trials, the observations follow the same distribution as the one predicted by the oracle, i.e., Θ_*t*_ = 1 for 51 ≤ *t* ≤ 100 and *Y*_*t*_ ∼ Bernoulli(*X*_*t*_). Finally after another abrupt change at trial 101, the observations become again independent of the oracle’s predictions, i.e., Θ_*t*_ = 0 for 101 ≤ *t* ≤ 150 and *Y*_*t*_ ∼ Bernoulli(0.5). We study the behavior of different surprise measures for both cases of transition from an unpredictable environment to a predictable one (at *t* = 51) and vice versa (at *t* = 101). To model the temporal dynamics of belief in such a volatile environment, we use exact Bayesian inference as in Proposition 1 with change-point probability *p*_*c*_ = 0.03 – see Appendix C: Methods for case-studies for details.

**Figure 10:**
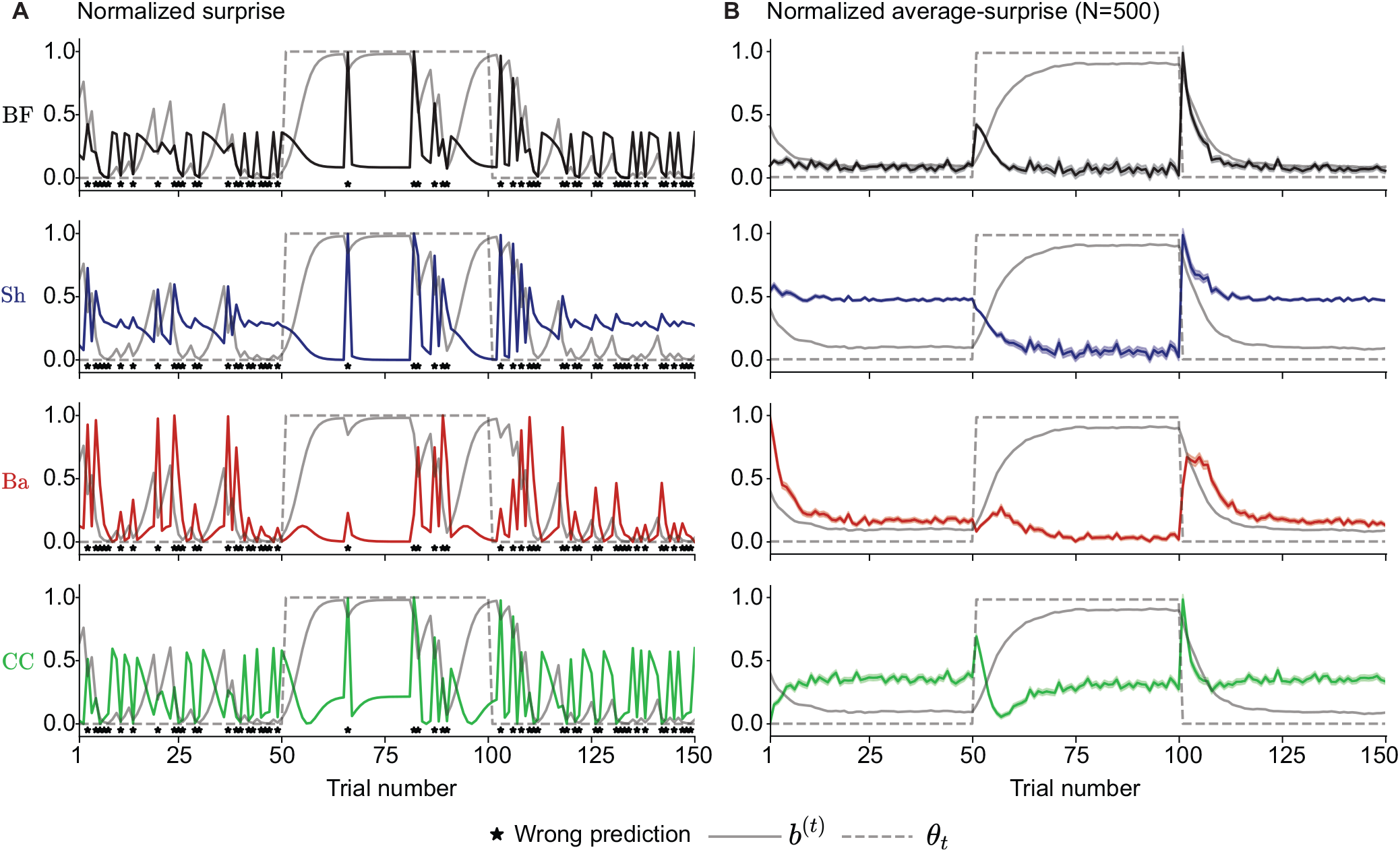
Association learning task. **A**. The Bayes Factor (BF, black), the Shannon (Sh, blue), the Bayesian (Ba, red), and the Confidence Corrected (CC, green) surprise over time. The exact same experimental sequence was used for all four surprise measures. The dashed grey curve shows Θ_*t*_, and the solid grey curve shows *b*^(*t*)^ over time. The oracle’s predictions at time *t* are either *X*_*t*_ = 0.9 or *X*_*t*_ = 0.1. The stars show the time-points when the oracle’s predictions were wrong. **B**. The averaged version of panel A (over 500 random seeds). The shaded areas indicate the standard error of the mean. The surprise sequences were linearly normalized between 0 and 1 before plotting in both panels. The change-point probability *p*_*c*_ is assumed to be 0.03. Note that the dependence of different surprise measures on *p*_*c*_ is mainly through its influence on the dynamics of belief updating, in contrast to Fig. 9 where the belief *b*^(*t*)^ is assumed to be fixed – see Appendix C: Methods for case-studies for details. The qualitative behavior of 𝒮_BF_ and 𝒮_Sh_ remains the same for other values of *p*_*c*_. The qualitative behavior of 𝒮_Ba_ and 𝒮_CC_ also remains the same for other values of *p*_*c*_ up to 0.2 (i.e., when there is on average one change-point per 5 time-points), but they behave similarly to 𝒮_Sh_ for the more volatile environments (i.e., for higher values of *p*_*c*_). We note that *p*_*c*_ *>* 0.2 (i.e., when there is on average more than one change-point per 5 time-points)is not quite realistic.

Before the first change-point at trial 51, participants have been in an unpredictable environment for a long time and have no trust in the oracle (and rather a strong belief in the unpredictability of the environment), i.e., *b*^(*t*)^ ≈0 – a situation similar to the extreme left in the plots of Fig. 9. The opposite is true before the second change-point at trial 101, where participants have been in a predictable environment for a long time and have trust in the predictability of the environment, i.e., *b*^(*t*)^ ≈1 – a situation similar to the extreme right in the plots of Fig. 9. We now show that different surprise measures have clearly different predictions after each change-point.

After the first change-point at trial 51, there is (on average) a jump in 𝒮_BF_ (Fig. 10B first row), signaling a change in the environment. After the change, 𝒮_BF_ decreases until the next change at trial 101 where it jumps again, followed by another decreasing period. Thus, 𝒮_BF_ efficiently and explicitly indicates environmental changes. This observation is consistent with the statement of Proposition 1. 𝒮_CC_ (Fig. 10B last row) has a similar behavior; a minor difference compared to S_BF_ occurs after the first change-point where, after a jump, 𝒮_CC_ decreases quickly before it goes up again. This behavior is due to its U-shape relation with *b*^(*t*)^ in Fig. 9. 𝒮_CC_ is the second most informative measure to detect changes.

The behavior of 𝒮_Sh_ and 𝒮_Ba_ (Fig. 10B second and third rows, respectively) are different from 𝒮_BF_ and 𝒮_CC_. One can interpret the temporal average of 𝒮_Sh_ as an indicator of the level of overall unpredictability of the environment – without a distinction between expected and unexpected unpredictability (Soltani & Izquierdo, 2019; Yu & Dayan, 2005). After the first change-point, when the environment becomes more predictable, the average 𝒮_Sh_ gradually decreases from its baseline value, and after the second changepoint, it jumps and then goes back to the original baseline. Finally,_Ba_ has smaller jumps with longer latency after each change-point. This observation is consistent with its inverted-U-shape relation with *b*^(*t*)^ in Fig. 9. Therefore, 𝒮_Sh_ and 𝒮_Ba_ are less informative about environmental changes than 𝒮_BF_ and 𝒮_CC_.

In an actual experiment, a behavioral or biological variable *Z* can be measured throughout the experiment (Fig. 2A1) – similar to our proposed experiments for the example of the oddball tasks (Fig. 7 and Fig. 8). In order to examine with which measure of surprise *Z* correlates or whether *Z* is involved in the biological mechanism behind adaptive learning, one can compute its average over different sequences of stimuli, time-locked to change-points from unpredictable to predictable environments. Because different surprise measures have qualitatively different predictions for this average (c.f. Fig. 10B), we predict that their contributions to *Z* can be dissociated.

To summarize, experiments similar to our proposed experiment can be used for a theory-driven and principled search for signatures of adaptive learning in humans and animals (c.f. Behrens et al., 2007; Marzecová et al., 2019; Nassar et al., 2012; Soltani and Koechlin, 2021; Yu and Dayan, 2005). Moreover, our results indicate that if surprise is used as a signal to modulate learning speed (Gerstner et al., 2018; Iigaya, 2016; Yu & Dayan, 2005), then the Bayes Factor surprise is the best candidate, followed by the Confidence Corrected surprise. We found that neither Shannon surprise nor Bayesian surprise is a reliable indicator of environmental changes. Since the Bayes Factor surprise is the most informative measure about abrupt changes (c.f. Proposition 1), it should be considered in (i) future experimental studies to dissociate the contributions of different surprise measures to explaining behavioral or biological measurements in volatile environments and (ii) future theoretical studies to find normative interpretations for the biological mechanism behind surprise-modulated learning.

### Case-study 3: Exploration by surprise-seeking

In our third case-study, we investigate the link between surprise and decision-making. Seeking surprise has often been interpreted as a strategy for exploring available actions and building an accurate model of the environment (Dubey & Griffiths, 2020; Schulz & Gershman, 2019; Sutton & Barto, 2018). Surprise-seeking exploration strategies have been proven efficient in machine learning (Achiam & Sastry, 2017; Burda et al., 2019; Little & Sommer, 2013; Mobin et al., 2014; Pathak et al., 2017; Schmidhuber, 2010) and have been used to model aspects of human curiosity and exploratory behavior (Dubey & Griffiths, 2020; Gottlieb & Oudeyer, 2018; Gottlieb et al., 2013; Schulz & Gershman, 2019). Here, we study how different surprise measures influence action-selection during surprise-seeking. To do so, we study two thought experiments and propose an actual experiment where seeking surprise gives rise to different exploration strategies, depending on the definition of surprise.

### General setting: Task formulation

All our experiments can be formulated in the form of *reward-free N*-armed bandit tasks (Sutton & Barto, 2018). At time *t*, a participant chooses the arm number *x*_*t*_ ∈ {1, …, *N*} and observes a binary outcome *y*_*t*_ ∈ {1, 2}. In particular, *y*_*t*_ is similar to a reward-indicator in classic bandit tasks (Behrens et al., 2007), but it has no meaning of money or reward in our task. We assume there is no change in the environment so that *p*_*c*_ = 0 and Θ = Θ_1_ = … = Θ_*t*_. The parameter Θ is a vector of *N* real values [*P*_1_, …*P*_*N*_], where *P*_*i*_ (taking value *p*_*i*_ ∈ (0, 1)) is the time-invariant probability of observing *Y*_*t*_ = 1 after taking the arm *X*_*t*_ = *i* – similar to the reward probability in classic bandit tasks except that no reward is given. More precisely, the time-invariant conditional distribution *P*_*Y* |*X*_ is characterized by

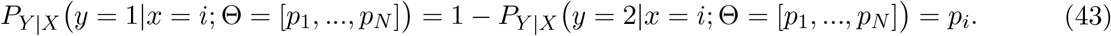

We consider the prior belief to be a Dirichlet distribution^4^

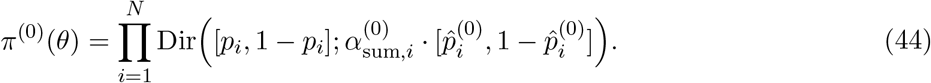

Using exact Bayesian inference, the belief *π*^(*t*)^(*θ*) at time *t* is also a Dirichlet distribution

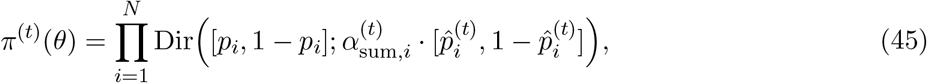

where 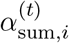 shows how many samples the belief for arm *i* is worth (Efron & Hastie, 2016), and 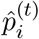 is the latest estimate of the probability of observing *Y*_*t*+1_ = 1 in the next time-step given action *X*_*t*+1_ = *i*. If action *i* has been chosen 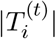 times until time *t*, then we have 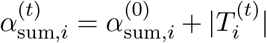 Therefore, 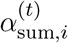 can also be interpreted as a count of how often action *i* has been chosen until time *t*. See Appendix C: Methods for case-studies for details.

### Thought experiment: (i) Optimal model-building

Before turning to surprise-seeking exploration policies, we study the optimal exploration policy in our reward-free bandit task. Imagine that an agent is instructed to always choose its next action *x*_*t*+1_ in a way to find the best estimate 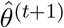 of the environment parameter Θ at the next time-step – i.e., to build the most accurate model of the environment (Schmidhuber, 2010). We define the accuracy of the estimate 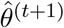 as the squared error between 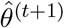 and the true parameter Θ. Then, to obtain (on average) the best estimate of the parameter at the next time-step *t* + 1, the optimal exploration policy is by definition to choose the action *x*_*t*+1_ = *i* that has in expectation the lowest mean squared error

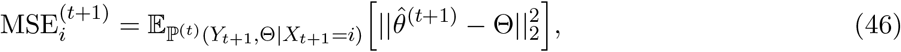

where ||.||_2_ stands for the, *𝓁*_2_-norm. Note that Θ is a random variable, and 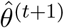 is the agent’s estimate of parameter Θ at *the next time-step t* + 1 which depends on *Y*_*t*+1_ and is itself also a random variable. Thus, the expectation in Eq. 46 is taken over all values of the parameter Θ and *the next observation Y*_*t*+1_, and it is conditioned on the previous actions *x*_1:*t*_, the previous observations *y*_1:*t*_, and *the hypothetical choice i of the next action X*_*t*+1_. The optimal policy can be re-written as choosing the action that maximizes an optimal gain function 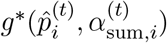, i.e.,

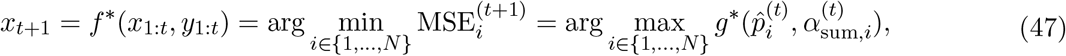

where (c.f. Appendix C: Methods for case-studies and Fig. 11A)

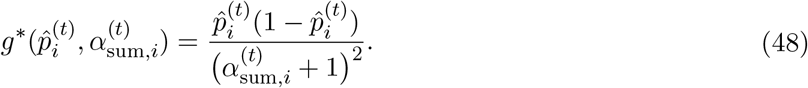

**Figure 11:**
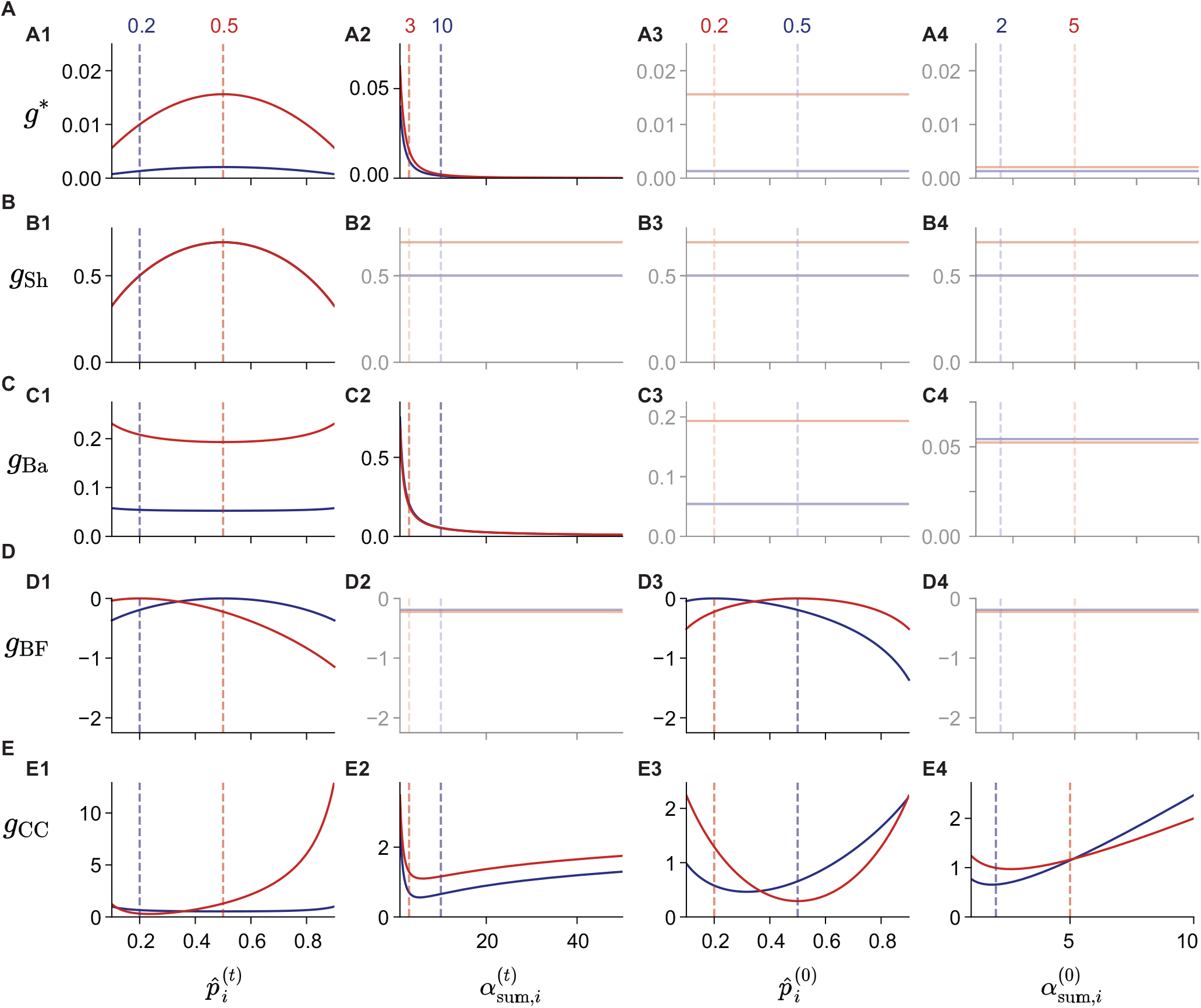
Gain functions for the reward-free bandit task. The optimal exploration policy and different surprise-seeking policies can be written as maximizing a gain function that depends on the current belief parameters 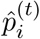 and 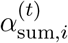 and the prior belief parameters 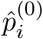 and 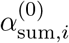. For two different choices of parameters, the blue 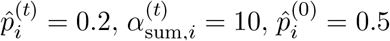, and 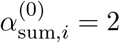) and the red (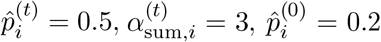 and 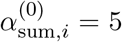) curves show the gain functions for **A**. the optimal policy and the policy based on seeking **B**. the Shannon (B1: the red curve covers the blue one), **C**. the Bayesian, **D**. the Bayes Factor, and **E**. the Confidence Corrected surprise as functions of 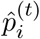 (left column, A1-E1), 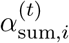 (A2-E2), 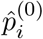 (A3-E3), and 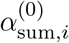 (A4-E4). For the solid red curves in the left column (A1-E1), 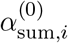,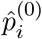 and 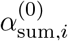 are fixed at their values corresponding to the red dashed lines in the other columns. The same rule applies to the blue curves and to the other columns. Faded panels indicate independence, i.e., when the gain function does not depend on (is constant with respect to) the corresponding parameter. See Appendix C: Methods for case-studies for details.

The optimal gain function *g*^∗^ indicates which action should be chosen (Fig. 11A). To grasp the idea of the optimal policy, let us first imagine that we have chosen arm 1 and 2 ten times each. If we observed *Y*_*t*_ = 1 after every single time we chose arm 1 but after only 50% of times when we chose arm 2, then we would naturally be more confident about our estimate of *p*_1_ than our estimate of *p*_2_. Thus, in order to increase the precision of our estimates, we should keep choosing arm 2. Consistent with this intuition, the optimal gain function *g*^∗^ has an inverted-U-relation with the estimated probability 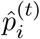 with its maximum at 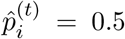 (Fig. 11A1). This means that the optimal policy is to pick the arm with the highest stochasticity in its outcome distribution (among the arms with the same 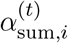).

Second, let us imagine that we have chosen arm 1 only once but have chosen arm 2 ten times. In this case, whatever the actual observations, we would still be less confident about our estimate of *p*_1_ than our estimate of *p*_2_. Thus, in order to increase the precision of our estimates, we should choose arm 1 more often. Consistent with this intuition, the optimal gain function *g*^∗^ is a decreasing function of 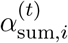 implying that the more often an agent chooses arm *i*, the less informative it becomes (Fig. 11A2). Moreover, for any two arms *i* and *j* with different estimated probabilities 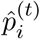 and 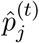, if arm *i* is less stochastic than the arm *j* (i.e., if 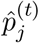 is closer to 0.5), then there are always choice counts 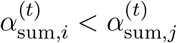 such that the arm *i* with less stochastic outcome gets selected according to *g*^∗^ – for example, in Fig. 11A1, the maximum of the blue curve (for 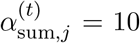) at 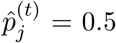 is below the minimums of the red curve 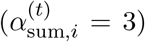 at 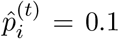 and 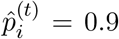. This implies that also the least stochastic arm gets selected occasionally and all arms get selected infinitely many times in the limit *t* → ∞. We note that *g*^∗^ is independent of the prior parameters 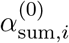 and 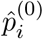 (Fig. 11A3 and A4).

To summarize, the optimal exploration policy prefers arms that (i) are more stochastic and (ii) have been chosen less often. See Appendix C: Methods for case-studies for details and proofs.

### Thought experiment: (ii) Surprise-seeking

Now, imagine that the primary goal of an agent, instead of building the most accurate model of the environment, is to choose its next action *x*_*t*+1_ in a way to be maximally surprised at the next time-step *t* + 1 (Burda et al., 2019; Little & Sommer, 2013; Mobin et al., 2014; Pathak et al., 2017; Storck et al., 1995). Our goal is to study different surprise-seeking exploration policies and compare them with the optimal policy for model-building discussed in the first thought experiment.

Given a measure of surprise 𝒮, a classic surprise-seeking behavior is to choose the action *i* which maximizes the expected 𝒮 of the next observation (Burda et al., 2019; Little & Sommer, 2013; Mobin et al., 2014; Pathak et al., 2017; Storck et al., 1995). Formally, we write

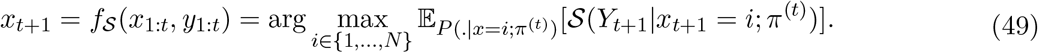

It is important to note that the policy in Eq. 49 is not invariant with respect to non-linear transformations of surprise measures. This is in contrast to our previous case-studies where non-linear transformations of surprise measures did not change their qualitative behavior (c.f. Fig. 2A). Eq. 49 will be applied to all surprise measures except 𝒮_BF_. For the Bayes Factor surprise, we define the surprise-seeking policy as

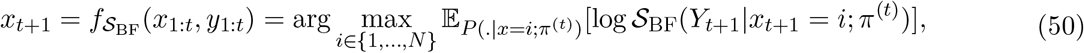

which is the same as seeking the difference in Shannon surprise (c.f. Proposition 2). Note that, because 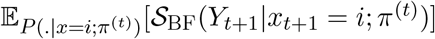 is always and by definition equal to 1, classic surprise-seeking (Eq. 49) with the Bayes Factor surprise is equivalent to uniformly random exploration.

Similar to the case of the optimal exploration policy (Eq. 47), we can define a gain function for each measure of surprise and write the corresponding surprise-seeking policy as the one that maximizes that gain function, i.e., in general, for a measure of surprise 𝒮, the surprise-seeking policy can be written as

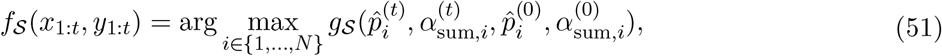

with *g*_𝒮_ the corresponding gain function; see Appendix C: Methods for case-studies for details. Different surprise measures give rise to different gain functions and show different preferences over actions (Fig. 11).

According to *g*_Sh_ (Fig. 11B1), arms with more stochastic outcomes (i.e., with 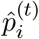 closer to 0.5) are preferred to arms with more deterministic outcomes (i.e., with 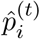 closer to 0 or 1), in agreement with the optimal gain function *g*^∗^ (Fig. 11A1). However, according to *g*_Ba_ the opposite is true (Fig. 11C1). The reason is that improbable events lead to huge changes in the agent’s belief such that more deterministic arms have higher expected 𝒮_Ba_. On the other hand, *g*_Ba_ is a decreasing function of 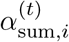 (Fig. 11C2) and in agreement with *g*^∗^ (Fig. 11A2). The preference of *g*_Ba_ for deterministic arms also decreases with increasing 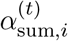 – e.g., the maximum of the blue curve in Fig. 11C1 has a smaller value than the minimum of the red curve. This means that independently of the difference in the stochasticity of their outcomes, *g*_Ba_ will eventually choose arms that have been chosen less often, in agreement with *g*^∗^. However, since 𝒮_Sh_ is a probabilistic mismatch surprise measure (Fig. 3), *g*_Sh_ is independent of 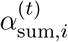: How often an arm has been chosen does not change the preference of *g*_Sh_ for more stochastic arms (Fig. 11B1 and Fig. 11B2). This observation is consistent with different behaviors of 𝒮_Sh_ and 𝒮_Ba_ with respect to confidence in our first case-study (Fig. 5). Therefore, seeking 𝒮_Ba_ has an asymptotic behavior similar to that of the optimal policy, whereas seeking 𝒮_Sh_ remains systematically different from the optimal policy (Fig. 11A-C). We note that both 𝒮_Sh_ and 𝒮_Ba_ are independent of the prior parameters 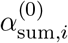 and 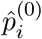 (Fig. 11B3-4 and Fig. 11C3-4, respectively). See Appendix C: Methods for case-studies for details and proofs.

𝒮_BF_ and 𝒮_CC_ rely by definition (Eq. 5 and Eq. 31, respectively) on a comparison between the current belief *π*^(*t*)^ and the prior belief *π*^(0)^. As a result, the prior parameters 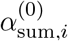 and 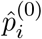 have a potential influence on *g*_BF_ and *g*_CC_ (Fig. 11D3-4 and E3-4, respectively) – in contrast to the optimal policy (Fig. 11A3-4). In particular, *g*_BF_ prefers arms for which the latest estimate of the outcome probability 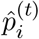 differs *least* from the prior outcome probability 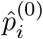 because the the Bayes Factor surprise 𝒮_BF_ tends to be high for samples that are likely to come from the prior distribution. Therefore, *g*_BF_ has an inverted-U-relation with both 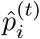 and 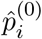 (Fig. 11D1 and D3, respectively). If we have 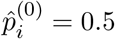 for all arms, then *g*_BF_ has the same preference as that of *g*_Sh_ – since 𝒮_BF_ and 𝒮_Sh_ are indistinguishable in this case (Fig. 2). *g*_BF_ is independent of 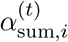 and 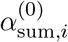 (Fig. 11D2 and Fig. 11D4). See Appendix C: Methods for case-studies for details and proofs.

The gain function for seeking the Confidence Corrected surprise *g*_CC_ prefers arms for which the latest belief about outcome probability *π*^(*t*)^(*p*_*i*_) differs *most* from the prior belief about outcome probability *π*^(0)^(*p*_*i*_). Therefore, the behavior of *g*_CC_ with respect to 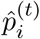 and 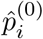 (Fig. 11E1 and E3, respectively) is qualitatively opposite to that of *g*_BF_ (Fig. 11D1 and D3, respectively). Overall, the preference of *g*_CC_ is different from that of the optimal gain function *g*^∗^ with respect to all parameters: 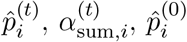, and 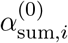 (Fig. 11E1-4). See Appendix C: Methods for case-studies for details and proofs.

To summarize, seeking surprise gives rise to different *sub-optimal* exploration policies depending on the definition of surprise. Seeking the Bayesian surprise gets eventually closer to the optimal policy with increasing 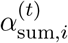, but seeking the Shannon or the Bayes Factor surprise is always systematically different from the optimal policy. The preferences when seeking the Confidence Corrected surprise are often opposite to the preferences of the optimal policy.

### Proposed experiment: Exploration in reward-free 10-armed bandit

In this section, we propose an actual experiment to study human exploratory behavior and test predictions of different exploration policies. We consider a 10-armed bandit task. To motivate participants to explore in the absence of any explicit reward, they are informed that their monetary payoff at the end of the experiment increases the closer their prediction 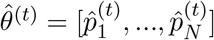 matches the true parameter *θ* = [*p*_1_, …, *p*_*N*_]:

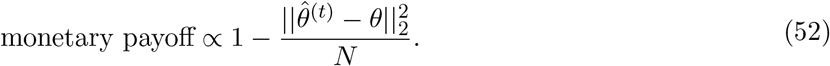

Therefore, to maximize payoff, participants need to choose actions *x*_1:*t*_ so as to best update their estimates 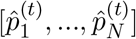 of the parameter. In order to dissociate learning and memory from decision-making, the Bayes-optimal estimate for 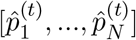 and its corresponding confidence can be updated automatically and shown on a computer screen (e.g., see Kobayashi et al., 2019; Prat-Carrabin et al., 2021). Then the only task of participants is to choose a strategy for action-selection – i.e., to choose the next action *x*_*t*+1_.

Our goal is to study (i) how well each policy performs in terms of model-building, and (ii) whether we can dissociate between different policies if we can only observe a finite number of their action choices. To do so, we first (i) simulate different policies and measure their performance over time. Then, we (ii) apply model-recovery (Wilson & Collins, 2019) to the simulated data in order to see if we can recover the true policy from a finite number of action-choices – i.e., to see if our task can distinguish between different policies on the level of action-choices. See Appendix C: Methods for case-studies for details. We consider two scenarios: in the first scenario, the prior outcome probability 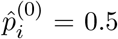 is the same for all arms – with 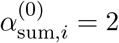 in the second scenario, 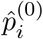 is chosen from a wide interval from 0.05 to 0.95 – with 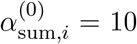. In the first scenario, the effect of 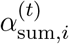 on the policies is dominant, whereas in the second scenario, 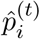 also plays a role in action-selection.

In the 1st scenario (Fig. 12A), maximizing *g*_Ba_ is almost as good as maximizing the optimal gain function *g*^∗^, and maximizing *g*_Sh_ is the same as maximizing *g*_BF_ – as expected from the shape of the gain functions (Fig. 11). Maximizing *g*_CC_ is the worst policy. Our results for model-recovery (Fig. 12B) show that different exploration policies can be distinguished from each other given observed action-choices – except for *g*_Sh_ and *g*_BF_ which are essentially the same in the 1st scenario. Importantly, despite the similar performance of *g*^∗^ and *g*_Ba_ in model-building (Fig. 12A), they are distinguishable at the level of action-choices (Fig. 12B).

**Figure 12:**
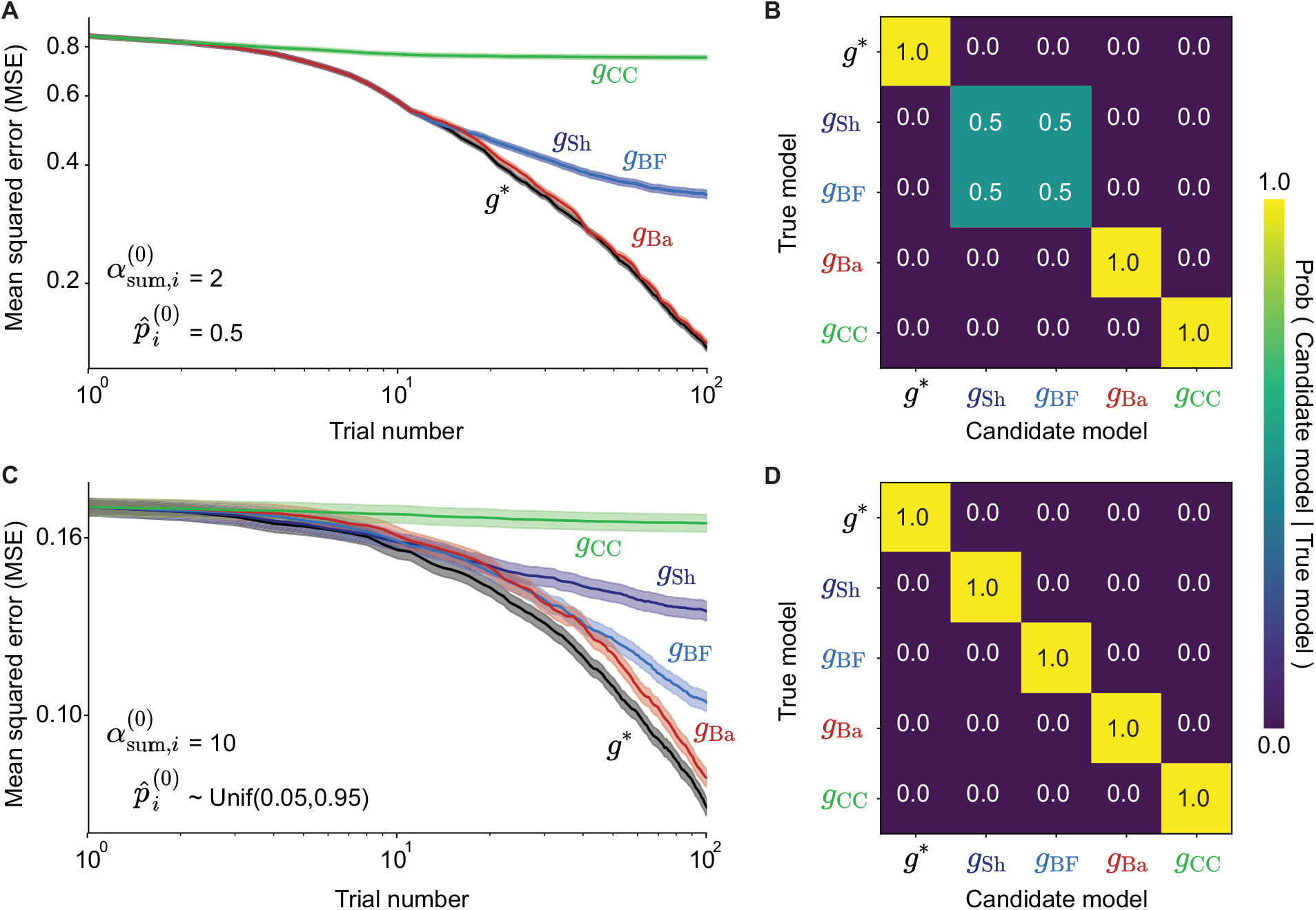
Exploration performance and model-recovery in a reward-free 10-armed bandit task. **A**. The mean squared error (MSE) of the estimate of the optimal policy (*g*^∗^) and different surprise-seeking policies as a function of time *t* – averaged over 500 random seeds. The prior parameters 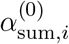 and 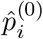 were fixed for all arms at 2 and 0.5, respectively. The shaded areas indicate the standard error of the mean. **B**. Confusion matrix for model recovery (Wilson & Collins, 2019) using data shown in panel A. We used classic Bayesian model selection (Efron & Hastie, 2016; Kass & Raftery, 1995) and, given a sequence of action-choices and observations (for one random seed), computed the posterior probability of different models. Each cell shows the posterior probability of a candidate model (corresponding column) given that the action-choices were made by one of the true models (corresponding row) – averaged over 500 random seeds. **C**. and **D**. The same as panels A and B, respectively, except that the prior parameter 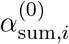 wasfixed at 10 for all arms, but 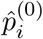 for each arm was chosen randomly between 0.05 and 0.95. Seeking Bayesian surprise (*g*_Ba_, red) remains closest to the optimal policy also for other choices and distributionsof the prior parameters 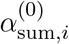 and 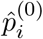 See Appendix C: Methods for case-studies for details.

In the second scenario, maximizing *g*_Sh_ differs from maximizing *g*_BF_ (Fig. 12C). By constantly comparing 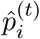 with 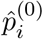, seeking 𝒮_BF_ has an indirect preference for arms that have been chosen less often; therefore, seeking S_BF_ has a better performance than maximizing *g*_Sh_. While the best surprise-seeking policy is still to maximize *g*_Ba_, its difference with the optimal policy becomes more obvious in the second scenario – when 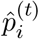 matters. In this scenario, *g*_CC_ is again the worst policy and all policies can be distinguished from each other given their action-choices (Fig. 12D).

To summarize, our proposed experiment can distinguish different exploration policies both at the level of (i) performance and (ii) action-choices with as few as 100 trials. We found that seeking the Bayesian surprise leads to an exploration policy closest to the optimal policy for model-building. This explains the popularity of the Bayesian surprise in the field of machine learning (Little & Sommer, 2013; Mobin et al., 2014; Schmidhuber, 2010; Storck et al., 1995). Seeking the Shannon (or Bayes Factor) surprise differs more from the optimal policy as time passes – as a result of their indifference with respect to 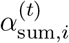. Seeking the Confidence Corrected surprise leads to a policy that favors situations that differ most from prior expectations, and the resulting policy is often opposite to the optimal policy. As a result, a reward-free multi-armed bandit task as described above can be used to study the degree of suboptimality of human exploration strategies and be used to compare them with different models of human curiosity (Dubey & Griffiths, 2020; Gottlieb & Oudeyer, 2018) and surprise-seeking policies.

### Summary and intermediate discussion (ii)

In the three case-studies, we showed that the 4 different categories of surprise in Fig. 4 have different predictions regarding (i) the perception of surprise, (ii) adaptive learning by surprise, and (iii) exploration by surprise-seeking. We designed experiments where these predictions can be tested in practice. Our theoretical analyses, therefore, can be used to correlate physiological variables with different measures of surprise, search for signatures of adaptive learning in the brain, and study human curiosity and exploratory behavior.

Our results suggest that the Bayes Factor surprise is the most suitable measure of surprise to modulate learning in volatile environments (Frémaux & Gerstner, 2016; Gerstner et al., 2018; Iigaya, 2016; Liakoni et al., 2021; Soltani & Izquierdo, 2019; Yu & Dayan, 2005) and the Bayesian surprise is the most suitable measure of surprise to model curiosity as an innate mechanism in humans for exploration – whenever model-building is crucial for better future performance (Dubey & Griffiths, 2019, 2020; Schmidhuber, 2010; Singh, Lewis, & Barto, 2010; Singh, Lewis, Barto, & Sorg, 2010; Storck et al., 1995). In particular, we showed that a surprise measure that is useful to modulate the learning speed should necessarily compare the current belief with the prior belief (Fig. 9 and Fig. 10), while we also showed that such a comparison leads to suboptimal exploration in surprise-seeking policies (Fig. 11 and Fig. 12). Similarly, we showed that a surprise measure that is useful for exploration should increase with increasing uncertainty (Fig. 11 and Fig. 12), whereas we also showed that such a behavior is slow and suboptimal in detecting change-points (Fig. 9 and Fig. 10). In other words, our results show that the very features that make the Bayes Factor surprise an appropriate measure for learning make it an unsuitable measure for exploration and vice versa for the Bayesian surprise.

Unlike the Bayes Factor and the Bayesian surprise, the Shannon surprise (c.f. Eq. 9 and Eq. 10) considers less likely events always as more surprising. Surprise in natural language is defined as ‘the feeling or emotion excited by something unexpected’ (Oxford English Dictionary, n.d.). If we focus on the term ‘unexpected’, identify it with ‘unlikely under the current belief’, and neglect the terms ‘feeling’ and ‘emotion’, then our results suggest that the Shannon surprise measures a quality closely related to the definition of surprise in natural language (i.e., the dictionary definition of surprise). However, we observed that the Confidence Corrected surprise has a more intuitive behavior in some cases where confidence (or commitment to a belief) plays an important role (Fig. 6), despite its counter-intuitive behavior in some other situations (Fig. 5).

### Regularized Shannon surprise: A new direction

We argued that the Confidence Corrected surprise has a more intuitive behavior than the other measures in the CEO selection experiment because it explicitly accounts for confidence (Fig. 6). However, because it treats the confidence for a *correct* prediction in the same way as the confidence for a *wrong* prediction, its behavior is against common sense in some other experiments (Fig. 5 and Fig. 7). In this section, we overcome this issue by providing a modified definition for the Shannon surprise which only penalizes the confidence for a *wrong* prediction.

We call the modified measure the *Regularized Shannon surprise* and define it as

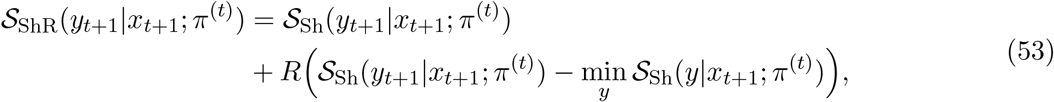

where *R* : ℝ^+^ →ℝ^+^ can be any continuous function that satisfies: (i) *R*(0) = 0, and (ii) *R*(*z*) is an increasing function of *z* for all *z* ∈ℝ^+^. As a result, we have the following two properties for the regularized Shannon surprise:

1. Whenever *y*_*t*+1_ is the most expected observation, i.e., *y*_*t*+1_ = arg min_*y*_ 𝒮_Sh_(*y*|*x*_*t*+1_; *π*^(*t*)^), we have 𝒮_ShR_(*y*_*t*+1_|*x*_*t*+1_; *π*^(*t*)^) = 𝒮_Sh_(*y*_*t*+1_|*x*_*t*+1_; *π*^(*t*)^). This means that the confidence for a correct prediction does not penalize surprise.
2. Whenever *y*_*t*+1_ is not the most expected observation, the difference between the regularized Shannon surprise and the Shannon surprise, i.e., 𝒮_ShR_(*y*_*t*+1_|*x*_*t*+1_; *π*^(*t*)^) −𝒮_Sh_(*y*_*t*+1_|*x*_*t*+1_; *π*^(*t*)^), is a decreasing function of the Shannon surprise of the most likely observation 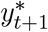 = arg min_*y*_ 𝒮_Sh_(*y*|*x*_*t*+1_; *π*^(*t*)^). This means that for a fixed Shannon surprise of *y*_*t*+1_, the more we expect another observation 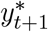, the more surprised we are by observing 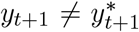, consistent with our expectation for the CEO selection experiment (Fig. 6A1).

As a simple choice, let us consider *R*(*z*) = *z*, i.e.,

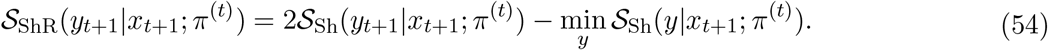

Then, the predictions of the regularized Shannon surprise 𝒮_ShR_ are the same as the predictions of the Shannon surprise 𝒮_Sh_ for the cases where 𝒮_Sh_ in the coin flipping thought experiment and the classic oddball task (compare Fig. 13A-E with Fig. 5 and Fig. 7A-C). Importantly, the EEG amplitude at around 450ms in the visual oddball task we analyzed has a behavior consistent with the predictions of 𝒮_ShR_ but not with the predictions of 𝒮_CC_ (compare Fig. 13D-E with Fig. 7). In the belief in forecast and association learning experiments, 𝒮_ShR_, similar to 𝒮_Sh_, can be interpreted as a measure of unpredictability and unexpectedness in the environment (Fig. 13I-K). In this regard, S_ShR_ is also a good model of the dictionary definition of surprise. On the other hand, in the CEO selection experiment and the generalized oddball task, the confidence regularization in 𝒮_ShR_ makes it more similar to 𝒮_CC_ than 𝒮_Sh_ (compare Fig. 13F-H with Fig. 6 and Fig. 8): Given a fixed estimated probability 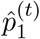 for the selection of the 1st candidate (*Y*_*t*+1_ = 1), the regularized Shannon surprise for *Y*_*t*+1_ = 1 increases with increasing expectation for the selection of another candidate (e.g., *Y*_*t*+1_ = 2) – in agreement with the results of SanMiguel et al., 2021. Therefore, 𝒮_ShR_ captures those features of 𝒮_Sh_ that make it similar to the dictionary definition of surprise but adds a dependence upon confidence.

**Figure 13:**
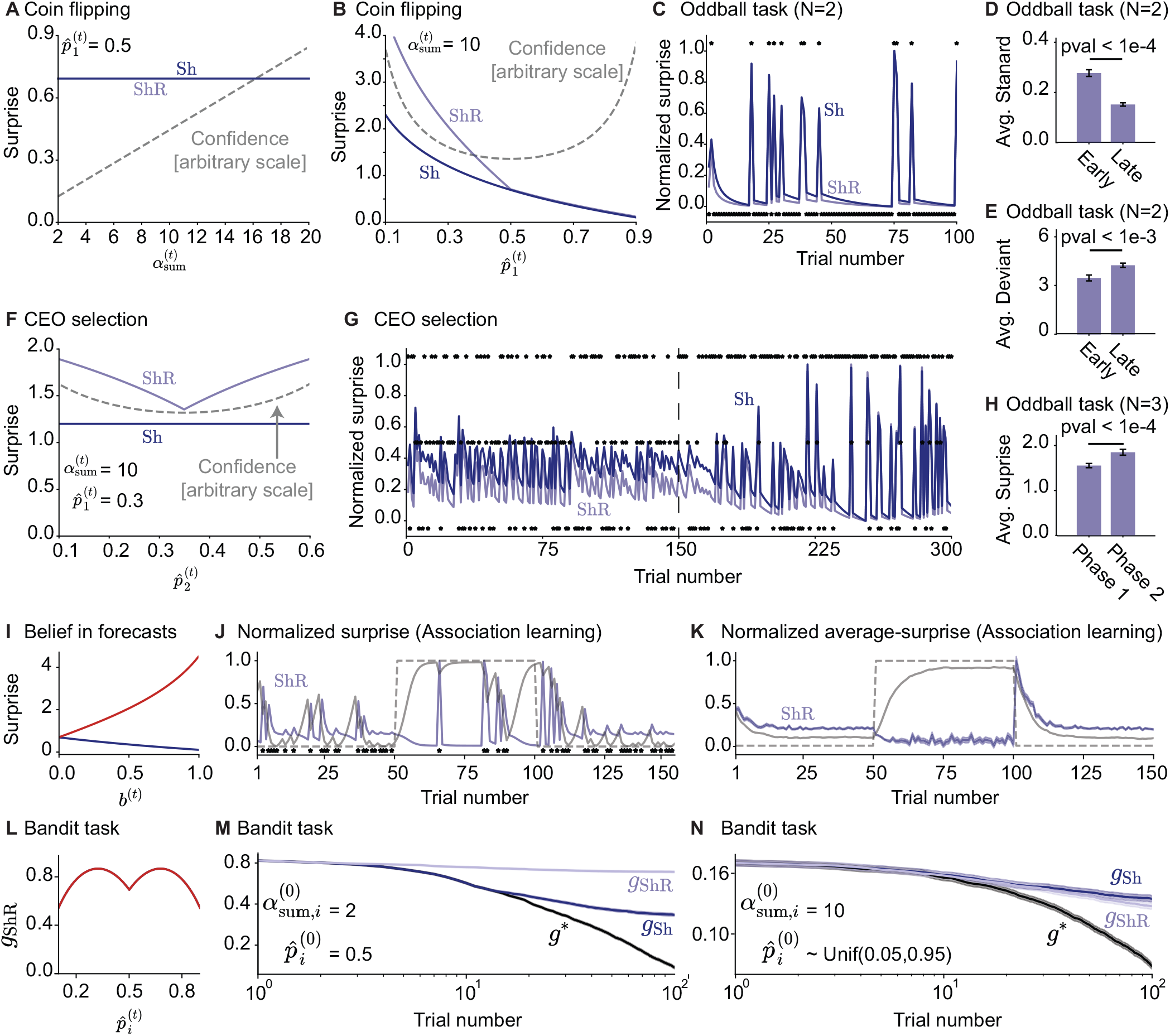
The behavior of the regularized Shannon surprise for different case-studies. Each panel corresponds to one panel in Fig. 5-Fig. 12. Sh (dark blue) corresponds to the Shannon surprise and ShR (light blue) corresponds to the regularized Shannon surprise. The regularized Shannon surprise has the same behavior as the Shannon surprise in panels A-E as well as in panels I-K, but it has the same behavior as the Confidence Corrected surprise in panels F-H. **A**. Coin flipping experiment: Surprise of *Yt*+1= 1 as a function of 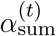 (c.f. Fig. 5A1). **B**. Coin flipping experiment: Surprise of *Y*_*t*+1_ = 1 as a function of 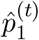 (c.f. Fig. 5B1). **C**. Classic oddball task: Surprise over time for standard (shown at -0.05) and deviant (shown at 1.05) stimuli (c.f. Fig. 7A). **D**. Classic oddball task: Average surprise of standard stimuli in the early and the late phase of the task (c.f. Fig. 7B). **E**. Classic oddball task: Average surprise of deviant stimuli in the early and the late phase of the task (c.f. Fig. 7C).**F**. CEO selection experiment: Surprise of *Y*_*t*+1_ = 1 as a function of 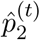 (c.f. Fig. 6A1). **G**. CEO selection experiment: Surprise over time (c.f. Fig. 8A). **H**. Generalized oddball task: Average surprise values of *Y*_*t*+1_ = 1 in Phase 1 and Phase 2 (c.f. Fig. 8C). **I**. Belief in forecasts experiment: The regularized Shannon surprise as a function of trust in media *b*^(*t*)^ for media being right (blue) or wrong (red) ; *b*^(0)^ = 0.5 (c.f. Fig. 9B). **J**. Association learning experiment: Surprise over time for one random seed; stars indicate when oracle’s prediction has been wrong (c.f. Fig. 10A). **K**. Association learning experiment: Surprise over time (as in panel J) averaged over 500 random seeds (c.f. Fig. 10B). **L**. Bandit experiment: The gain function *g*_ShR_ corresponding to the policy of seeking ShR as a function of 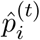 (c.f. Fig. 11). *g*_ShR_ is independent of 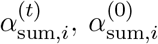, and 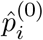 **M-N**. Bandit experiment: Exploration performance for seeking ShR (light blue) compared to seeking Sh (dark blue) and the optimal policy *g*^∗^ (black) for **M**. 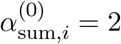 and 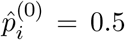 (c.f. Fig. 12A) and **N**. 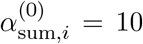 and 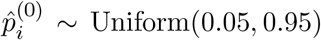 (c.f. Fig. 12B). See Appendix E: Regularized Shannon surprise for details.

Finally, our results show that, similar to seeking 𝒮_Sh_ or 𝒮_CC_, seeking 𝒮_ShR_ is a sub-optimal exploration policy and has a poor performance for model-building (Fig. 13L-N and Appendix C: Methods for case-studies).

## General discussion

What does it formally mean to be surprised? How can we experimentally distinguish between different surprise measures? How do different surprise measures relate to different computational roles of surprise in the brain? And how do they relate to the word ‘surprise’ in natural language? To address these question, we reviewed 16 surprise measures in a common mathematical framework and studied their links to perception, learning, and decision-making. We identified the conditions under which they are indistinguishable (Fig. 2) and provided a technical (Fig. 3) and a conceptual (Fig. 4) categorization of these measures.

Our results suggest that the class of prediction surprise measures (Fig. 4) is the closest one to the dictionary definition of surprise (see Summary and intermediate discussion (ii)). However, following Faraji et al., 2018, we also suggested that a surprise measure should explicitly capture the notion of confidence in order to explain intuitive aspects of surprise perception. We therefore proposed the ‘Regularized Shannon surprise’ (Eq. 53) and show how its predictions can be tested experimentally. Our predictions are supported by recent experimental evidence (SanMiguel et al., 2021).

We found that the very features that make a surprise measure suitable for adaptive learning are in conflict with the ones that make it suitable for exploration. In particular, adaptive behavior as observed in humans is achieved only if a surprise from the class of change-point detection measures modulates learning (Fig. 4), whereas a close-to-optimal exploration strategy is achieved only if a surprise from the class of information-gain surprise measures drives action-selection (Fig. 4). Change-point detection surprise measures compare the probability of an event under the prior belief with that under the current belief, which is why they are useful in adaptive learning (Liakoni et al., 2021; Soltani & Izquierdo, 2019; Yu & Dayan, 2005). In the sense of ‘unexpected under the current belief’, change-point detection surprise measures are also similar to the dictionary definition of surprise. Information-gain surprise measures, however, essentially differ from the definition of surprise in natural language. We, therefore, suggest to avoid the term ‘surprise’ when referring to these measures. Specifically, we suggest the names ‘Bayesian information gain’ and ‘Postdictive belief-update’ instead of ‘Bayesian surprise’ and ‘Postdictive surprise’ – similar to Kolossa et al., 2015.

Based on our theoretical results, we proposed experimental paradigms where different surprise categories make different predictions. A natural direction for future studies is to test these predictions experimentally. A wide range of questions can be addressed, either directly or indirectly, such as ‘which surprise-seeking policy is most similar to human exploratory behavior in different experiments?’ (Dubey & Griffiths, 2020; Gottlieb & Oudeyer, 2018), ‘which surprise measure correlates best with pupil diameter (Antony et al., 2021; Nassar et al., 2012; Preuschoff et al., 2011), with different EEG components (Gijsen et al., 2021; Kolossa et al., 2015; Mousavi et al., 2020; Visalli et al., 2021), or with fMRI bold activity (Gläscher et al., 2010; Konovalov & Krajbich, 2018; Loued-Khenissi & Preuschoff, 2020)?’, and even ‘which surprise measure explains best the neural activity in the locus coeruleus?’ (Sara, 2009).

Some important open questions on the mechanistic level (Marr, 1982) are ‘which neural circuits are involved in computation of different surprise measures?’ (see Fiser et al., 2010; Knill and Pouget, 2004; Soltani and Wang, 2010 for examples of neural models of probabilistic inference), ‘how can surprise-modulated adaptive learning be implemented at the level of synaptic plasticity?’ (Berlemont & Nadal, 2021; Gerstner et al., 2018; Iigaya, 2016; Illing et al., 2021), and ‘how can surprise-seeking exploration strategies be implemented in the brain?’ (see Basanisi et al., 2020 for an example). There are also important open questions on the computational and algorithmic levels (Marr, 1982) such as ‘how can we model the influence of attention on surprise perception?’ (Gottlieb & Oudeyer, 2018), ‘how can we formulate other surprise theories like Palm’s theory of surprise (Palm, 2012) or the theory of surprise as a compression measure (Schmidhuber, 2010) in our mathematical framework?’, and ‘how do different surprise measures contribute to formation (Rouhani & Niv, 2021), segmentation (Antony et al., 2021; Rouhani et al., 2020), and modification (Gershman et al., 2017; Sinclair & Barense, 2018) of memory?’.

In conclusion, our results articulate and unify many of the existing paradigms for the study of surprise and suggests various new directions for further theoretical and experimental studies of surprise and its roles in the brain function.

## Appendix A: Special cases and links to related works

Several existing models and experimental paradigms are special cases of Eq. 1 and Eq. 2. The standard generative models for studying passive learning in volatile environments (Adams & MacKay, 2007; Fearnhead & Liu, 2007; Liakoni et al., 2021; Nassar et al., 2012; Nassar et al., 2010; Wilson et al., 2013) is obtained if we remove the cue variables *X*_1:*t*_ (Fig. 1B). For example, in the Gaussian experiment of Nassar et al., 2012; Nassar et al., 2010, *Y*_*t*_ is a sample from a Gaussian distribution with a mean equal to Θ_*t*_ and a known variance, and *π*^(0)^ is a very broad uniform distribution. Liakoni et al., 2021 extensively studied this generative model and showed that both exact and approximate Bayesian inference in this generative model lead to surprise-modulated learning rules. In the next sections, we show that the same holds true for our more general generative model.

Variants of bandit and reversal bandit tasks (Behrens et al., 2007; Findling et al., 2021; Horvath et al., 2021) can be modeled by considering the cue variables *X*_1:*t*_ as actions *A*_1:*t*_ (Fig. 1C). For example, in the experiment of Behrens et al., 2007, *X*_*t*_ = *A*_*t*_ is one of the two possible actions that participants can choose, *Y*_*t*_ is the indicator of whether they are rewarded or not, and Θ_*t*_ indicates which action is rewarded with higher probability. In this setting, ℙ(*x*_*τ*_ |*x*_*τ*−1_, *y*_*τ*−1_) = ℙ (*x*_*τ*_) is the probability that participants take action *x*_*τ*_, independently of the dynamics of the environment.

The minimal model of human inferences about binary sequences of Meyniel et al., 2016 (Fig. 1D) is obtained if the cue variable *X*_*t*_ is equal to the previous observation *Y*_*t*−1_. There, *Y*_*t*_, conditioned on *Y*_*t*−1_, is a sample from a Bernoulli distribution with parameter Θ_*t*_. In this setting, we have 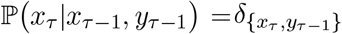. This class of generative models have also been successfully used to explain the neural signatures of surprise via encoding (Gijsen et al., 2021; Maheu et al., 2019; Meyniel, 2020) and decoding (Modirshanechi et al., 2019; Mousavi et al., 2020) models.

Classic Markov Decision Processes (MDPs) (Sutton & Barto, 2018) can also be written in the form of our generative model. To reduce our generative model to an MDP, we set *p*_*c*_ = 0, consider the observation *Y*_*t*_ as the pair of the current state and immediate reward value, and consider the cue variable *X*_*t*_ as the previous pair of action and observation (or state) (*A*_*t*−1_, *Y*_*t*−1_) (Fig. 1E). In this setting, we have 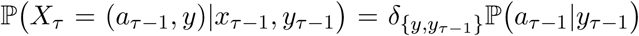, where ℙ(*a*_*τ*−1_|*y*_*τ*−1_) is called the action selection policy in Reinforcement Learning theory (Sutton & Barto, 2018) and is independent of the dynamics of the environment. The theory of Reinforcement Learning for MDPs has been frequently used in neuroscience and psychology to model human reward-driven decision-making (Daw et al., 2011; Gläscher et al., 2010; Huys et al., 2015; Lehmann et al., 2019; Niv et al., 2015; Xu et al., 2021).

## Appendix B: Proofs

In this appendix, we provide proofs for our Propositions and Corollaries mentioned in section Theories of surprise: A technical review in the main text. We also provide further results for the Bayesian and postdictive surprise in Lemma 1 and 2 and Remark 1.

### Proof of Proposition 1

The proof is in essence the same as the proof of Proposition 1 of Liakoni et al., 2021. We write

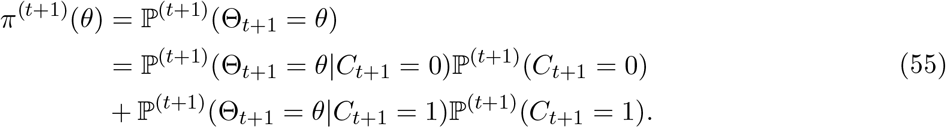

We use Bayes’ rule and write ℙ ^(*t*+1)^(Θ_*t*+1_ = *θ*|*C*_*t*+1_ = 0) (c.f. the 1st term in Eq. 55) as

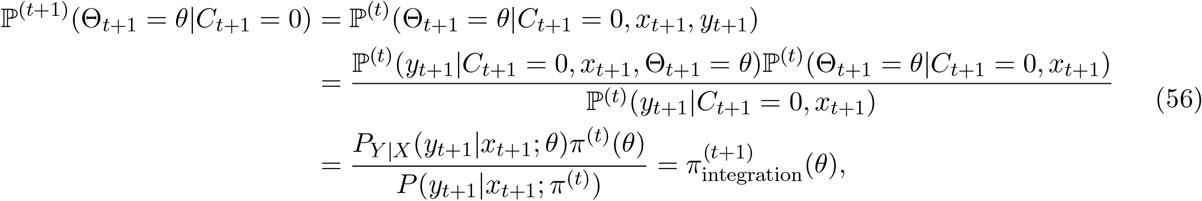

and similarly

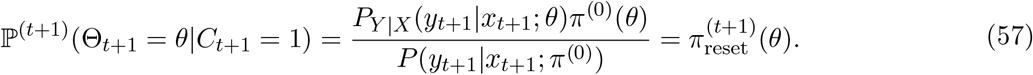

Then, for ℙ ^(*t*+1)^(*C*_*t*+1_ = 1) and ℙ ^(*t*+1)^(*C*_*t*+1_ = 0) = 1 −ℙ ^(*t*+1)^(*C*_*t*+1_ = 1) we have

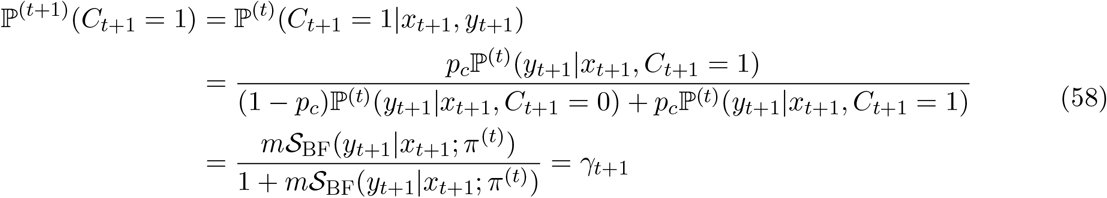

with 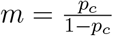. Therefore, the proof is complete by substituting these terms in Eq. 55.■

### Proof of Proposition 2

Based on the definition of the adaptation rate *γ*_*t*+1_ (c.f. Proposition 1), we have

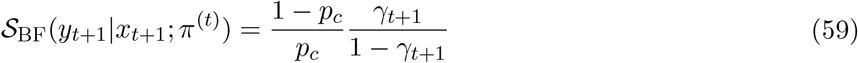

For the difference in the 1st definition of the Shannon surprise (c.f. Eq. 9), we can write

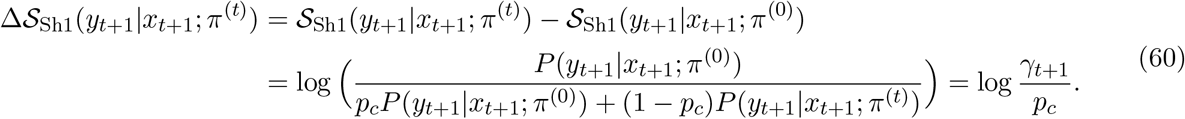

As a result, we have *γ*_*t*+1_ = *p*_*c*_ exp Δ 𝒮 _Sh1_(*y*_*t*+1_|*x*_*t*+1_; *π*^(*t*)^) and hence

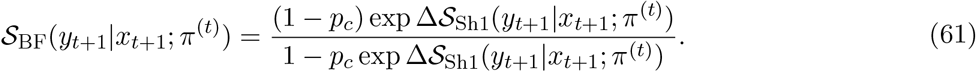

The proof is more straightforward for the difference in the 2nd definition (c.f. Eq. 10) where we have

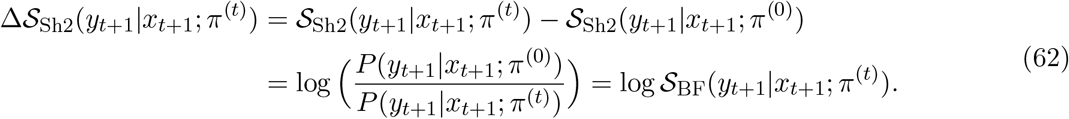

Therefore, the proof is complete. ■

### Proof of Proposition 3

Based on the definitions of the two versions of the Shannon surprise (c.f. Eq. 9 and Eq. 10), we have

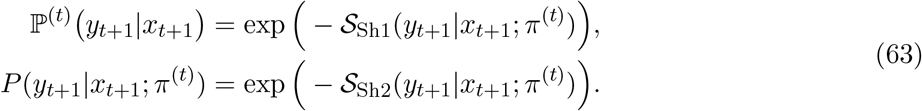

The proof is complete by using these equations and replacing the probabilities in Eq. 14 and Eq. 15. ■

### Proof of Proposition 4

For a categorical task with *N* categories and one-hot coded observations, we have (c.f. Eq. 17 and Eq. 18)

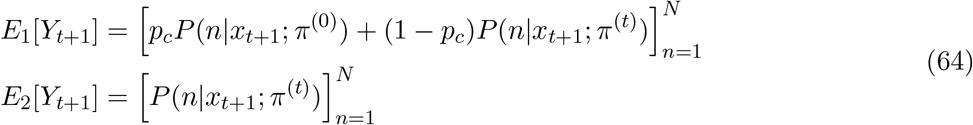

where 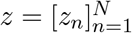 is an *N* -dimensional vector with *z*_*n*_ the *n*th element. To be able to prove the proposition for *E*_1_[*Y*_*t*+1_] and *E*_2_[*Y*_*t*+1_] simultaneously, we define 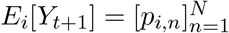, where *p*_1,*n*_ = *p*_*c*_*P* (*n*|*x*_*t*+1_; *π*^(0)^) +(1 − *p*_*c*_)*P* (*n*|*x*_*t*+1_; *π*^(*t*)^) and *p*_2,*n*_ = *P* (*n*|*x*_*t*+1_; *π*^(*t*)^).

We show the one-hot coded vector corresponding to category *m* ∈ {1, …, *N*} by *e*_*m*_. For the absolute error surprise, we have (c.f. Eq. 19)

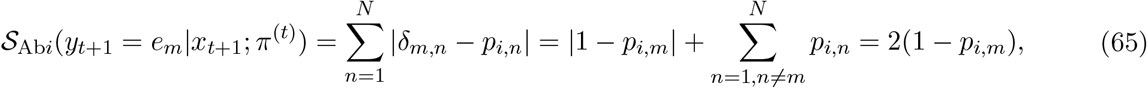

which is the same as 2 𝒮_SPE*i*_(*y*_*t*+1_ = *e*_*m*_|*x*_*t*+1_; *π*^(*t*)^) (c.f. Eq. 14 and Eq. 15).

For the squared error surprise, we have (c.f. Eq. 19)

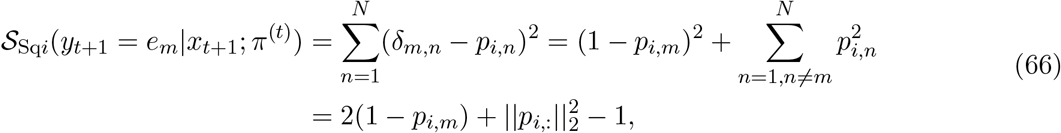

where we have 2(1 − *p*_*i,m*_) = 2 𝒮_SPE*i*_(*y*_*t*+1_ = *e*_*m*_|*x*_*t*+1_; *π*^(*t*)^) and

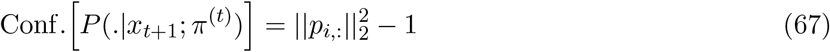

shows the ℓ_2_-norm of the estimate vector *p*_*i*,:_ as a measure of confidence; 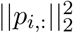 takes its maximum value when the prediction has a probability of 1 for one category and zero for the rest, and 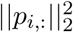 takes its minimum when it is distributed uniformly over all categories. Therefore, the proof is complete. ■

### Proof of Proposition 5

Assume that *Y*_*t*+1_ ∈ ℝ^*N*^ given the cue *x*_*t*+1_ and the belief *π*^(*t*)^ has a Gaussian distribution with a covariance matrix *σ*^2^*I*, i.e.,

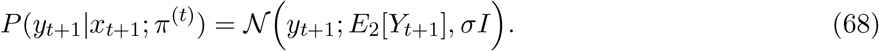

We then have

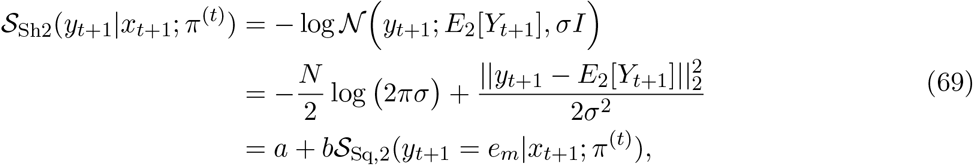

where *a* = −*N* log (2*πσ*) /2 and *b* = 1/(2*σ*^2^). Therefore, the proof is complete. ■

### Theoretical results for the Bayesian surprise

#### Lemma 1.

*(Relation between the Bayesian surprise and the Shannon surprise) In the generative model of Definition 1, the Bayesian surprise can be written as*

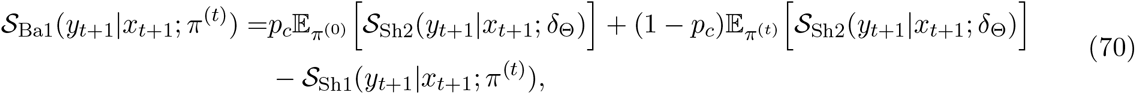

*and*

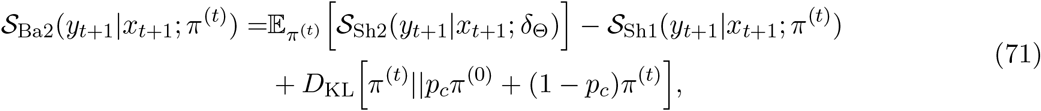

*where δ*_*θ*_ *is a delta-distribution at θ*.

*Proof:* For the 1st definition of the Bayesian surprise (c.f. Eq. 23), we have

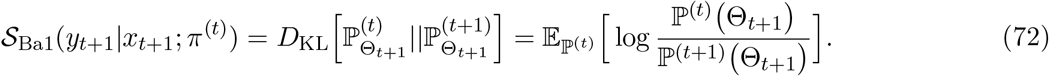

We know

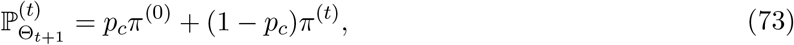

and

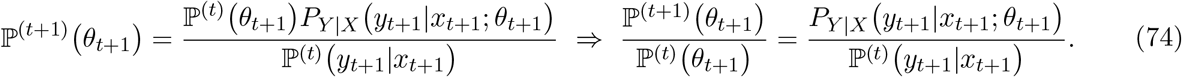

We, therefore, have

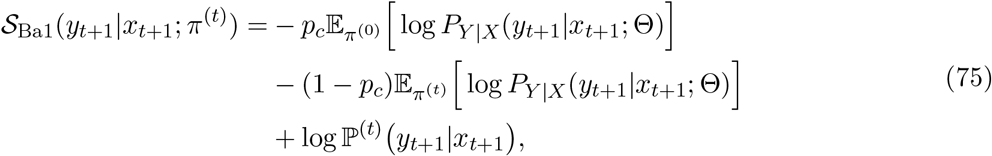

which is equivalent to (c.f. Eq. 9 and Eq. 10)

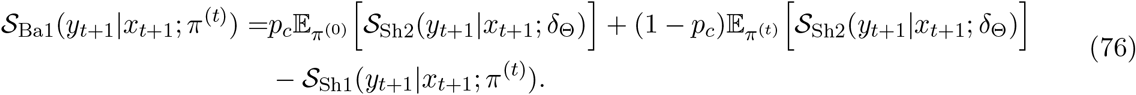

For the 2nd definition of the Bayesian surprise (c.f. Eq. 24), we have

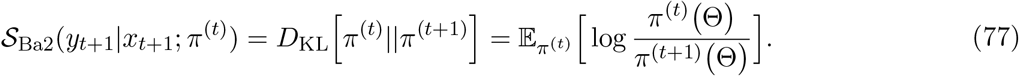

We use Eq. 22 and Eq. 74 and write

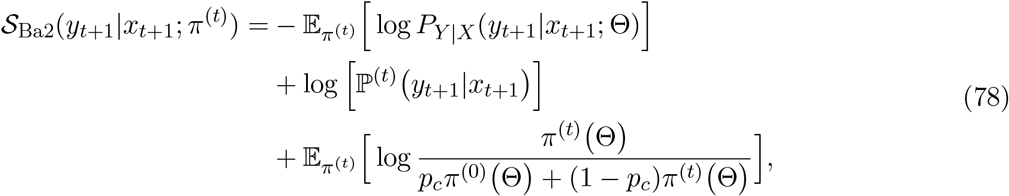

which is equivalent to (c.f. Eq. 9 and Eq. 10)

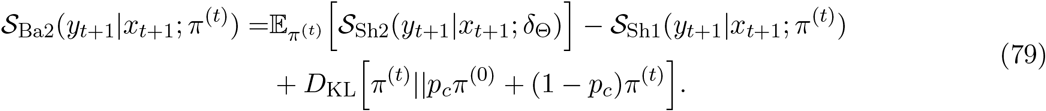

Therefore, the proof is complete. ■

#### Remark 1.

*When the change point probability is zero, i*.*e. p*_*c*_ = 0, *the Bayesian surprise is equal to the expected Shannon surprise minus the Shannon surprise, i*.*e*.,

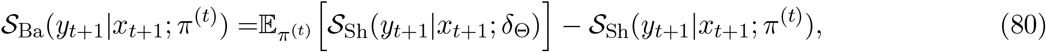

*where* 𝒮_Ba_ = 𝒮_Ba1_ = 𝒮_Ba2_ *and* 𝒮_Sh_ = 𝒮_Sh1_ = 𝒮_Sh2_.

*Proof:* The remark is the direct consequence of Lemma 1. ■

### Theoretical results for the postdictive surprise

#### Lemma 2.

*(Relation between the postdictive surprise and the Shannon surprise) In the generative model of Definition 1, the postdictive surprise can be written as*

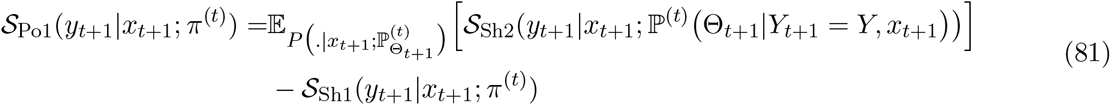

*and*

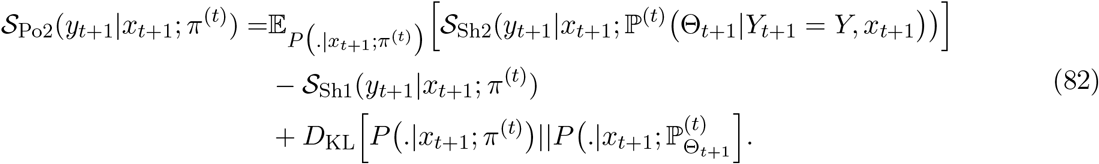

*Proof:* We first prove the equality for 𝒮_Po1_ for which we have (c.f. Eq. 25)

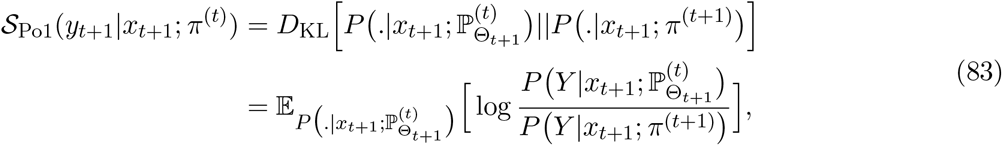

where

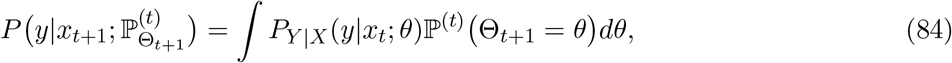

and, using Bayes’ rule,

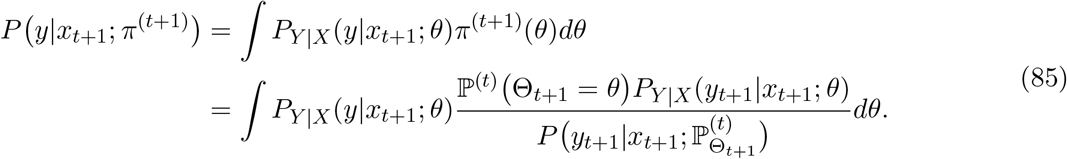

Using the Bayes’ rule and the definition of the marginal probability (c.f. Eq. 4), we can find

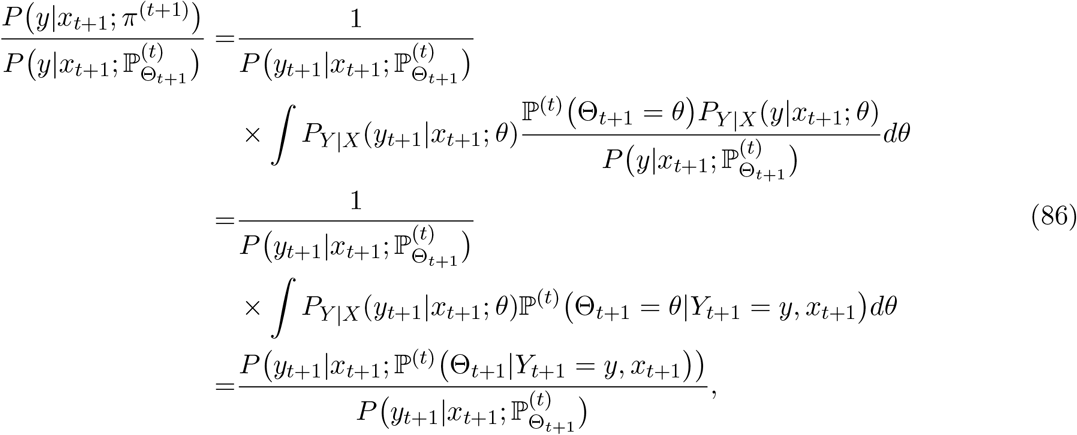

and as a result (using Eq. 9 and Eq. 10)

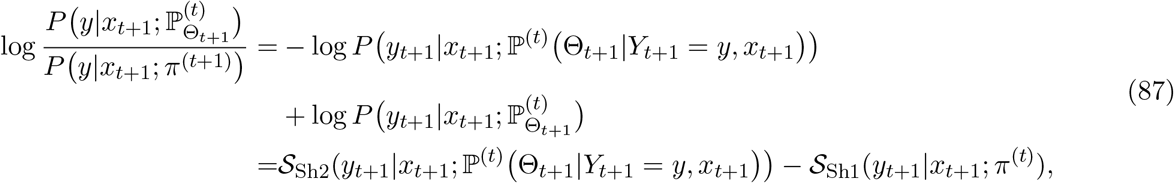

which, using Eq. 83, makes the proof complete.

To prove the 2nd equality, we note that (c.f. Eq. 26)

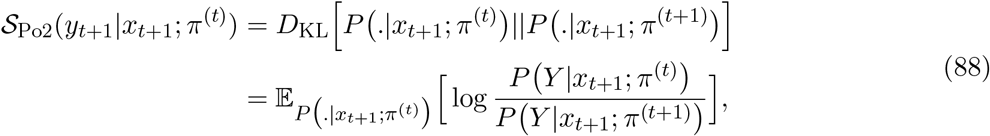

and

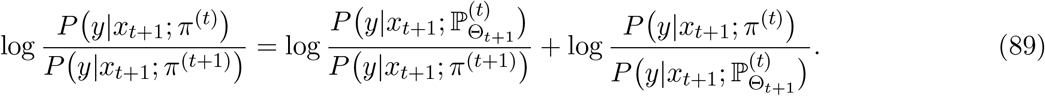

Therefore, using Eq. 87 and the definition of *D*_KL_, the proof is complete. ■

#### Proof of Proposition 6

First, we prove the statement for the 2nd definition of the Confidence Corrected surprise (c.f. Eq. 31) for which we have

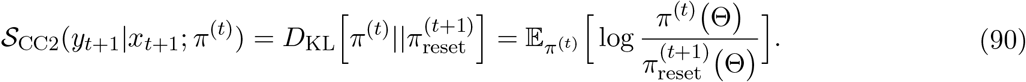

Using the definition of 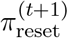in Proposition 1, we can write

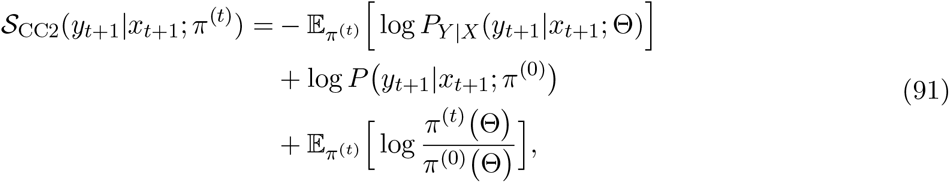

which is equivalent to (c.f. Eq. 9 and Eq. 10)

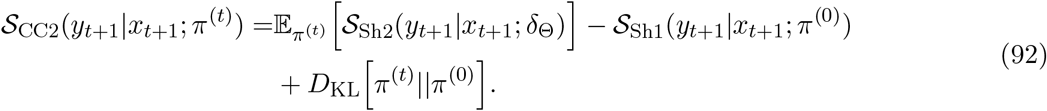

Now, we can replace 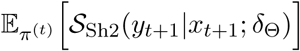 by using Eq. 79 and have

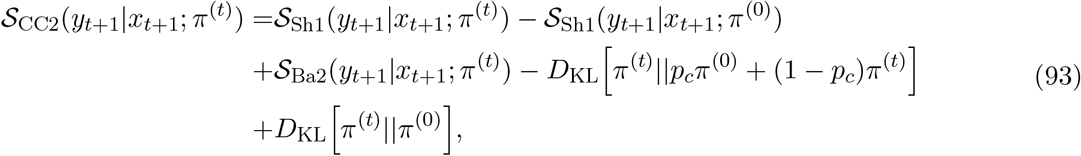

which is the same as Eq. 33. For the 1st definition of the Confidence Corrected surprise (c.f. Eq. 28), we can repeat all steps to have

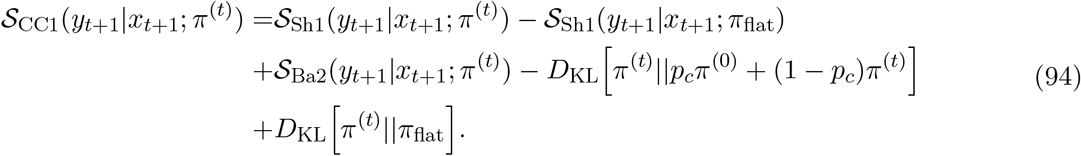

If *π*^(*t*)^ is absolutely continuous with respect to *π*_flat_, then we have *D*_KL_[π^(t)^ ‖ π_flat_] =*C* [π^(t)^] − *C* π_flat_], which completes the proof.■

#### Proof of Corollary 1

The corollary is the direct conclusion of Eq. 60 and Eq. 62. ■

#### Proof of Corollary 2

Let us show the set of possible observations by 𝒴. We assume that 𝒴 is bounded, i.e., |𝒴 | *<* ∞. By assumption, we have *P* (*y*_*t*+1_|*x*_*t*+1_; *π*^(0)^) = 1*/*|𝒴 |. We therefore (using Eq. 5, Eq. 9, and Eq. 10) have

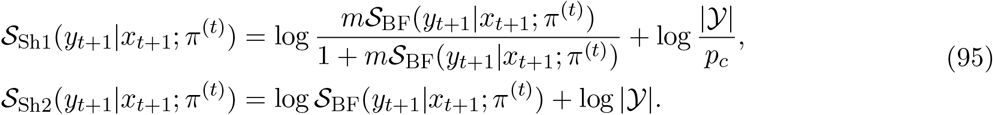

Both relations are invertible. Therefore, the proof is complete. ■

#### Proof of Corollary 3

In the limit of *p*_*c*_ → 1, we have 𝒮 _Sh1_(*y*_*t*+1_|*x*_*t*+1_; *π*^(*t*)^) = 𝒮 _Sh1_(*y*_*t*+1_|*x*_*t*+1_; *π*^(0)^) (c.f. Eq. 9) which implies that Δ𝒮 _Sh1_(*y*_*t*_ +1|*x*_*t*+1_; *π*^(*t*)^) (c.f. Proposition 2) i n Eq. 33 is equal to 0. Similarly, in the limit of *p*_*c*_ → 1, we have *D*_KL_[*π* ^*(t)*^||*p*_*c*_*π*^(0)^ + (1 – *p*_*c*_)*π*^*(t)*^) = *D*_KL_]*π* ^*(t)*^||π ^(0)^ Therefore, in the limit of *p*_*c*_ → 1 and given Eq. 33, we have 𝒮_CC2_(*y*_*t*+1_|*x*_*t*+1_; *π*^(*t*)^) = 𝒮_Ba2_(*y*_*t*+1_|*x*_*t*+1_; *π*^(*t*)^). ■

## Appendix C: Methods for case-studies

In this appendix, we provide methods for the analyses of section Different definitions make different predictions: Case studies in the main text.

### Methods for case-study 1

The formulas for surprise calculations in Fig. 5-Fig. 8 are given in this section. Note that we always have 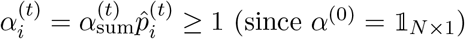; this bound determines axes’ boundaries in Fig. 5 and Fig. 6.

#### Surprise formulas

The Shannon surprise (c.f. Eq. 10) in the categorical is

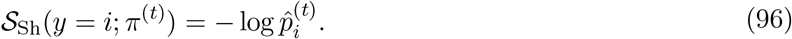

To compute the Bayesian and the Confidence Corrected surprise, we need to use the general formula for the KL-divergence between two Dirichlet distributions parameterized by *α* = [*α*_1_, …, *α*_*N*_] and *β* = [*β*_1_, …, *β*_*N*_] as (Gijsen et al., 2021)

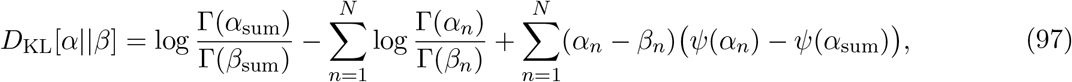

where we define 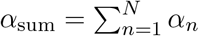and 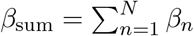, Γ is the gamma function, and *ψ* is the digamma function (Abramowitz & Stegun, 1948).

Using the general formula, the Bayesian surprise (c.f. Eq. 24) can be computed as

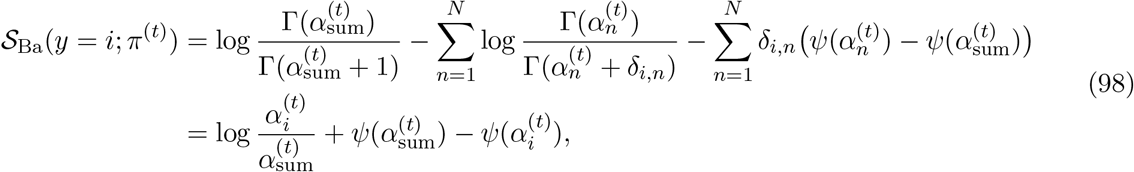

where we used 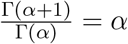 (Abramowitz & Stegun, 1948) to simplify the expression. We, therefore, have

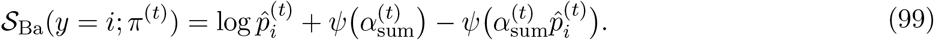

We can also rewrite Eq. 99 by using the following series representation of the digamma function for any *x >* 0 (Blagouchine, 2016)

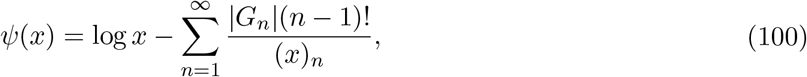

where *G*_*n*_ are ‘Gregory coefficients’ and (*x*)_*n*_ = *x*(*x* + 1)…(*x* + *n* − 1) is raising factorial. As a result, the Bayesian surprise is

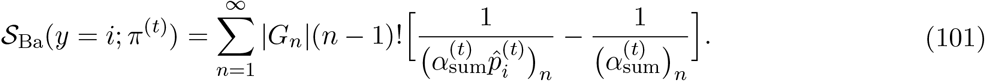

We use this representation in some of our proofs below.

The formula for the Confidence Corrected surprise can be written as

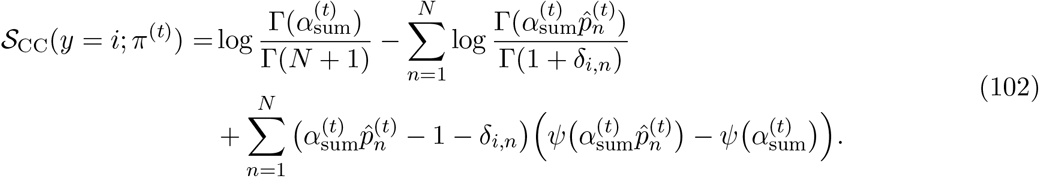

#### Influence of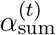

When 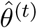 is fixed, then the Shannon surprise (Eq. 96) is constant with respect to 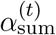. For the Bayesian surprise (Eq. 99), it is straightforward to show, using Eq. 101 and the fact that 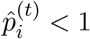, that

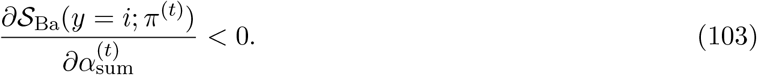

Therefore, the Bayesian surprise S_Ba_(*y* = *i*; *π*^(*t*)^) is always a decreasing function of 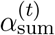. Moreover, using asymptotic (for 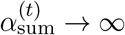) approximation of the digamma function (Abramowitz & Stegun, 1948), we have

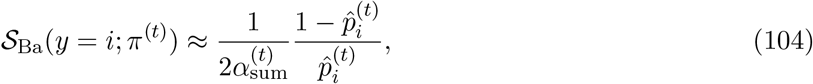

which also concludes that 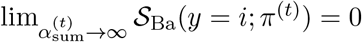.

We could not make any general statement about the relation of the Confidence Corrected surprise with 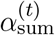 (c.f. Fig. 5 and Eq. 102). However, using asymptotic (for 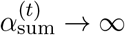) approximation of the gamma and digamma functions (Abramowitz & Stegun, 1948), we have

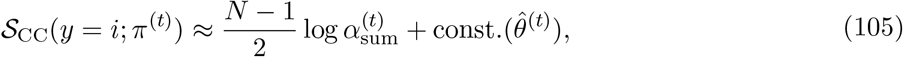

which concludes that 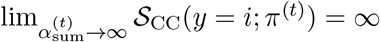.

#### Influence of 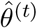

When 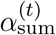 is fixed, then the Shannon surprise (Eq. 96) is decreasing with respect to 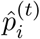 According to Eq. 101, the Bayesian surprise is also always a decreasing function of 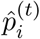 Moreover, both Shannon and Bayesian surprise are constant with respect to 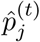 for any *j* /= *i*.

We could not make any general statement about the relation of the Confidence Corrected surprise with 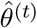 (c.f. Fig. 5, Fig. 6, and Eq. 102).

#### Simulation details

For Fig. 7A-C, for a given random seed, we sampled *Y*_*t*_ ∼ Cat({0.9, 0.1}) for 1 ≤ *t* ≤ 100. To model the temporal dynamics of surprise values, we use the common assumption (Gijsen et al., 2021; Maheu et al., 2019; Meyniel et al., 2016; Modirshanechi et al., 2019; Yu & Cohen, 2009) that participants use a simple leaky integration for the update of their beliefs, i.e.,

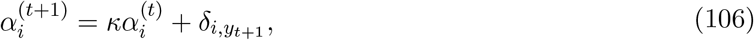

where *κ* ∈ [0, 1] is the leak parameter that determines how fast old observations are forgotten, and *δ* is the Kronecker delta function. With this update rule, we have the guarantee that 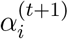 does not grow unlimited and participants are never too sure about their predictions (as long as *κ <* 1). To extract surprise values, we put 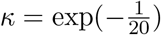 equivalent to an integration over a window of approximately 20 observations (Meyniel et al., 2016). However, note that our predictions are independent of the exact value of *κ*, and we found the same qualitative behavior of different surprise measures for 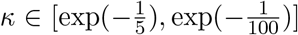 For Fig. 7B and Fig. 7C, we first averaged the surprise of *Y*_*t*_ = 1 (i.e., standard stimulus) and *Y*_*t*_ = 2 (i.e., deviant stimulus) in the early and the late phase of the task separately for each random seed. The values showed in the figure correspond to the mean and the standard error of the mean (over 40 different seeds) of the average-surprise of *Y*_*t*_ = 1 (Fig. 7B) and the average-surprise of *Y*_*t*_ = 2 (Fig. 7C).

For Fig. 8, for a given random seed, we sampled *Y*_*t*_ ∼ Cat({0.3, 0.35, 0.35}) for 1 ≤ *t* ≤ 150 and *Y*_*t*_ ∼ Cat({0.3, 0.1, 0.9}) for 151 ≤ *t* ≤ 300. To extract surprise values, we updated the belief at each time using leaky integration (c.f. Eq. 106) with 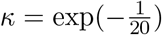. However, note that our predictions are independent of the exact value of *κ*, and we found the same qualitative behavior of different surprise measures for 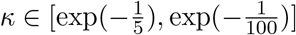. For Fig. 8C, we first averaged the surprise of *Y*_*t*_ = 1 in the averaging-period of phase 1 (76 ≤ *t* ≤ 150) and the averaging-period of phase 2 (226 ≤ *t* ≤ 300) separately for each random seed. The values showed in the figure correspond to the mean and the standard error of the mean (over 40 different seeds) of the average-surprise of *Y*_*t*_ = 1.

### Methods for case-study 2

The formulas for surprise calculations and the temporal update rule for the belief for data shown in Fig. 9 and Fig. 10 are given in this section.

#### Surprise formulas

Different measures of surprise for observing *y*_*t*+1_ = 1 given the cue *x*_*t*+1_ ∈ [0, 1] are calculated below. To compute the surprise of *y*_*t*+1_ = 2 given the cue *x*_*t*+1_ ∈ [0, 1], one only needs to replace all *x*_*t*+1_ by 1 − *x*_*t*+1_.

Using Eq. 42, the Bayes Factor surprise (c.f. Eq. 5) and the Shannon surprise (c.f. Eq. 10) can be written as

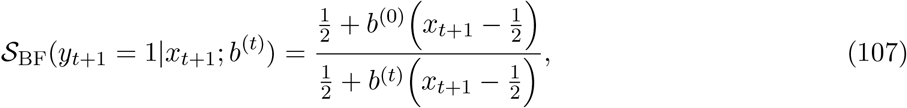

and

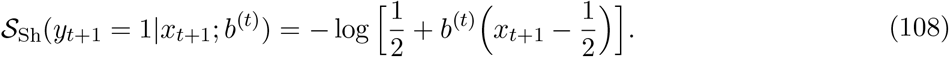

We define 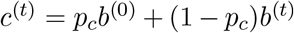 and 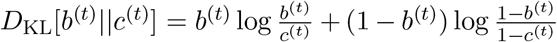. To compute the Bayesian surprise, we use the result of Lemma 1; to do so, we compute

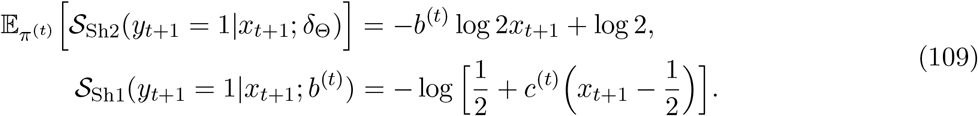

We, therefore, have

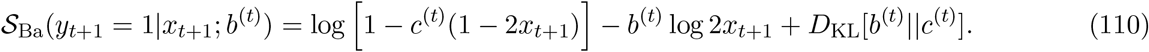

With similar tricks, the Confidence Corrected surprise can be computed as

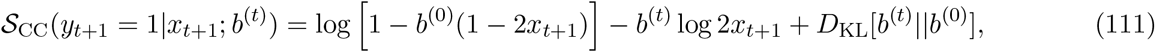

where we define 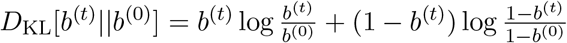.

#### The update rule

Using Proposition 1 for the update of the belief, we have

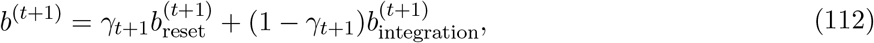

where *γ*_*t*+1_ is the adaptation rate as define in Proposition 1, and

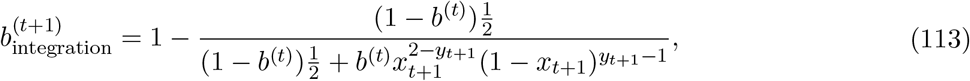

and similarly

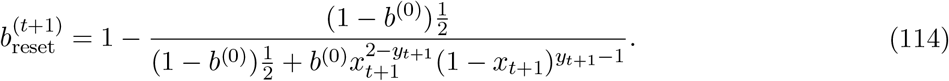

#### Simulation details

We fixed the parameters *θ*_1:150_ as shown in Fig. 10: *θ*_1:50_ = *θ*_101:150_ = 0 and *θ*_51:100_ = 1. Then, for each random seed, we randomly sampled the cue variables *x*_1:150_ independently: *X*_*t*_ ∼ Uniform({0.1, 0.9}), i.e., at each point, the oracle chooses one of the possible outcomes and assigns a probability of 0.9 to it. We then, given the same random seed, sampled the observations *y*_1:150_ as described before: *Y*_*t*_ ∼ Cat({0.5, 0.5}) whenever Θ_*t*_ = 0, and *Y*_*t*_ ∼ Cat({*x*_*t*_, 1 − *x*_*t*_}) whenever Θ_*t*_ = 1, where Cat stands for categorical distribution. Fig. 10A shows data generated for one random seed, and Fig. 10B shows the average belief and surprise over 500 random seeds.

### Methods for case-study 3

The formulas for the gain functions and the details of the update rules for Fig. 11 and Fig. 12 are given in this section.

To simplify the notation, we define, for all *p* ∈ (0, 1), *q* ∈ (0, 1), and *Y* ∈ {1, 2}

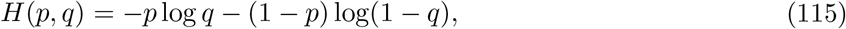

and

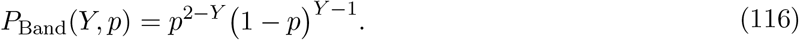

#### The belief and the update rule

Given the prior belief as in Eq. 44, the belief at time *t* is

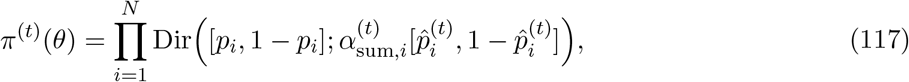

where

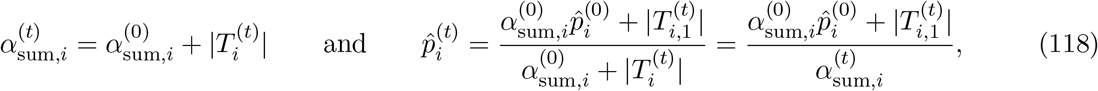

where 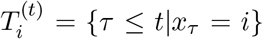 is the set of time points when arm *i* has been chosen, 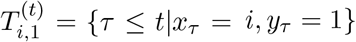 is the set of time points when arm *i* has been chosen and *y*_*τ*_ = 1 has been observed, and 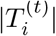 and 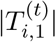 are the size of 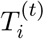 and 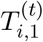, respectively. The marginal probability is

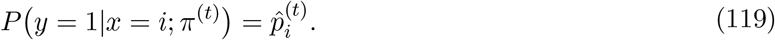

#### The optimal gain function

Using Eq. 117 and Eq. 118, we can re-write Eq. 46

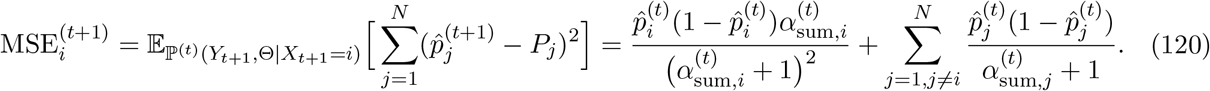

Therefore, the optimal strategy is

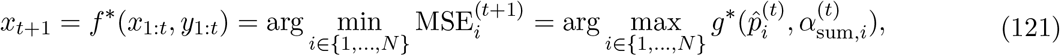

with the gain function

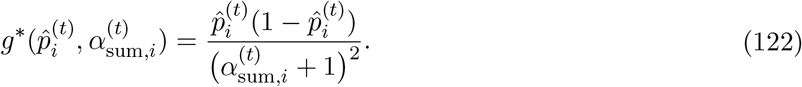

*g*^∗^ is independent of 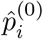 and 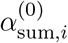, and its behavior with respect to the other two variables is as follows:

- With respect to 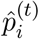 for any fixed 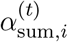, we have 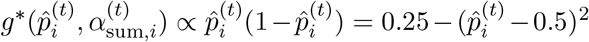 which has an inverted-U-shape, is an even function with respect to 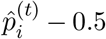, and has its maximum at 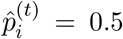. Moreover, consider a combination of 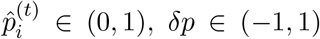 and 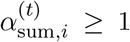 for which we have 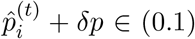 and 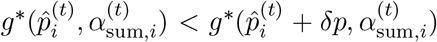. Then, there always exists a *δα >* 0 such that 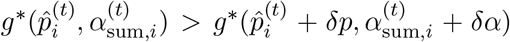. Therefore, independent of the difference in the stochasticity level (i.e. *δp*), as an arm gets to be chosen more often (as *δα* increases) it eventually becomes less informative.
- With respect to 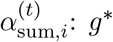 is always decreasing.

#### Gain function for the Bayes Factor surprise

For the Bayes Factor surprise, following Eq. 5 and Eq. 50, we have

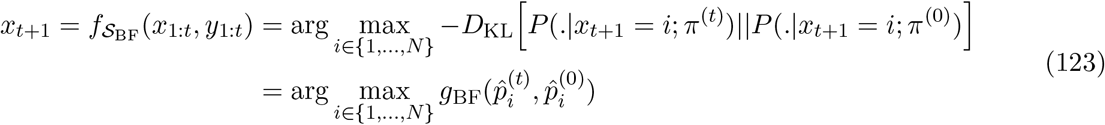

with the corresponding gain function (c.f. Eq. 115)

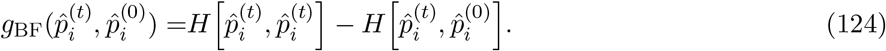

*g*_BF_ is independent of 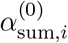 and 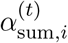, and its behavior with respect to the other two variables is as and follows:

- With respect to 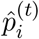: it has an inverted-U-shape with its maximum at 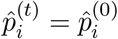.
- With respect to 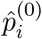: it has an inverted-U-shape with its maximum at 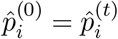.

#### Gain function for the Shannon surprise

For the Shannon surprise (c.f. Eq. 10) we have

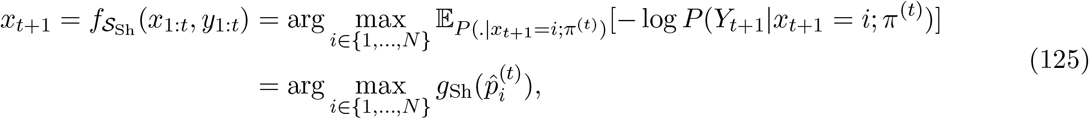

with the corresponding gain function (c.f. Eq. 115)

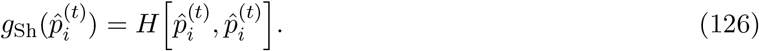

*g*_Sh_ is independent of 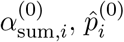,its behavior with respect to the other two variables is as follows: and 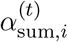, and it has an inverted-U-relation with 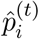 with its maximum at 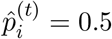.

#### Gain function for the Bayesian surprise

For the Bayesian surprise Eq. 24 we have (using Eq. 97, Eq. 99, and Eq. 116)

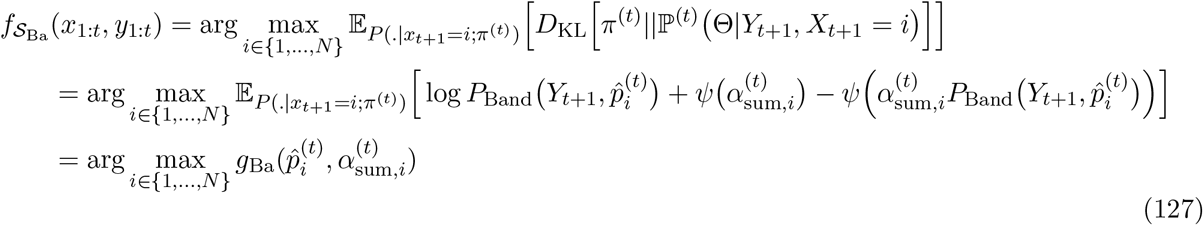

with the corresponding gain function

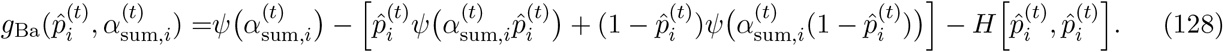

Using the expanded representation of 𝒮_Ba_ in Eq. 101, we can rewrite the gain function as

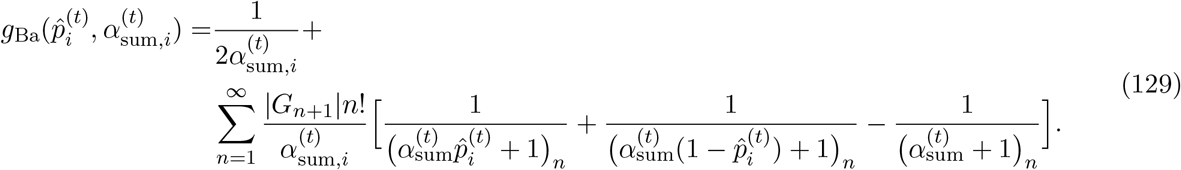

*g*_Ba_ is independent of 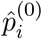 and 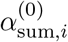, and its behavior with respect to the other two variables is as follows:

- With respect to 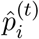: Using Eq. 129 and the fact that 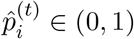, it is straightforward to show that the partial derivative of *g*_Ba_ with respect to 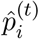 is negative for 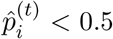, is positive for 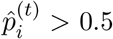, and is equal to 0 for 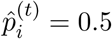. Therefore, *g*_Ba_ has a U-relation with respect to 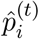 with its minimum at 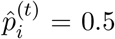. However, using asymptotic approximation of the digamma function (Abramowitz & Stegun, 1948), we have 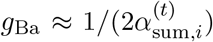 for large values of 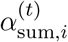. which is independent of 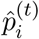. Therefore, seeking Bayesian surprise becomes independent of 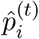 as 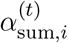 increases.
- With respect to 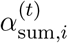: using the fact that 𝒮_Ba_ is always a decreasing function of 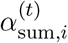 (c.f. Eq. 103) independent of the observation *Y*_*t*+1_ and its probability 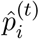, we can conclude that *g*_Ba_ is also a decreasing function of 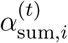.

#### Gain function for the Confidence Corrected surprise

For the Confidence Corrected surprise, by using Proposition 6, we have

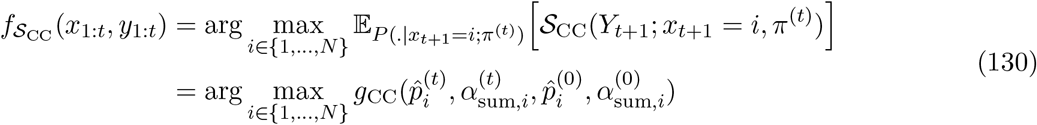

with the corresponding gain function

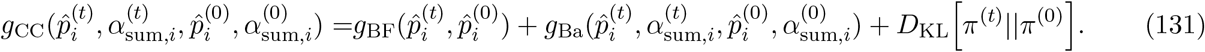

*g*_CC_ depends on all our 4 variables 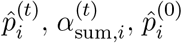, and 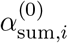. We could not make any general statement about the relation of *g*_CC_ with these variables. However, using asymptotic approximation of the gamma and digamma functions (Abramowitz & Stegun, 1948), we have

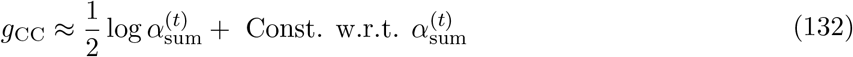

for large values of 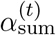. Therefore, *g*_CC_ is an increasing function of 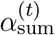 for large values of 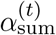.

#### Simulation details

We first put 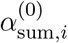 equal to 2 and 10 for scenarios 1 and 2, respectively. Then,for a given random seed, we randomly sampled prior parameter 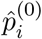 for *i* ∈ {1, …, 10}, differently for two scenarios: 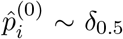 for scenario 1 and 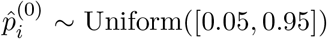 Uniform([0.05, 0.95]) for scenario 2. Given the same random seed, we build the environment by sampling 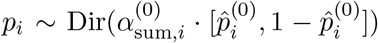 independently for *i* ∈ {1, …, 10} – note that *p*_*i*_s are not known to the agents. We then ran 6 different algorithms separately, corresponding to (1) *g*\*, (2) *g*_Sh_, (3) *g*_BF_, (4) *g*_Ba_, (5) *g*_CC_, and (6) *g*_ShR_. At time *t*, each algorithm computed its corresponding gain function *g* for all actions. Then, it chose the action with highest gain as *x*_*t*_ – when there was a tie, the action with the smaller index was chosen, e.g., action 2 was preferred to action 6. Given their actions *x*_*t*_, different algorithms observed different observations *y*_*t*_. Then, each algorithm updated its belief, and this procedure was repeated until *t* = 100. At time *t*, the mean-squared error 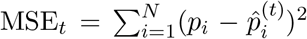 was computed as a measure of performance. Fig. 12A, Fig. 12C, and Fig. 13M-N show average MSE_*t*_ over 500 random seeds as a function of time.

#### Model recovery

To perform model-recovery, we characterized each exploration policy as a stochastic model with a softmax policy over a gain function: We assumed that the probability of taking action *x*_*t*+1_ = *i* by the agent that uses the gain function *g* is

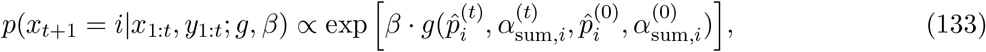

where *β* is a free parameter. Then, given a sequence of actions *x*_1:*t*_ and observations *y*_1:*t*_ (for one random seed) and for a gain function *g*, we approximated *p*(*x*_1:*t*_|*y*_1:*t*_; *g*) by max_*β*_ *p*(*x*_1:*t*_|*y*_1:*t*_; *g, β*). Because we have the same number of free parameters for all models, this approximation is equivalent to using Schwarz approximation – also known as Bayesian Information Criterion; BIC (Efron & Hastie, 2016; Neath & Cavanaugh, 2012; Schwarz, 1978). We then considered a uniform prior over policies and found the posterior probability of each policy given an observed sequence of action-choices: *p*(*g*|*x*_1:*t*_, *y*_1:*t*_) ∝ *p*(*x*_1:*t*_|*y*_1:*t*_; *g*). The confusion matrices in Fig. 12B and Fig. 12D show average *p*(*g*|*x*_1:*t*_, *y*_1:*t*_) over 500 random seeds.

## Appendix D: EEG analysis

In this appendix, we provide the methods and results for the EEG analysis reported in Fig. 7.

### Dataset

We used a publicly available dataset published in Robbins et al., 2018. The dataset has been originally collected for the study reported in Hairston et al., 2014 and includes the behavioral and the EEG data of 18 adult participants in a visual oddball experiment. The details of the experiment are provided in Robbins et al., 2018; we briefly review the main features here.

Participants were instructed to respond to the standard and deviant stimuli separately by pressing different buttons. The standard stimulus was an image of a US soldier, and the deviant stimulus was an image of an enemy combatant. At each trial, one of the two images was randomly chosen (with roughly 1/8 probability of choosing the deviant image (Modirshanechi et al., 2019)) and presented for 150ms on a computer screen. Each participant completed three blocks of (approximately) 89 trials each. EEG signals were sampled at a rate of 512Hz in a 64-channel electrode space. The data was preprocessed to remove line noise, detect and interpolate bad electrodes, and remove artifacts (see Robbins et al., 2018 for details). We used the preprocessed data.

### Further pre-processing

For each participant, we averaged the EEG signals over the central electrodes (Cz, C1, and C2) which we then further smoothed (moving averaging with the window of length 50ms) and standardized (zero mean and unit variance). We extracted EEG trials from 100ms before to 900ms after the stimulus onset and removed the baseline activity by subtracting the mean calculated over the first 100ms. We excluded error trials (i.e., the trials where participants either pressed the wrong button or did not press any button, error rate: 3.8% ± 2.4%, range: 0.4% to 11.3%) from further analyses.

### Analysis

For each participant, we computed the event related potentials (ERPs) separately for the standard and the deviant stimuli, averaged over all trials. Fig. 7D shows the mean and the standard error of the mean (over participants) of the standard and the deviant ERPs. We used one-sample t-test (FDR controlled by 0.1 (Benjamini & Hochberg, 1995)) and found time windows where the standard and the deviant ERPs were significantly different. Then, for participant *i* ∈ {1, …, 18}, we found the time-points where the deviant ERP had its maximum in time windows of 450-600ms (which we call *T*_*i*_, corresponding to *Z* in Fig. 7D) and 600-800ms (which we call 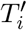, corresponding to *Z*′ in Fig. 7D). Then, for participant *i* and trial *t*, we averaged the ERP amplitudes at time windows of *T*_*i*_ ± 20ms (called *z*_*i,t*_) and 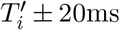 (called 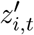) – see Kolossa et al., 2015; Kopp and Lange, 2013 for similar approaches. Then, we averaged the *z*_*i,t*_ and 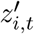 of the standard and the deviant stimuli in the early and the late phase of the task separately for each participant. The values showed in Fig. 7E and Fig. 7F corresponds to the mean and the standard error of the mean (over 18 participants) of the average-*Z* (or the average-*Z*′) at standard trials (Fig. 7E) and the average-*Z* (or the average-*Z*′) at deviant trials (Fig. 7F).

### Results

*Z* is a candidate for the amplitude of the EEG P3a component and *Z*′ for the amplitude of the EEG P3b component (Kolossa et al., 2015; Kopp & Lange, 2013; Visalli et al., 2021). It has been reported that the latency of the EEG P300 component varies depending on task features (Kutas et al., 1977; Magliero et al., 1984), but we also note that the latency of *Z* and *Z*′ is different from the latency of P3a and P3b in previous surprise studies (Kolossa et al., 2015; Kopp & Lange, 2013; Visalli et al., 2021). Therefore, we consider our results complementary to the earlier results.

Our results show that *Z* has a behavior consistent with the Shannon surprise. In contrast to previous studies that found P3a (in tasks different from oddball task) to be a signature of the Bayesian surprise (Kolossa et al., 2015; Visalli et al., 2021), our results provide strong evidence against a correlation between *Z* and the Bayesian or the Confidence Corrected surprise (Fig. 7E and Fig. 7F). Our results show that the prediction of the Bayesian surprise is not consistent with the behavior of *Z*′ either (Fig. 7F). If *Z*′ is interpreted as the P3b amplitude, then this result supports earlier findings that P3b reflects the Shannon surprise (Kolossa et al., 2015; Kopp & Lange, 2013; Visalli et al., 2021). However, our results cannot determine whether the Shannon or the Confidence Corrected surprise is a better fit for *Z*′ and suggest a mixed effect (e.g., a correlation between *Z*′ and a linear combination of the Shannon or the Confidence Corrected surprise) which can be tested by further and more advanced analyses.

## Appendix E: Regularized Shannon surprise

In this Appendix, we consider *R*(*z*) = *z* and derive the formulas used in Fig. 13.

### Case-study 1

Using Eq. 96, we can write

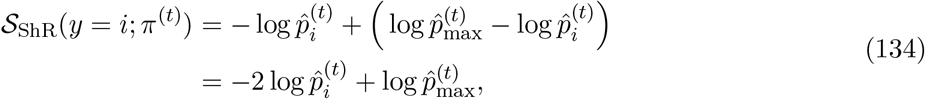

where

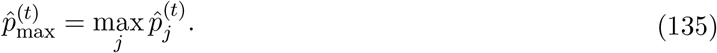

As a result, we have

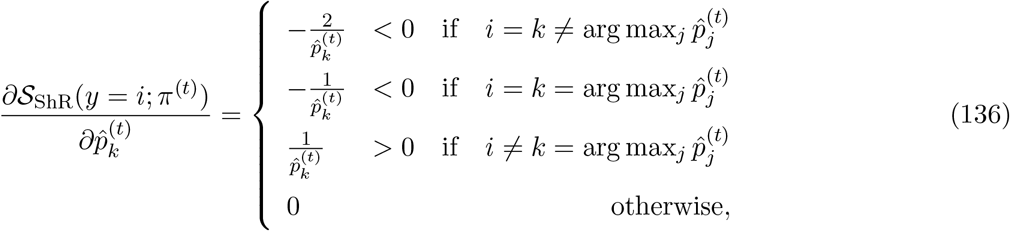

The regularized Shannon surprise is constant with respect to 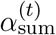.

### Case-study 2

We define

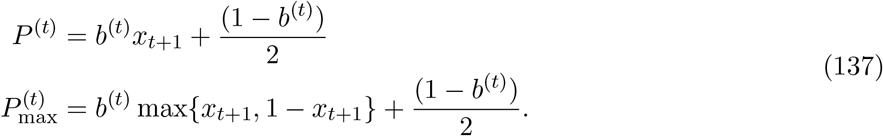

Using Eq. 108, we have

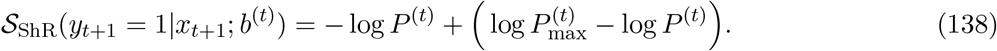

### Case-study 3

Using the general form of the surprise seeking strategies (c.f. Eq. 49) and the definition of the regularized Shannon surprise (c.f. Eq. 53) we can write

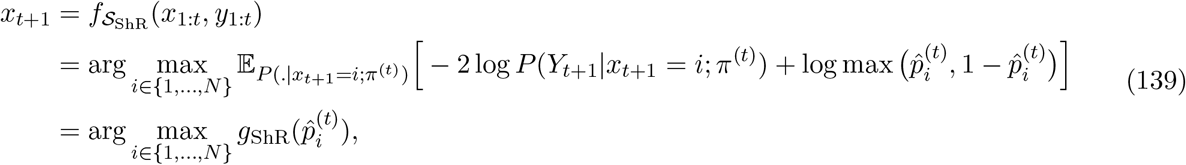

with the corresponding gain function

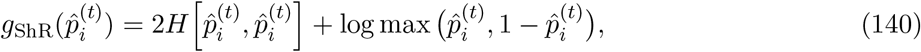

which is an even function of 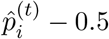. For 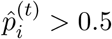, the derivative of the gain function is

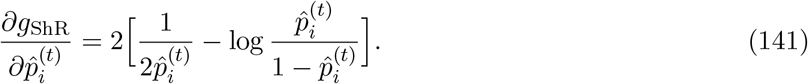

The derivative is positive for 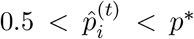, is negative for 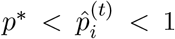, and is 0 for 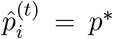, where *p*^∗^ ≈0.68. Therefore, the gain function has an inverted-W-relation with 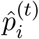 with its maximum at 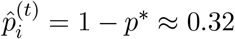 and 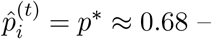 corresponding to the most preferred arms.

## Footnotes

1 For example, assume that 𝒮 = *f* (𝒮′) for an invertible function *f*. If an estimator of the variable *Z* is found using the measure 𝒮 as 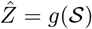, then we can rewrite the same estimator in terms of 𝒮′ as 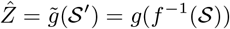. Because *g*(𝒮) and 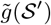 essentially have the same explanatory power given any function *g* and any measure of performance, the two surprise measures 𝒮 and 𝒮′ are equally informative about the variable *Z* in this regard. However, this is not necessarily true if one restricts the estimators to a particular class of functions – e.g., if the estimators are constrained to be linear with respect to surprise measures while *f* is nonlinear. Such limitations can be avoided by using non-parametric statistical methods like Spearman or Kendall correlations (Corder & Foreman, 2014).

2 If the initial belief *π*^(0)^ is a Dirichlet distribution, then *π*^(*t*)^ is also a Dirichlet distribution at any time *t* for a wide range of learning rules (Faraji et al., 2018; Liakoni et al., 2021; Maheu et al., 2019; Markovic et al., 2021; Meyniel et al., 2016; Modirshanechi et al., 2019; Ryali et al., 2018; Yu & Cohen, 2009) – since the Dirichlet distribution is the conjugate prior of the categorical distribution (Efron & Hastie, 2016).

3 We note that the qualitative behavior of confidence in Fig. 5 and Fig. 6 is the same for both definitions *C*[*π*^(*t*)^] and CatConf(*t*).

4 Note that *p*_*i*_ is the probability of observing *Y*_*t*_ = 1 for the arm *X*_*t*_ = *i*, whereas in the the first case-study, *p*_*i*_ was the probability of observing *Y*_*t*_ = *i* and there was no cue variable *X*_*t*_. Similarly, 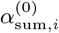 is an *arm-specific* belief parameter, in contrast to the general belief parameter 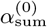 in the the first case-study.

